# Three-dimensional structured illumination microscopy with enhanced axial resolution

**DOI:** 10.1101/2022.07.20.500834

**Authors:** Xuesong Li, Yicong Wu, Yijun Su, Ivan Rey-Suarez, Claudia Matthaeus, Taylor B. Updegrove, Zhuang Wei, Lixia Zhang, Hideki Sasaki, Yue Li, Min Guo, John P. Giannini, Harshad D. Vishwasrao, Jiji Chen, Shih-Jong J. Lee, Lin Shao, Huafeng Liu, Kumaran S. Ramamurthi, Justin W. Taraska, Arpita Upadhyaya, Patrick La Riviere, Hari Shroff

## Abstract

We present two distinct, complementary methods for improving axial resolution in three-dimensional structured illumination microscopy (3D SIM) with minimal or no modification to the optical system. First, we show that placing a mirror directly opposite the sample enables 4-beam interference with higher spatial frequency content than 3D SIM illumination, offering near-isotropic imaging with ∼120 nm lateral and 160 nm axial resolution. Second, we develop an improved deep learning method that can be directly applied to 3D SIM data, obviating the need for additional hardware. This procedure results in ∼120 nm isotropic resolution and can be combined with denoising to facilitate volumetric imaging spanning dozens of time points. We demonstrate the potential of these advances by imaging a variety of cellular samples, delineating the nanoscale distribution of vimentin and microtubule filaments, observing the relative positions of caveolar coat proteins and lysosomal markers, and visualizing rich cytoskeletal dynamics within T-cells in the early stages of immune synapse formation.

## Introduction

Three-dimensional structured illumination microscopy (3D SIM^1^) excites the sample with nonuniform illumination, providing additional information outside the diffraction-limited passband that is encoded in the fluorescence captured by a series of diffraction-limited images. Decoding this extra information mathematically yields a super-resolution reconstruction with approximately doubled resolution compared to widefield microscopy in lateral and axial dimensions. Although this gain is more modest than that offered by other methods^2^, in thin samples 3D SIM offers other advantages including superb optical sectioning, relatively low illumination dose and high acquisition speed (enabling ‘4D’ volumetric time-lapse imaging in living cells^3,4^), and compatibility with arbitrary fluorophores (facilitating multicolor super-resolution imaging^5^). This unique combination of attributes has provided insight into bacterial cell division^6-8^; DNA repair^9^ and replication^10^; inter-parasite communication^11^ and host-parasite interactions^12^; gene regulation^13,14^; the immune synapse^15^; and nuclear^5^, chromosomal^16^, membrane^17^, centriole^18^, and mitochondrial^19^ architecture.

Although the axial resolution offered by 3D SIM is markedly better than that of widefield microscopy, it is still limited to ∼300 nm using high numerical aperture (NA) objectives. This value is slightly worse than the lateral resolution limit of ∼250 nm in widefield microscopy, and considerably worse than the ∼120 nm lateral resolution of 3D SIM. Thus, 3D SIM reconstructions are anisotropic, distorting and obscuring fine features along the axial dimension. Reducing or eliminating the anisotropy is desirable, yet relatively few solutions exist^20,21^, and none have been widely adopted.

Here we demonstrate two complementary methods for markedly improving axial resolution in 3D SIM, with minimal or no modification to the 3D SIM optical path. First, we show that placing a mirror directly opposite the sample enables 4-beam interference, producing higher axial spatial frequency components in the illumination pattern and corresponding improvements in resolution. Using this method, we obtain an axial resolution of ∼160 nm, much closer to the ∼120 nm lateral resolution of 3D SIM and thus producing nearly isotropic reconstructions. Second, we develop improved deep learning algorithms that operate on 3D SIM data (i.e., without the mirror), producing reconstructions with isotropic ∼120 nm spatial resolution. This computational method may be further combined with denoising algorithms, enabling high-quality, isotropic 4D super-resolution imaging over dozens of volumes. We showcase the power of these methods on a plethora of fixed and live cellular samples. In vegetative and sporulating bacteria, we visualized membranes, cell division proteins, and core components of the spore coat. In eukaryotic cells, we inspected membrane-encased actin filaments and pores that traversed thin membrane extensions; delineated the nanoscale positioning of vimentin and microtubules; observed the relative spatial distribution of caveolar coat proteins and performed time-lapse volumetric imaging of mitochondrial, lysosomal, and cytoskeletal dynamics.

## Results

### Four-beam interference for higher axial resolution

Spatial resolution and optical sectioning in 3D SIM are determined by the convolution of the structured illumination pattern’s spatial frequency components with the widefield detection optical transfer function (OTF, **Supplementary Fig. 1a, b**)^1^. Anisotropic spatial resolution is a direct consequence of limited angular aperture on both the illumination and detection side: (1) the three wave vectors whose interference produces the structured illumination pattern lie on a spherical cap with limited angular extent; (2) the limited angular range over which fluorescence is collected produces an OTF with greater lateral than axial extent. We thus considered strategies to increase angular aperture to improve axial resolution in 3D SIM.

On the illumination side, if the angle between mutually coherent illumination beams is increased beyond the limit imposed by the numerical aperture (NA) of the objective lens – either by using a mirror^22^ or a second objective lens^23^ – the resulting interference pattern may contain higher axial illumination frequency components up to a limit of 2n/*γ* for counter-propagating beams, where n is the index of refraction and *γ* the wavelength of illumination. Such ‘standing wave microscopes’^23^ were demonstrated decades ago and – for very thin samples – indeed unveil axial detail obscured in 3D SIM. For thicker samples, however, gaps in the OTF support (**Supplementary Fig. 1c**) preclude optical sectioning and generate severe axial ‘ringing’ in the resulting images.

These problems are resolved in I^5^S^20^, whereby two diametrically opposed objectives introduce six mutually coherent illumination wavevectors whose interference yields 19 illumination frequency components instead of the seven in 3D SIM. The same objectives are used to collect fluorescence, which is also coherently interfered. The combination of coherent illumination and coherent detection (**Supplementary Fig. 1d**) produces reconstructions with an impressive ∼100 nm isotropic spatial resolution. However, I^5^S also presents severe practical challenges. First, the two paths for illumination and fluorescence require more optics than traditional 3D SIM, adding cost and complexity and diminishing sensitivity. Second, and more importantly, due to the short coherence length of fluorescence, the emission paths must be carefully aligned to near zero path length difference, and the alignment maintained to much better than one wavelength. In practice this requires active feedback of multiple optical elements, further adding to instrument complexity. Third, refractive index (RI) mismatch between sample and immersion fluid introduces aberrations. We suspect these reasons might explain why the only demonstration of I^5^S on biological samples^20^ was limited to relatively small fields of view (minimizing spatially-varying phase aberrations), fixed cells with special mounting protocols (designed to minimize RI mismatch), and single color applications (since each additional color would likely require re-adjustment of the optical path).

An intriguing alternative to I^5^S was recently proposed^21^, whereby the central illumination beam in 3D SIM is isolated with a low-NA objective opposite the sample, re-imaged to a mirror, and reflected back to the sample. In this manner, a fourth, reflected beam interferes with the original 3-beams, yielding a 4-beam interference pattern with finer axial structure than the illumination pattern in conventional single objective 3D SIM (**Supplementary Fig. 1e**). If a high NA objective lens is used to collect the fluorescence emission, the theoretically predicted axial resolution is still substantially better than 3D SIM, albeit worse than I^5^S. The main advantage of this method is the relatively simple optical design compared to I^5^S.

Even with this proposal, notable challenges exist. First, while simpler than in I^5^S, the optics necessary to reflect the central beam still require stable alignment and add complexity relative to single objective 3D SIM (e.g., two objectives are still required). Second, the reflected beam must traverse multiple optical elements, adding undesirable wavefront distortion. Third, such distortion will also be introduced by differences between the refractive index of air (the medium in which the additional optical elements are placed) and buffer (in which the sample is placed). Fourth, and perhaps most importantly, the additional optical path length required would likely span almost a meter. This implies that the illumination source must have a coherence length of at least this length, so that interference between direct and reflected beams is possible. This requirement rules out common single-mode diode laser sources often used in microscopy.

We reasoned that placing a mirror directly opposite the sample by immersing it in the sample fluid would facilitate 4-beam SIM (**Fig. 1a-c**), offering advantages over the previously proposed design^21^: (i) the mirror can be placed close (easily within 100 μm) to the sample, enabling the use of commonly available, short coherence length diode lasers; (ii) the short optical path length from coverslip to mirror and back to coverslip implies that interferometric stability only needs to be maintained over this length scale; (iii) aberrations due to index mismatch can be minimized.

**Fig. 1.**
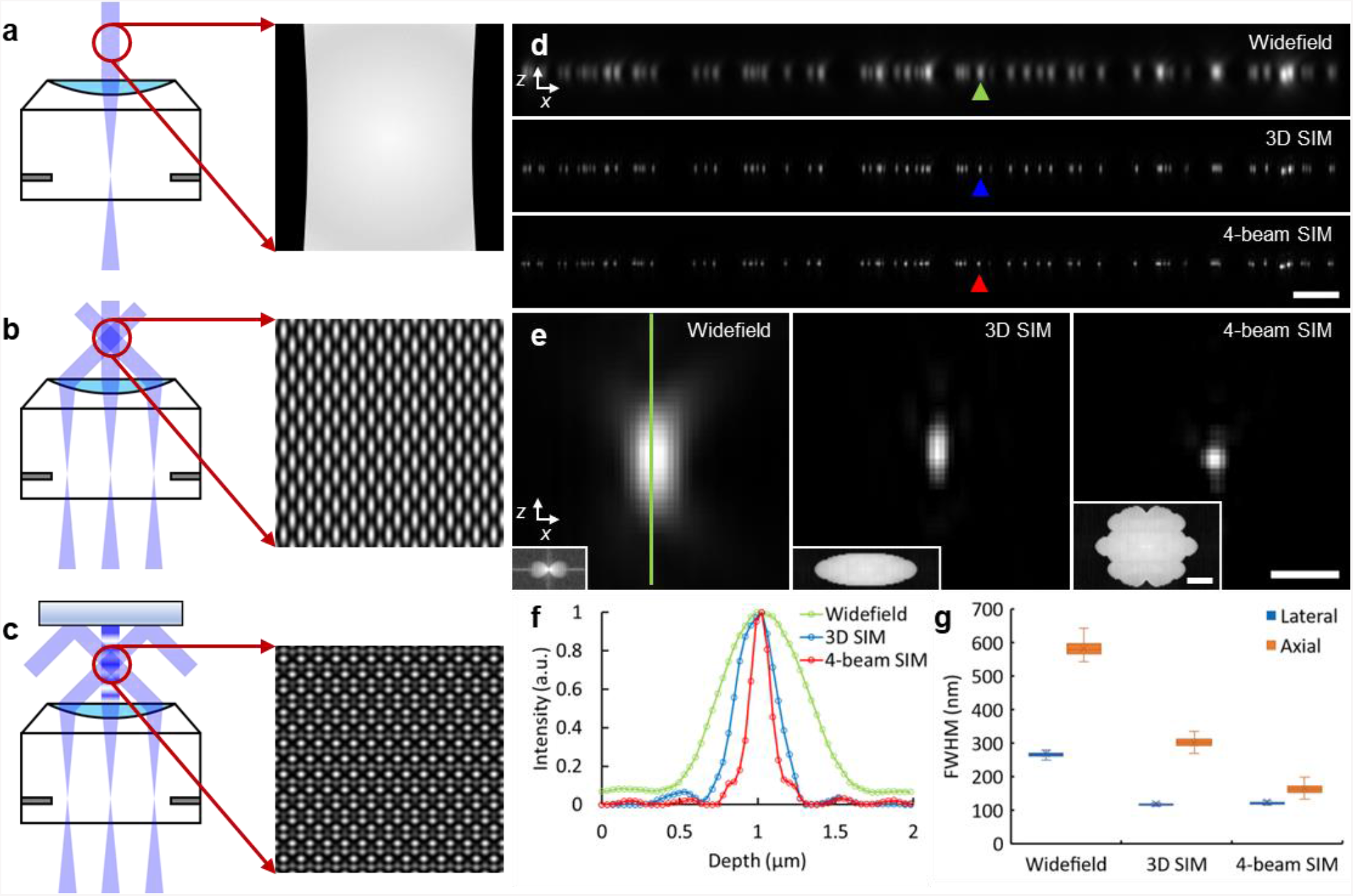
Improving axial resolution in three-dimensional structured illumination microscopy. **a-c**) Schematic representations of beam illumination at objective back focal plane and sample planes for widefield microscopy (single beam illumination, **a**)), 3D SIM (three beam illumination, **b**)), and 4-beam SIM (a mirror opposite the sample is used to back reflect the central beam, producing four beam interference, **c**)). Higher magnification illumination views at right show fine axial structure in 4-beam SIM pattern, absent in 3D SIM or widefield microscopy. **d**) Axial cross-sectional views of 100 nm beads, as imaged in widefield microscopy (top), 3D SIM (middle), and 4-beam SIM (bottom). **e**) Higher magnification views of bead highlighted by colored arrowheads in **d**), illustrating progressive improvement in axial resolution. Insets show magnitude of optical transfer functions (k_x_/k_z_ plane) derived from images. **f**) Line profiles corresponding to bead images shown in **e**), taken along vertical green line in **e**). **g**) Quantification of lateral (blue) and axial (orange) full width at half maximum (FWHM) for *N* = 102, 100, 99 beads for widefield microscopy, 3D SIM, and 4-beam SIM, respectively. See also **Supplementary Table 1**. Scale bars: 2 μm **d**) and 500 nm **e**); 1/200 nm^-1^ for Fourier transform insets in **e**).

To demonstrate this method, first we simulated illumination and detection configurations for commercially available objectives, finding that either a 1.35 NA silicone oil immersion or a 1.27 NA water immersion objective support 4-beam SIM imaging (**Supplementary Fig. 2**). Second, we constructed a 3D SIM system that served as the base for our method (**Supplementary Figs. 3-7**), using established protocols^24,25^ to confirm the quality of our illumination pattern and raw data (**Supplementary Fig. 8**). Third, we mounted and immersed a piezoelectrically controlled mirror directly over the sample, enabling 4-beam SIM (**Supplementary Fig. 9, Supplementary Video 1**).

We initially imaged 100 nm yellow-green beads using the 1.35 NA objective to characterize our 4-beam SIM (**Fig. 1d, e**). Using 45.6% iodixanol^26^ to match the RI of the silicone oil, thereby minimizing spherical aberration^27^ and focal shift^28^, we collected 15 images (5 phases per orientation, 3 orientations) per plane and reconstructed image stacks similarly to conventional 3D SIM (**Methods**). As expected, 4-beam SIM maintained the ∼2-fold lateral resolution enhancement of 3D SIM over widefield microscopy (4-beam: 124 +/- 12 nm full width half maximum (FWHM), 3D SIM: 119 +/- 11 nm, widefield: 268 +/- 16 nm, *N* = 99, 100, 102 measurements, respectively) while offering ∼2-fold better axial resolution than 3D SIM (4 beam: 163 +/- 13 nm, 3D SIM: 301 +/- 13 nm, widefield: 581 +/- 23 nm, **Fig. 1f, g, Supplementary Table 1**). We obtained similar results using the 1.27 NA water lens (**Supplementary Fig. 10, Supplementary Table 1**). Bead imaging also highlighted the importance of keeping the illumination pattern maxima centered on the detection focal plane, as even a ∼40 nm relative shift resulted in discernible axial ringing in the reconstructions (**Supplementary Fig. 11a**). To address this issue, we developed a bead-based feedback scheme that kept both focal plane and mirror stable to within 10-20 nm (**Supplementary Figs. 11b, c, 12, Methods**).

### Near-isotropic super-resolution imaging in biological samples

We next investigated the applicability of 4-beam SIM to biological samples, first using the 1.35 NA silicone oil lens on iodixanol index-matched samples (**Fig. 2**). On ∼1 μm thick, live, vegetative *B. subtilis* cells stained with CellBrite Fix 488, 4-beam SIM provided crisp lateral (**Fig. 2a**) and axial (**Fig. 2b, Supplementary Video 2**) views of cell membranes. Widefield microscopy barely resolved the upper and lower cell membranes in axial views, which were better visualized in 3D SIM. However, 4-beam SIM provided even more distinct images of membranes (**Fig. 2b**), as seen visually and from quantification of the apparent membrane thickness (∼160-175 nm, similar to the axial resolution estimates on beads, **Fig. 1g**). In some cases, we observed fine membrane invaginations that appeared indistinct or badly blurred in 3D SIM (**Fig. 2c, d**), underscoring the ability of 4-beam SIM to unveil axial detail otherwise masked by diffraction.

**Fig. 2.**
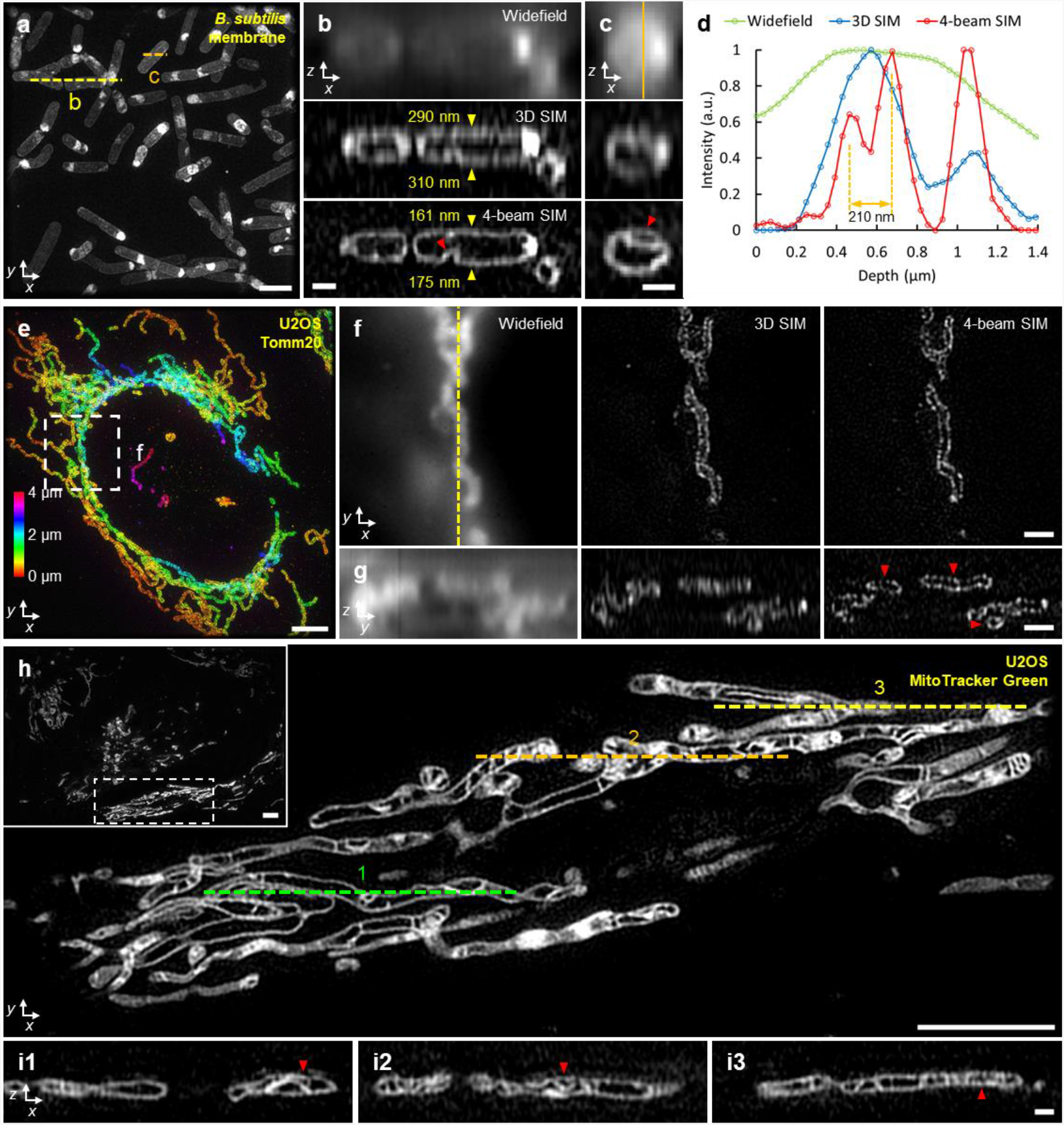
4-beam SIM enables near-isotropic imaging of biological samples. **a**) Maximum intensity projection of live vegetative *B. subtilis* stained with CellBrite Fix 488, marking membranes, imaged in 4-beam SIM. **b, c**) Axial views along yellow **b**) and orange **c**) dotted lines in **a**), comparing widefield microscopy (top), 3D SIM (middle), and 4-beam SIM (bottom). Yellow arrowheads in **b**) highlight upper and lower cell membranes; red arrowheads highlight membrane invaginations. See also **Supplementary Video 2. d**) Line profiles corresponding to vertical orange line in **c**). **e**) Maximum intensity projection of fixed U2OS cell labeled with Tomm20 primary and rabbit-AlexaFluor 488 secondary antibodies, marking outer mitochondrial membrane. Image is depth coded as indicated. Higher magnification lateral views (single planes) **f**) corresponding to white dashed rectangle in **e**) are shown, comparing widefield microscopy (left), 3D SIM (middle), and 4-beam SIM (right) are indicated, in addition to corresponding axial views **g**) taken across vertical yellow dashed line in **f**). Red arrowheads highlight void regions obscured in 3D SIM and widefield microscopy. See also **Supplementary Video 3. h**) Overview (inset) and higher magnification view of single lateral plane of mitochondria labeled with MitoTracker Green FM in live U2OS cell, highlighting inner mitochondrial substructure. **i**) Axial cross sections taken along green, orange, and yellow dashed lines in **h**) highlight fine substructure within mitochondria (red arrowheads). See also **Supplementary Video 4**. All data were acquired with 1.35 NA silicone immersion objective, with samples index-matched in 45.6% iodixanol. Scale bars: 2 μm **a**); 500 nm **b, c, i**); 4 μm **e, h**); 1 μm **f, g**).

Next, we examined thicker, eukaryotic samples over imaging fields spanning tens of microns in each lateral dimension (**Fig. 2e-i**). When performing widefield imaging of fixed U2OS cells immunolabeled for Tomm20 (**Fig. 2e**), marking the outer mitochondrial membrane, diffraction obscured the inner mitochondrial space. By contrast, the superior resolving power of 3D SIM and 4-beam SIM produced lateral views where such regions void of label were easily discerned (**Fig. 2f**). In axial views, however, only 4-beam SIM could reliably resolve the void regions, as the relatively poorer axial resolution of widefield or 3D SIM artificially ‘filled in’ and distorted the inner mitochondrial space (**Fig. 2g, Supplementary Video 3**). In a second example, we stained living U2OS cells with MitoTracker Green FM, marking the internal mitochondrial space (**Fig. 2h-i, Supplementary Video 4**). In contrast to 3D SIM^3^, 4-beam SIM enabled visualization of fine mitochondrial substructure in axial (**Fig. 2i**) as well as lateral views.

During experiments with the 1.35 NA silicone oil lens, we noticed that imaging samples immersed in the RI-matched iodixanol solution displayed substantially greater bleaching than samples immersed in PBS (**Supplementary Fig. 13**). This fact, combined with the unknown effect of iodixanol on living samples at 45.6% composition and the awkward sample handling associated with such a viscous buffer, prompted us to use the 1.27 NA water lens for subsequent experiments.

Using this lens, we first verified that the resolution enhancement obtained on beads (**Supplementary Fig. 10**) extended to biological samples by imaging live, vegetative *B. subtilis* labeled with DivIVA-GFP (**Supplementary Fig. 14a**). DivIVA is known to assemble at nascent division sites^29^, and is thought to spatially regulate cell division. Like earlier 3D SIM measurements, we resolved the ‘double-ring’ DivIVA arrangement at the site of active division with 4-beam SIM. However, the near-isotropic resolution of our method also provided clearer axial views of the circularly shaped rings (**Supplementary Fig. 14b, Supplementary Video 5**), which otherwise assumed a distorted ovoid appearance in 3D SIM measurements^29^.

We next extended our method for dual-color imaging (**Fig. 3**). Illuminating the sample with distinct wavelengths (e.g., 488 nm and 561 nm) produces different axial offsets between the illumination pattern maxima and detection focal plane. Ideally this offset would be minimized for each illumination wavelength^24^, thereby maximizing pattern contrast, optical sectioning, and axial resolution. For 4-beam SIM, minimizing this offset proved mandatory to minimize axial ringing in the reconstructions (i.e, ‘gaps’ in the OTF, **Supplementary Fig. 15**). We thus determined the optimal position of our camera for each illumination wavelength, translating it when switching from one color channel to the next (**Methods**).

**Fig. 3.**
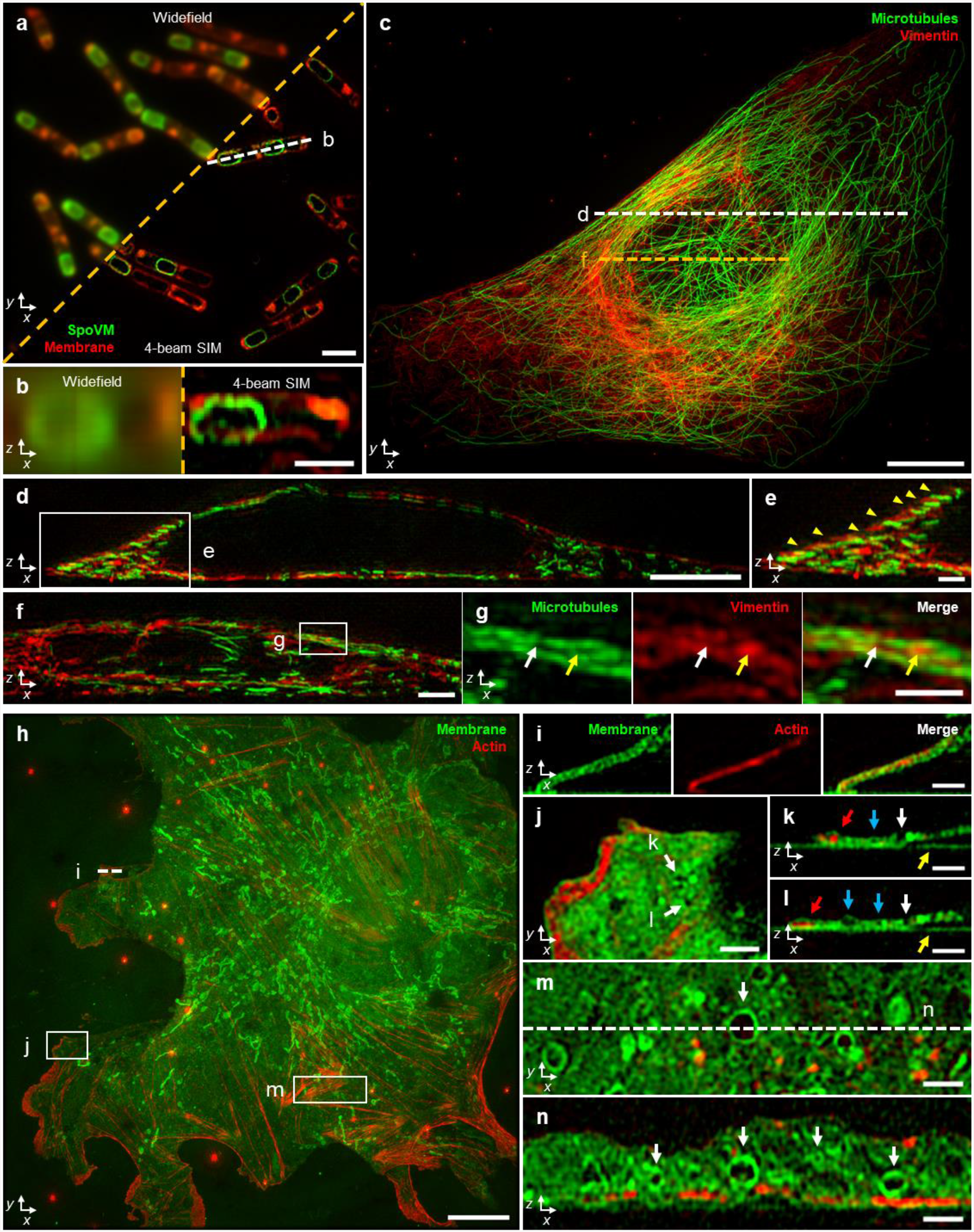
Near-isotropic imaging in two colors via 4-beam SIM. **a**) Single lateral plane of live, sporulating *B. subtilis* with SpoVM-GFP label, marking spores (green); and CellBrite Fix 555 label, marking membrane (red). **b**) Axial view (single plane) along white dotted line in **a)**. Images to the left of orange dashed line in **a, b**) show widefield images for comparison. **c)** Maximum intensity projection of fixed U2OS cell with Alexa Fluor 488 immunolabeled microtubules (green) and Alexa Fluor 594 immunolabeled labeled vimentin (red). **d**) Axial view corresponding to white dashed line in **c**). **e**) Higher magnification view of white rectangular region in **d**), indicating apparent alternating stratification of microtubules and vimentin filaments (yellow arrowheads). Image is a maximum intensity projection over 10 planes. **f**) Axial view corresponding to dashed orange line in **c)**. Image is a maximum intensity projection over 5 planes. **g**) Higher magnification view of white rectangular region in **f**), highlighting apparent ‘filling’ of inter-microtubule gaps by vimentin (arrows). Microtubule (left), vimentin (middle), and merged (right) images are shown. **h**) Maximum intensity projection image of fixed mouse liver sinusoidal endothelial cell with CellBrite Fix 488 label, marking membrane (green) and Alexa Fluor 568 Phalloidin, marking actin filaments (red). See also **Supplementary Video 6. i**) Axial view corresponding to white dashed line in **h**), highlighting membrane signal encapsulating actin. Image is a maximum intensity projection over 3 planes. **j**) Higher magnification view of white rectangular region in **h**) with membrane pores highlighted (white arrows). **k, l**) Corresponding axial views to **j**), again highlighting the same pores (white arrows). Red arrows: actin encapsulated within membrane; blue arrows: void areas enclosed by membrane; yellow arrows: nonspecific labeling of coverslip with membrane dye. **m**) Higher magnification view of rectangular region in **h**), with accompanying cross-sectional view **n)** corresponding to white dashed line in **m**), highlighting membrane-bound organelles (white arrows). Scale bars: 2 μm **a, f**); 1 μm **b, e, g, i-n**); 10 μm **c, h**); 5 μm **d**). All images were acquired with 4-beam SIM unless otherwise noted.

In a first example, we imaged live, sporulating *B. subtilis* cells expressing GFP-tagged SpoVM and additionally labeled with CellBrite Fix 555 (**Fig. 3a**). SpoVM preferentially binds to micron-scale convex membranes^30^, marking these surfaces to direct assembly of the proteinaceous ‘coat’ surrounding the developing spore^31^. Dual-color 4-beam SIM enabled crisp visualization of SpoVM localized to mature spores within the context of the surrounding membrane in the mother cell. Such detail was lost in widefield imaging, particularly when inspecting axial views (**Fig. 3b**).

Second, we immunolabeled vimentin intermediate filaments and microtubules in fixed U2OS cells and imaged these dense cytoskeletal networks with dual-color 4-beam SIM (**Fig. 3c-g**). The observed lack of colocalization in axial views (**Fig. 3d**) underscores the axial resolution of our method. Instead of colocalization, we observed an apparent ‘stratification’ of vimentin and microtubule fibers in the perinuclear area (**Fig. 3e**). Intriguingly, we also observed examples in which local enrichments of vimentin appeared to fill the ‘gaps’ between microtubules (**Fig. 3f, g**). The close network apposition we observe is consistent with a recent live imaging study showing that vimentin filaments can ‘template’ the microtubule network to stabilize cell polarity during migration^32^.

Finally, we imaged Alexa Fluor 568 phalloidin-labeled actin filaments and CellBrite Fix 488-labeled membranes in fixed mouse liver sinusoidal endothelial cells (LSECs) (**Fig. 3h-n, Supplementary Video 6**). In these samples, axial views revealed actin filaments encapsulated within fine membrane protrusions (likely filopodia, **Fig. 3i**) as well as smaller actin enrichments within the lamellar region (**Fig. 3j-l**). LSECs contain nanoscale pores (‘fenestra’) that filter material passing between blood and hepatocytes. Similar to earlier studies using 3D SIM^17^, we resolved vertically oriented pores that appeared to traverse the extent of thin membrane regions, as well as hollow regions (devoid of actin or membrane dye) fully encapsulated within the membrane (**Fig. 3j-l**). Elsewhere, in thicker regions of the cell, we observed many variably sized void regions, with some just larger than our resolution limit and others, microns in diameter (**Fig. 3m, n**). In almost all cases, we discerned enrichment of the CellBrite marker around voids, suggesting they represent membrane-bound organelles.

### An improved deep learning method provides an alternate means to improve axial resolution

In the process of validating our 4-beam SIM system, we imaged hundreds of cells. Two persistent challenges motivated us to consider alternate means of enhancing axial resolution. First, even though careful adjustment and stabilization of the instrument reliably yielded high quality reconstructions (**Figs. 1-3, Supplementary Figs. 10, 14a, b**), we occasionally observed ringing artifacts. Such artifacts were most obvious when imaging fine structures smaller than our resolution limit, such as immunolabeled microtubules. Indeed, given that we mostly observed such artifacts when imaging microtubules at the boundary of the nucleus (**Supplementary Fig. 14c-f**), we suspect that slight variations in refractive index within the cell lead to localized wavefront distortions which in turn produce reconstruction artifacts.

Second, although we were able to successfully image whole living cells (**Fig. 2a-d, h-i**; **Fig. 3a, b**; **Supplementary Fig. 14a, b**), volumetric time-lapse (‘4D’) imaging proved challenging. Even when using the relatively bright and photostable dye Potomac Gold^33^ to ubiquitously label cell membranes in live cells, we found that phototoxicity^34^ limited experiment duration (**Supplementary Fig. 14g, h, Supplementary Video 7**). In hindsight, this is unsurprising: imaging even a modestly sized 4 μm thick volume with 4-beam SIM entails 1,000 raw images (assuming 15 images/plane and our 60 nm axial sampling interval) with each raw image capture illuminating almost the entire volume of the cell.

Given that 3D SIM introduces less dose than 4-beam SIM, is more robust to wavefront distortions (**Supplementary Fig. 14d**), and has been shown to enable sustained 4D imaging^3^, we considered computational strategies for improving the axial resolution of 3D SIM without introducing additional illumination dose. As deep learning has been shown capable of enhancing spatial resolution in fluorescence microscopy^35-39^, we evaluated a method that improves axial resolution by i) blurring and downsampling lateral views to resemble lower resolution axial views and ii) learning to reverse this degradation based on the higher resolution lateral view ground truth^35,40^. The strength of this approach is that the training data itself contains the ground truth. A key assumption is that the structures of interest appear similar regardless of the viewing direction. Indeed, when evaluating the method on simulated data of randomly oriented spheres, shells, and lines that had been blurred with an anisotropic PSF, we were able to restore the structures so that they appeared isotropic (**Supplementary Fig. 16a, b**).

However, when we attempted to restore 3D SIM data using the same method, although the network improved axial resolution for some structures (**Supplementary Fig. 16c**), it also artificially distorted the shape or even lost other structures (**Supplementary Fig. 16d-h**), likely because axial specimen views looked quite different than the lateral specimen views the network was trained on. We reasoned that network output could be improved if the network was directly exposed to axial information during the training process. We thus presented the network with axial (xz) 3D SIM views that had been blurred and downsampled to yield data with isotropic resolution equivalent to that of the axial resolution of 3D SIM. We then trained the network to reverse the degradation along the lateral direction, for which higher resolution ground truth exists (**Supplementary Fig. 17, Methods**). Motivated by a recent pipeline we developed for multiview confocal super-resolution microscopy^41^, applying the trained network on six digitally rotated views of unseen, similarly degraded 3D SIM data then enabled the improvement of 1D resolution along arbitrary directions. Fusing all such resolution-enhanced views so that the best resolution in each view was preserved^42^ yielded a final prediction with isotropic resolution (**Fig. 4a, Methods**).

**Fig. 4.**
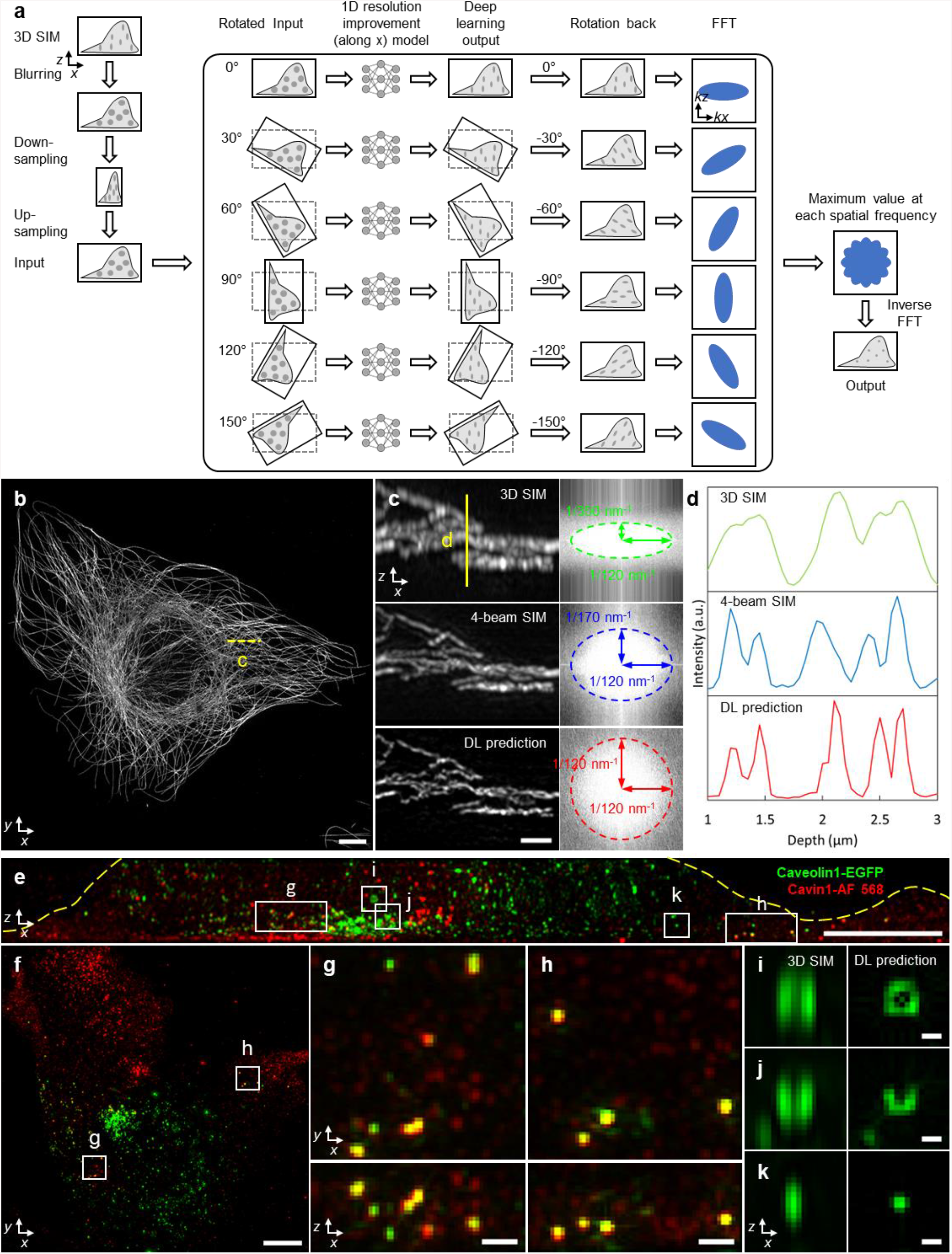
Deep learning for axial resolution enhancement. **a**) Schematic of deep learning process. Three-dimensional SIM (3D SIM) image volumes are blurred, downsampled, and upsampled (each along the lateral x direction) to render isotropic, low-resolution input data (resolution equivalent to axial resolution in 3D SIM). This input is (i) rotated in 30-degree increments; (ii) each rotation passed through a deep learning model to improve resolution along the x direction; (iii) rotated back to the original frame; (iv) Fourier transformed; (v) the maximum value, taken over all rotations, at each spatial frequency recorded, and (vi) finally inverse Fourier transformed to yield an output prediction with improved, isotropic resolution. See also **Supplementary Fig. 17, Methods. b**) Alexa Fluor 488 immunolabeled microtubules in a fixed U2OS cell. Maximum intensity projection of deep learning (DL) prediction is shown. **c**) Left: higher magnification axial view indicated by yellow dotted line in **b**), comparing 3D SIM (top), 4-beam SIM (middle), and DL prediction (bottom). Images are generated by computing maximum intensity projection over 20 pixels in the y direction. Right: magnitudes of Fourier transforms corresponding to images at left, with indicated spatial frequencies bounding the major and minor ellipse axes. **d**) Line profiles corresponding to yellow solid line in **c**). **e, f**) Fixed mouse embryonic fibroblast with Caveolin-1 EGFP with GFP booster (green) and Alexa Fluor 568 immunolabeled Cavin-1 (red). Axial **e**) and lateral **f**) maximum intensity projections of DL predictions are shown. Yellow dotted line in **e**) has been added to better delineate cell boundary. **g, h**) Higher magnification views of rectangular regions in **e, f**) with lateral (top) and axial (bottom) projections indicated. **i, j, k**) Higher magnification axial views of regions indicated in **e**), comparing 3D SIM input (left) to DL prediction (right). See also **Supplementary Fig. 19**. Scale bars: 5 μm **b, e, f**); 1 μm **c**); 500 nm **g, h**); 200 nm **i, j, k**).

Comparing images of the same sample produced by 3D SIM, 4-beam SIM, and our modified deep learning prediction (**Fig 4b-d**) facilitated validation of our method. For example, when inspecting immunolabeled microtubules in fixed U2OS cells (**Fig. 4b**), although all three methods offered similar lateral resolution (**Fig. 4c**), fine axial features blurred in 3D SIM were resolved with 4-beam SIM and the network prediction, which showed close visual (**Fig. 4c**) and quantitative (**Fig. 4d**) agreement. We obtained similar results on membrane-stained, live *B. subtilis* and immunolabeled Tomm20 in fixed U2OS cells (**Supplementary Fig. 18**).

Next, we performed two-color imaging of Caveolin-1 and Cavin-1, components of the caveolar coat. Caveolae are 70-100 nm diameter membrane invaginations that can detach from the plasma membrane and move through the cytoplasm, playing key roles in lipid metabolism and trafficking^43^. We fixed mouse embryonic fibroblasts expressing Caveolin-1-EGFP and additionally immunolabeled Cavin-1 with Alexa Fluor 568, performed 3D SIM imaging, and then applied our network to the 3D SIM images (**Fig. 4e-k**). Caveolin-1 and Cavin-1 labels mostly marked distinct caveolae pools (**Fig. 4e, f**), although we also observed a smaller pool of caveolae puncta that displayed colocalized signal (**Fig. 4g, h**). Unlike the Cavin-1 signal, which mostly decorated structures sized at or below our resolution limit, Caveolin-1 appeared to label a more heterogenous pool of caveolae (**Fig. 4i-k**). Hints of such heterogeneity existed in the input 3D SIM data, but were obscured by diffraction. By contrast, the network prediction appeared to resolve ring-shaped structures (**Fig. 4i**), partial rings (**Fig. 4j**), and spherical puncta (**Fig. 4k**). We also found that Caveolin-1 localized to larger ring-shaped structures of varying size, possibly lipid droplets (**Supplementary Fig. 19**).

### Multi-step deep learning enables 4D super-resolution imaging with isotropic spatial resolution

One of the easiest ways to extend imaging duration in fluorescence microscopy is simply to lower the intensity of the illumination. This approach is ultimately limited by the SNR of the raw data, a particularly important constraint in conventional 3D SIM reconstruction, which is highly susceptible to noise^44^. In response to this challenge, several recent studies have employed deep learning to denoise SIM data^38,45-47^. Motivated by this prior work, we developed a multi-step denoising approach (**Fig. 5a, Supplementary Fig. 20a, Methods**). First, we gathered matched pairs of low and high SNR volumes (∼10-fold difference in illumination intensity), training a residual channel attention network (3D RCAN^37^) to denoise low SNR volumes that would, without denoising, produce unacceptably noisy 3D SIM reconstructions. Next, we applied a generalized Wiener filter to produce an intermediate 3D SIM reconstruction from the denoised low SNR volumes. Finally, we trained a second 3D RCAN to additionally denoise the intermediate 3D SIM reconstruction, further reducing patterned noise artifacts (**Fig. 5b-d, Supplementary Fig. 20b, c**). This procedure provided reconstructions that were visually and quantitatively superior to those produced by all other strategies we tested (**Supplementary Figs. 21, 22, Supplementary Tables 2, 3**), including using only the first 3D RCAN and subsequent Wiener filter; modifying the RCAN to incorporate all 15 low SNR input volumes to directly (without a Wiener filter) predict the 3D SIM reconstruction; using DenseDeconNet^48^, a different type of network; or eliminating the Wiener filter between the sequential 3D RCAN networks. In this work, we built multistep denoising models for outer mitochondrial membranes (Tomm20), lysosomal membranes (LAMP1), interior lysosomal markers (LysoTracker Red), and microtubules.

**Fig. 5.**
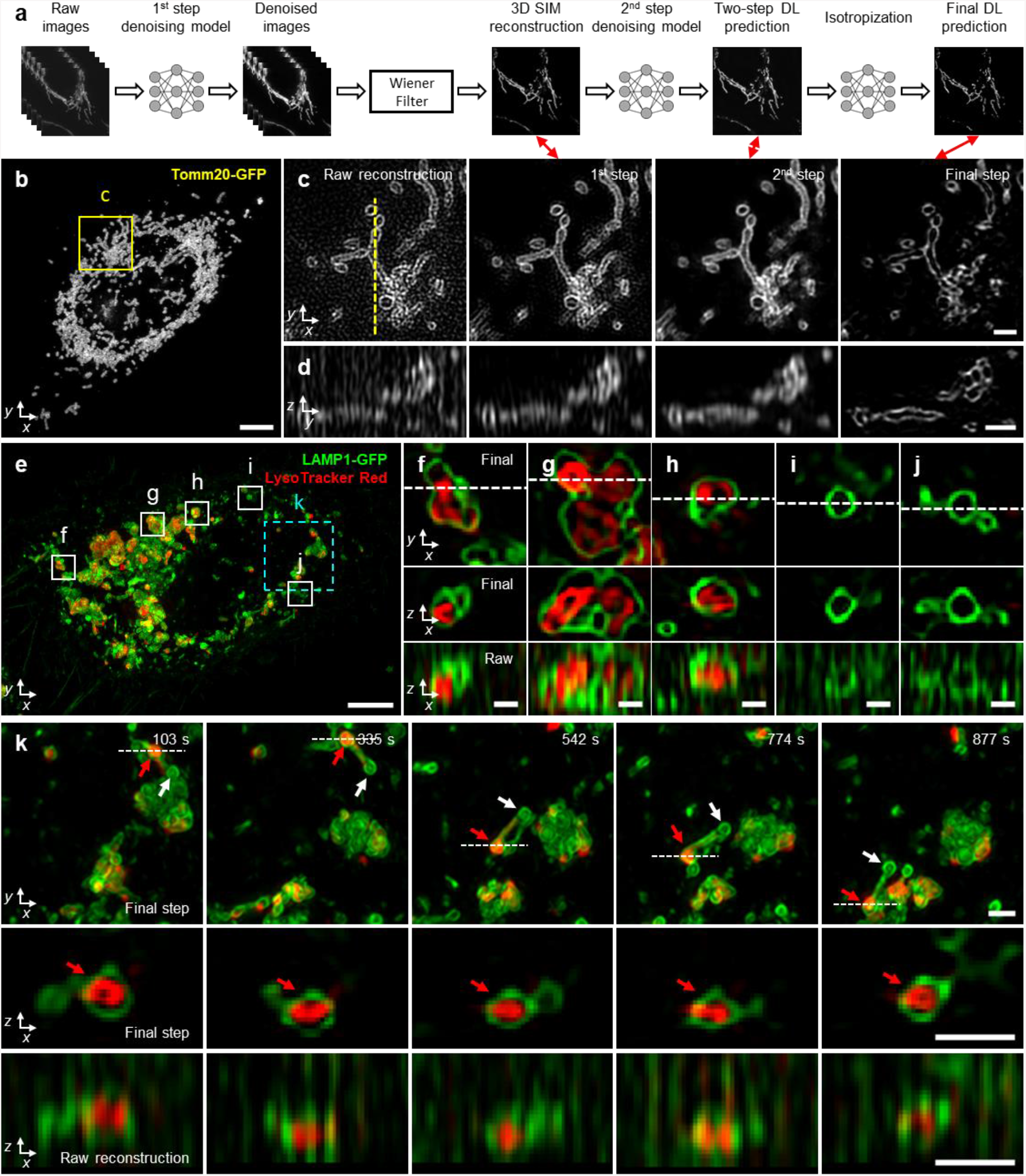
Denoising and axial resolution enhancement facilitate 4D super-resolution imaging with isotropic resolution. **a**) Schematic illustrating workflow for applying deep learning to raw input data. Sets of raw images (5 phases x 3 orientations) are denoised, combined with a generalized Wiener filter, the resulting 3D SIM reconstruction denoised, and passed through our axial resolution enhancement workflow (**Fig. 4a**) to yield an isotropic, denoised, super-resolution prediction. See also **Fig. 4a, Supplementary Fig. 17, 20. b**) Maximum intensity projection of final prediction for Tomm20-GFP label in a live U2OS cell, 25th time point from 50-time point volumetric series. See also **Supplementary Video 8. c**) Single lateral plane corresponding to yellow rectangular region in **b**), illustrating progressive improvement from 3D SIM reconstruction based on raw input data; after first denoising model and Wiener filter; after applying second denoising model; and after isotropization model. Double-headed red arrows show corresponding steps in schematic **a**). **d**) As in **b**), but for axial plane indicated by yellow dashed line in **c**). **e**) Maximum intensity projection of final prediction for live U2OS cell expressing lysosomal marker LAMP1-GFP (green) and additionally labeled with LysoTracker Red to mark the lysosome interior (red). First time point from 60-time point volumetric series is shown, see also **Supplementary Video 10. f-j**) Higher magnification views of white rectangular regions in **e**), illustrating lateral views (top), axial views along white dashed lines in lateral views (middle), and comparative axial views from 3D SIM reconstructions (bottom, ‘Raw’). **k**) Higher magnification view of cyan dashed rectangular region, emphasizing dynamics at selected time points. See also **Supplementary Video 12**. Top: lateral views, red arrow emphasizes lysosomal subregion filled by LysoTracker Red dye, vs. white arrow indicating unfilled region; middle: axial view corresponding to white dashed lines; bottom: comparative 3D SIM axial view. Scale bars: 5 μm **b, e**); 1 μm **c, d, k**); 500 nm **f-j**).

Our multi-step denoising pipeline produced high-quality 3D SIM predictions that could be directly passed to our axial resolution enhancement method (**Fig. 5a**), thereby producing high SNR reconstructions with ∼120 nm isotropic resolution from low SNR input. The improvement in SNR and resolution offered by deep learning was particularly striking when visualizing subcellular organelles and their dynamics. For example, by lowering the illumination intensity to ∼0.5 W/cm^2^, we could perform 3D SIM imaging of EGFP-Tomm20 in live U2OS cells (**Fig. 5b, Supplementary Video 8**) for 50 volumes without observing significant photobleaching or obvious signs of phototoxicity. Performing the direct 3D SIM reconstruction on the low SNR raw data revealed the expected Tomm20 signal at the outer mitochondrial membrane in lateral views, although the signal was contaminated with patterned noise (**Fig. 5c**) and obscured by diffraction in axial views (**Fig. 5d**). Denoising through successive 3D RCANs progressively improved SNR, although only the final step provided an axial clarity commensurate with lateral views (**Fig. 5d, Supplementary Video 8**). We observed similar improvements in SNR and axial resolution when imaging lysosomal dynamics with an EGFP-LAMP1 marker in live U2OS cells (**Supplementary Video 9**).

To demonstrate the potential of our deep learning pipeline for live dual-color imaging, we performed volumetric imaging over 60 time points of U2OS cells expressing EGFP-LAMP1, marking lysosomal membranes, and additionally labeled with LysoTracker Red dye, which labeled the interior of lysosomes (**Fig. 5e-k, Supplementary Videos 10**,**11**). Although some lysosomes appeared as ‘textbook’ discrete vesicular structures (**Fig. 5i, j**), our reconstructions revealed considerable structural heterogeneity, with some lysosomes assuming a ‘multibud’ structure (**Fig. 5f, g**) and others appearing as tubules (**Fig. 5k**). Interestingly, Lysotracker Red localized to the interior of many, but not all, lysosomes. In some cases, LysoTracker Red preferentially localized to different membrane-bound regions even within a single lysosomal structure (**Fig. 5f, g, k**). Such compartmentalized staining appeared stable over our ∼25-minute recording, suggesting limited turnover of the dye despite rapid dynamics of the parent structure (**Fig. 5k, Supplementary Video 12**). Given that LysoTracker Red is known to stain acidic organelles, perhaps this result reflects underlying pH differences between and even within lysosomes. Regardless, such fine structural details were largely obscured in axial views of 3D SIM reconstructions derived from the raw data.

Finally, we investigated microtubule dynamics in Jurkat T cells as they were activated and spread on anti-CD3 coverslips (**Fig. 6, Supplementary Videos 13-16**). This system has been used extensively to study the cytoskeletal remodeling that takes place during the early stages of immunological synapse formation, including actin ring formation and centrosome polarization^49-51^. By lowering the illumination intensity sufficiently, we recorded a 3D SIM time-series spanning 100 volumes (one volume every 12.8 s) without significantly bleaching our EMTB 3x GFP microtubule marker or introducing noticeable phototoxicity. This ∼21-minute duration recording proved long enough to observe dramatic remodeling of the cytoskeleton, particularly after applying our denoising and axial resolution enhancement pipeline (**Fig. 6a, b**). The deep learning prediction revealed finer detail and many more filaments than 3D SIM reconstructions based on the raw input data, which were degraded by noise and relatively poor axial resolution (**Fig. 6c, d**). In turn, this improvement in image quality facilitated inspection of the rapidly remodeling microtubule network and was particularly helpful in elucidating the interplay of distinct cytoskeletal elements occurring perpendicular to the coverslip surface. For example, we segmented two microtubule filaments (**Fig. 6b**) that appeared to grow towards each other, apparently merging and encircling the nucleus before disassociating (**Fig. 6e, Supplementary Video 13**). We also identified the centrosome, tracking its ‘downward and inward’ movement from its original peripheral location towards the coverslip. Centrosome polarization in T-cells has been described as biphasic and driven by the association of microtubule bundles with molecular motors (dynein) at the synapse^51,52^. Intriguingly, centrosome repositioning correlated closely with the inward movement of one of the segmented microtubule bundles (**Fig. 6f, Supplementary Video 14**), suggesting close physical coupling between different parts of the network, perhaps resulting from the coordinated action of microtubule associated motors. We also observed numerous ‘buckling’ events of individual microtubule filaments (**Fig. 6g, Supplementary Fig. 23, Supplementary Videos 15, 16**), presumably also the result of active force generation by molecular motors^49^.

**Fig. 6.**
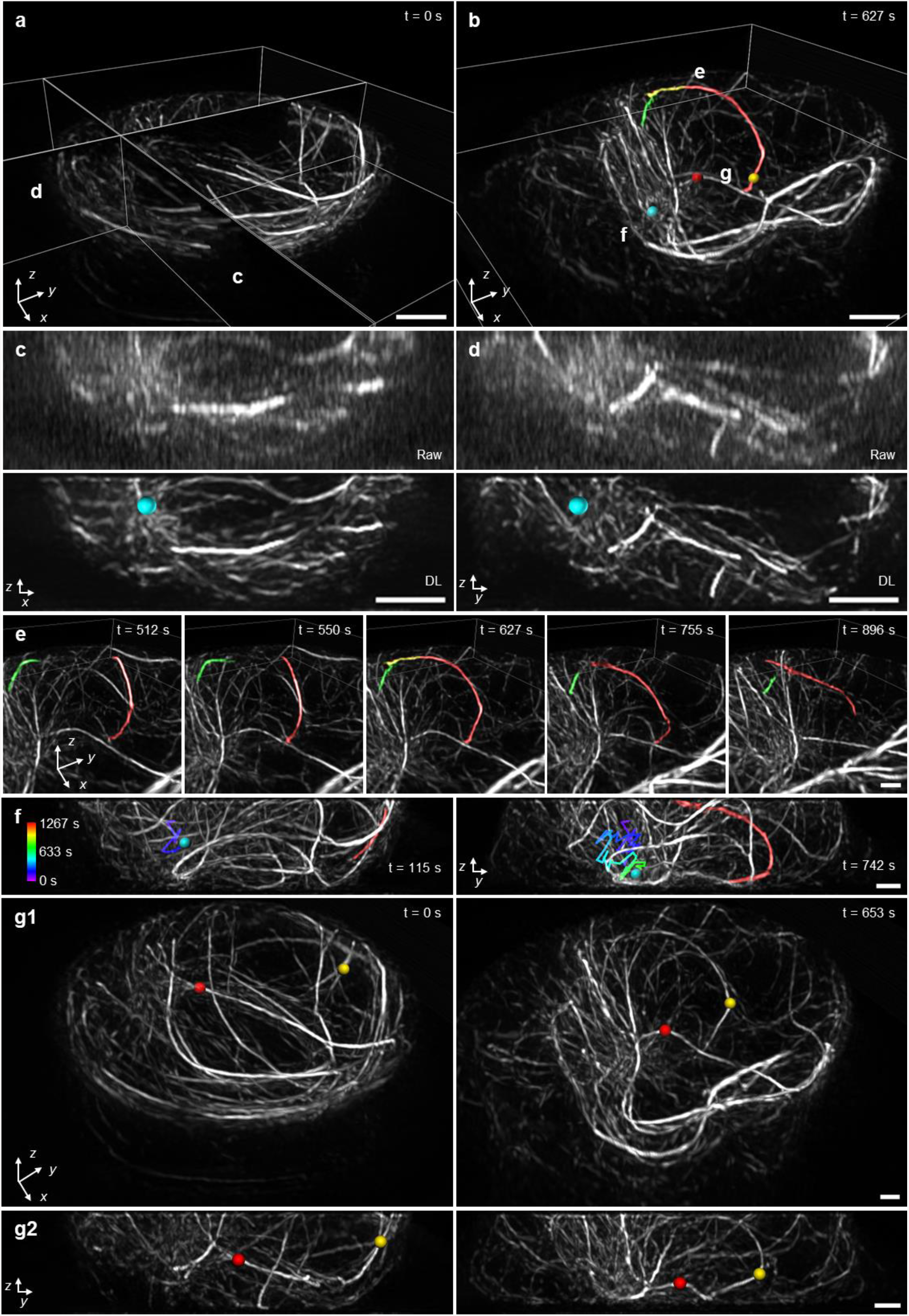
Denoising and axial resolution enhancement unveil rich microtubule dynamics within a living immune cell. **a, b**) Selected volumetric reconstructions of Jurkat T cells expressing EMTB 3x GFP are shown from 100 time point series (volumes recorded every 12.8 s), in perspective views. In **b**), the centrosome (cyan sphere), overlapping microtubules (red, yellow, green), and buckling microtubules (red and yellow spheres) are shown. **c, d**) Comparative axial views (maximum intensity projections over 2 μm thickness in ‘y’) of 3D SIM reconstructions (‘Raw’, upper row) and deep learning output (‘DL’, bottom raw) corresponding to planes in **a)**. Position of centrosome is indicated (cyan sphere). **e**) Selected time points corresponding to subregion marked in **b**), emphasizing two microtubule filaments that are initially separated (t = 512, 550 s), merge over the nucleus (t = 627 s), and separate again (t = 755 s, 896 s). See also **Supplementary Video 13. f**) Axial views (projections over 8 μm thickness) that emphasize correlated, inward movement of centrosome (cyan sphere, previous trajectory temporally coded as indicated in color bar) and microtubule filament (red, the same filament shown in **b, e**). See also **Supplementary Video 14. g**) Lateral (**g1**) and axial views (**g2**) emphasizing buckling of two microtubule filaments (red and yellow spheres, as shown in **b**)). Pre-buckling: left columns; post-buckling: right columns. Both **g1** and **g2** are projections over 8 μm thickness. See also **Supplementary Video 15**. Scale bars: 2 μm **a-d**); 1 μm **e-g**).

## Discussion

Given the three-dimensional nature of most biological samples and the inherent anisotropy of the PSF, improving the axial resolution of fluorescence microscopy can directly translate into new biological insight. Thus far, however, most methods have focused on improving lateral, rather than axial resolution. Here we addressed this issue by markedly improving the axial resolution of 3D SIM^1^, a broadly used super-resolution method well-suited for studying single cells. First, we showed that adding a mirror to a 3D SIM system enables near-isotropic spatial resolution. Besides its simplicity, the main advantage of this ‘physics-based’ 4-beam SIM method is that it does not require prior information about the sample. By contrast, the second computational method can be applied to 3D SIM systems without hardware modification. Instead, prior sample information is embedded into a series of neural networks, which can then predict a denoised image reconstruction with isotropic spatial resolution. As emphasized here, these techniques provide distinct means for axial resolution enhancement, yet in principle they could be combined, e.g., denoising the input to 4-beam SIM so that illumination intensity may be lowered, thereby reducing phototoxicity and improving the technique’s potential for 4D imaging.

Both methods can be further improved. The 4-beam SIM system requires active drift correction and more precise alignment of the illumination pattern with the focal plane than 3D SIM. While our bead-based algorithm (**Supplementary Fig. 12**) met both requirements, hardware-based correction^53^ would provide faster feedback, possibly obviating the need to image beads, and thus lowering the total illumination dose imparted to the sample. For applications in which the axial view alone is sufficient, collecting only five images (i.e., one orientation, five phases) would provide optical sectioning and axial resolution enhancement, improving speed (and reducing dose) by 3-fold. Also, for multi-color applications, the technique could benefit by implementing independent detection paths for each color, bypassing the time-consuming need to realign the detection path as manually implemented here.

The multi-step deep learning pipeline is currently time-consuming to implement, requiring the collection of ∼50 volumetric pairs per network, and ∼12 hours for training all networks. Once trained, application of the networks is faster, requiring ∼5 minutes per volume (each 500 × 500 × 80 voxels). Given continued development in the rapidly growing field of deep learning, improved networks with fewer parameters^54^ are likely to significantly shrink these times in the future. In considering our multi-step approach, we maintained as network input the 15 raw image volumes required for traditional 3D SIM reconstruction. We note that one route to faster and more gentle imaging is to train the network to accept fewer input images (e.g., via fewer phases or coarser axial sampling), although the quality of such reconstructions is currently inferior to that obtained using the entire stack^47^. Perhaps the largest caveats with any deep learning method are that the quality of the prediction is tied to the quality of the training data, and that generalizability to data unlike that of the training set remains questionable. Finally, we note that the spatial resolution of both the 4-beam and deep learning approaches may be improved by incorporating photoswitching^55^, albeit with accompanying reduction in temporal resolution and increase in illumination dose.

Despite these qualifications, in their current form, the methods we present outperform recent state-of-the-art SIM implementations. Relative to a recent implementation of lattice light-sheet microscopy employing structured illumination for lateral resolution enhancement and coherent detection for improved axial resolution (3D-iLLS)^56^, our techniques offer a 2-3 fold improvement in acquisition speed, a 2-3 fold improvement in volumetric resolution, a ∼40-fold improvement in imaging volume size, an order of magnitude more time points (using deep learning) and less susceptibility to reconstruction artifacts. Relative to live-cell PA-NL SIM LLSM^57^, our techniques offer similar speed and resolution, a ∼1.5-fold improvement in imaging volume size, 2.5-5 fold more timepoints (using deep learning), and achieves this performance without a photoswitchable fluorophore or a multi-objective imaging system. In addition to the gains on conventionally prepared samples illustrated here, our methods may improve SIM reconstructions on samples prepared for correlative super-resolution fluorescence and electron microscopy^58^, which currently suffer from poor axial resolution. Finally, although we focused on improving the performance of 3D SIM in this work, we suspect our methods may prove useful in improving the axial resolution of other 3D imaging techniques.

## Author Contributions

Conceived project: X.L., Y.W., H.Shroff. Performed simulations of microscope: X.L. with guidance from P.L.R., and H.Shroff. Wrote acquisition code: X.L. Designed optical layout: X.L. and H.Shroff. Built optical system: X.L. Built an early prototype of the system: J.P.G., H.D.V., J.C. Wrote reconstruction software: X.L. with guidance from L.S. and P.L.R. Designed and implemented deep learning pipeline: Y.W. with input from X.L. and H.Shroff. Modified previous deep learning methods: Y.L., H.Sasaki. Assisted with registration: M.G. Designed experiments: X.L., Y.W., Y.S., I. R-S., C.M., T.B.U., Z.W., K.S.R., J.W.T., A.U., H.Shroff. Prepared samples: X.L., Y.S., I.R-S., C.M., T.B.U., Z.W., L.Z. Performed experiments: X.L., with guidance from Y.S., I.R-S., C.M., T.B.U., Z.W., H.Shroff. Analyzed data: X.L., Y.S., I.R-S. and H.S. with input from all authors. Wrote paper: X.L. and H.Shroff, with all authors participating in the drafting process. Supervised research: S-J. J.L., L.S., H.L., K.S.R., J.W.T., A.U., P.L.R., and H.Shroff. Directed overall project: H.Shroff.

## Acknowledgements

We thank Jiamin Liu for helping to test some of the deep learning models used in this work and for critical feedback on the manuscript, Arthur Melo and Oliver Daumke for the gift of mouse embryonic fibroblasts, Richard Lundmark for the Caveolin1-EGFP plasmid, George Patterson’s lab for the LAMP1-EGFP plasmid, Robert Fischer for helpful samples used in early benchmarking of microscope performance, Reto Fiolka and Talley Lambert for discussions on SIM, Keir Neuman for useful discussions about microscope stability and interferometry, Fred Lanni for sharing his deep knowledge of standing wave microscopy, Luke Lavis for advice about dyes and for the gift of Potomac Gold, Applied Scientific Instrumentation for help troubleshooting piezo stage issues, Yves Pommier’s lab for assisting with cell sorting for the U2OS stable cell lines, and David Ide for his superb machining skill and for making the pinhole mask used in this work. This research was supported by the intramural research programs of the National Institute of Biomedical Imaging and Bioengineering; the National Institute of Heart, Lung, and Blood; and the National Cancer Institute within the National Institutes of Health. A.U. acknowledges support from the grants NIH R01GM131054 and NSF PHY-1915534. We thank the Office of Data Science Strategy, NIH, for providing a seed grant enabling us to train deep learning models using cloud-based computational resources. H.L. acknowledges support from the National Key Research and Development Program of China (No. 2020AAA0109502) and the Talent Program of Zhejiang Province (2021R51004). H.S. and P.L.R. acknowledge the Whitman and Fellows program at MBL for providing funding and space for discussions valuable to this work.

## Methods

### Simulations of OTF support

The simulated 2D optical transfer function (OTF) supports (**Supplementary Figs. 1, 2**) were assembled based on imaging parameters and geometric considerations. The 3D OTF support of the widefield microscope is a toroidal solid, whose 2D analog in the *k*_*r*_, *k*_*z*_ plane consists of the area enclosed by four arcs (**Supplementary Fig. 1a**). The position and extent of these arcs in the spatial frequency domain were computed by considering the detection NA and emission wavelength of the widefield imaging system. The 3D SIM OTF was created by placing widefield OTFs at each 3D SIM illumination frequency component (**Supplementary Fig. 1b**). The standing wave microscope OTF support consists of the widefield OTF and two duplicates along the *k*_*z*_ axis, positioned at the spatial frequencies of the standing wave determined by the excitation wavelength and refractive index (**Supplementary Fig. 1c**). The OTF support for 4-beam SIM can be similarly derived by considering the area enclosed by widefield OTFs placed at the seven 3D SIM illumination frequency components, the standing wave spatial frequencies, and four additional frequency components determined by interference of the reflected beam with the two side beams (**Supplementary Fig. 1e**). Finally, as previously described^20^, the I^5^S OTF support is determined by considering the area enclosed by the placing the I^2^M OTF^59^ at each of the 19 illumination components produced in I^5^S (**Supplementary Fig. 1d**). 3D OTF supports were simulated by converting each 2D coordinate in the *k*_*r*_, *k*_*z*_ plane to a corresponding set of 3D spherical coordinates (**Supplementary Fig. 2**). Scripts for performing simulations are included as **Supplementary Software**.

### Homebuilt 3D SIM system

Our 3D SIM optical layout (**Supplementary Fig. 3**) was inspired by previous designs^1,3,4^. Two linearly polarized lasers (488 nm and 561 nm, Coherent, Sapphire 488 LP-300 mW and Sapphire 561 LP-200 mW) were combined via a 3-mm thick dichroic mirror (DM1: Semrock, Di03-R405/488/532/635-t3-25×36) and passed through an acousto-optic tunable filter (AOTF, AA Opto-Electronic, AOTFnC-400.650-TN) for rapid shuttering and intensity control. Illumination power was measured after the objective and the computed intensity at the sample plane varied between 0.5 – 25 W/cm^2^ based on a circularly illuminated area with diameter 90 μm. The first-order beam exiting the AOTF was selected (the zero-order beam was blocked by a beam dump, BD), expanded (L1 & L2: Thorlabs, TRH127-020-A-ML & ACT508-400-A-ML) and spatially filtered by a pinhole (P: Thorlabs, P30K), resulting in a beam with 15 mm 1/*e*^2^ diameter. The excitation beams were then redirected onto a phase-only nematic spatial light modulator (SLM, Meadowlark Optics, MSP1920-400-800-HSP8) at near normal incidence (< 6 degrees). A half wave plate (HWP: Thorlabs, AHWP10M-600) positioned prior to the spatial filter was used to adjust the direction of linear polarization, aligning it for maximum phase modulation by the SLM, thereby ensuring high contrast for the 15 patterns used in 3D SIM. An adjustable iris (Thorlabs, SM1D12) between the beam expander and SLM was set to 10 mm diameter, slightly shorter than the short edge of the SLM active area (10.6 mm). Expanding the beam while underfilling the SLM improves illumination uniformity while avoiding diffraction effects from the SLM boundary.

Lens L3 was positioned at one focal length after the SLM, producing a Fourier image of the illumination pattern at its focus. A pinhole mask (PM) placed at this plane served to filter out unwanted illumination orders due to SLM pitch and pattern pixelization. The spatially filtered beams emerging from the PM were imaged via another pair of relay lenses (lens pair L4 and L5 placed in 4f configuration) onto the back focal plane (BFP) of the objective lens. For each pattern orientation, the coherent 0^th^ and ±1^st^ order beams were collimated by the objective and interfered at the sample plane to form the 3D SIM illumination pattern. A liquid crystal polarization rotator (LCPR, Meadowlark Optics, LPR-200-0525-ACHR, achromatic) was used for rapid rotation of the polarization state post-SLM, producing s-polarized illumination at the sample and thus high illumination pattern contrast there. If using a silicone oil objective lens (Olympus, UPLSAPO 100x/1.35 NA), we used *f*_3_ = 250 mm, *f*_4_ = 300 mm, *f*_5_ = 250 mm (Thorlabs, AC508-250-A-ML & AC508-300-A-ML) producing a demagnification of 115.7 from the SLM to sample plane, given that *f*_*obj*_ = 1.8 mm. When using a water objective lens (Nikon, CFI SR Plan Apo 60x/1.27 NA), *f*_3_ = 300 mm, *f*_4_ = 250 mm, *f*_5_ = 250 mm, producing a demagnification of 90.1 from SLM to sample plane given that *f*_*obj*_ = 3.33 mm. In both cases, the usable field of view (FOV) was at least 90 μm by 90 μm. In addition to FOV, several other design criteria informed our choice of *f*_4_-*f*_5_. First, we picked *f*_3_ to be sufficiently large (>200 mm) to i) ensure sufficient room for near normal incidence of the illumination beam at the SLM and ii) clearly separate the Fourier components of the illumination at the PM, allowing clean filtering of these components relative to background orders. Similarly, we picked *f*_5_ to be long enough (>200 mm) to accommodate a turning mirror and the dichroic mirror.

Fluorescence was isolated post-objective via a dichroic mirror (DM2: Semrock, Di03-R488/561-t3-25×36) and imaged to a scientific complementary metal-oxide-semiconductor detector (sCMOS: PCO, Edge 4.2HQ) mounted on a multi-axis translation stage (Thorlabs, XR25P-K1) and a vertical-travel platform (Thorlabs, L490) via tube lens L6. Emission filters mounted in a filter wheel (FW: Applied Scientific Instrumentation, FW-1000) served to further reject illumination light and select appropriate spectral bands. In this work, two bandpass emission filters (Semrock, FF03-525/50-25 & FF02-617/73-25) and one notch emission filter (Semrock, NF03-405/488/561/635E-25) were used, depending on the sample. Bandpass filters were used when imaging yellow-green beads, red beads and all biological samples to avoid cross talk between spectral bands. The notch filter was used only when imaging orange beads to align the 4-beam SIM system for 2-color applications. When using the 1.35 NA silicone oil objective lens, we chose *f*_6_ = 165 mm (Thorlabs, TTL165-A). When using the 1.27 NA water objective, we chose *f*_6_ = 265 mm (Applied Scientific Instrumentation, C60-TUBE-265D). The resulting image pixel sizes for the silicone oil lens were 70.9 nm (91.7x magnification from sample to camera), and 81.8 nm (79.5x magnification from sample to camera) for the water lens. In both cases, the image pixel sizes were smaller than the Nyquist limit.

Sample and objective were held in a modular microscope frame with a motorized XY stage (Applied Scientific Instrumentation, RAMM and MS-2000 XYZ Automated Stage) used for lateral sample positioning and coarse focusing. A Z piezo stage (Applied Scientific Instrumentation, PZ-2150, 150 μm axial travel) attached to the stage was used to provide precise axial sample positioning (125 nm step size for 3D SIM, 60 nm step size for 4-beam SIM to ensure Nyquist sampling in z). Samples were deposited on high precision 25-mm coverslips (Thorlabs, CG15XH) that were mounted in a magnetic imaging chamber (Warner Instruments, QR-40LP) filled with imaging medium. The chamber was placed into a stage insert (Applied Scientific Instrumentation, I-3091 universal insert) mounted to the piezo stage (**Supplementary Fig. 9a**).

### 3D SIM pattern generation

As in previous 3D SIM systems, we used the SLM as a binary phase grating (each pixel producing a phase retardance of 0 or π radians) to generate periodic illumination patterns at the sample plane. To generate appropriate patterns, we carefully considered how to implement: i) the desired pattern orientations, ii) the desired line spacing (grating period) in each pattern, iii) the duty cycle appropriate for each pattern, and iv) the relative 2π/5 phase shifts between each of the five patterns required at each orientation.

First, 3D SIM typically uses patterns with three orientations spaced 60° apart to i) achieve near-isotropic lateral resolution and ii) fill in the ‘missing cone’ of axial spatial frequencies, thereby providing optical sectioning. In the pixelated coordinate system of the SLM (**Supplementary Fig. 4a**), we found it convenient to define a vector 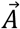, described by integer components (*A*_*x*_, *A*_*y*_), to specify pattern orientation. In this work, we chose 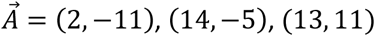 as the three pattern orientations, which corresponds to 10.3°, 70.3° and 130.2°. This choice allowed us to pick a grating period with a non-integer value, unlike orientations at 0° or 90°, which would restrict the grating period to integer values. We also found that 45° and 135° orientations should be avoided, as they caused many additional orders between the zero and first orders at the Fourier plane, making the filtering at the PM less efficient.

Second, 3D SIM uses three tightly focused beams at the BFP of the objective lens to produce the illumination pattern. The positions of the two side beams are typically located at 90-95% of the pupil radius. Using a higher radius (>95% of the pupil) decreases the amplitude of the highest lateral illumination spatial frequency to the point that it is difficult to detect, complicating conventional SIM reconstruction algorithms which rely on precise estimation of this parameter. On the other hand, using a substantially lower radius (<90% of the pupil) needlessly decreases resolution, especially along the axial dimension. In this work, we sought to position our side beams at 92% of the pupil radius, thereby achieving easily detectable pattern modulation while maintaining high resolution.

After the ratio of the side beam position to pupil radius of BFP *r* and system demagnification factor *M* from the SLM to the sample plane are determined, the corresponding SLM pattern period can be computed as 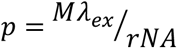, in which *λ*_*ex*_ is the excitation wavelength and *NA* is the numerical aperture of the objective. Taking *λ*_*ex*_ = 488 *nm* for example, the pattern period is 45.5 μm for the 1.35 NA silicone oil lens (*M* = 115.7) and 37.6 μm for the 1.27 NA (*M* = 90.1) water lens. Considering the 9.2 μm pixel size of the SLM, these periods correspond to 4.95 pixels and 4.09 pixels respectively. In practice, the period was fine-tuned for each of the three pattern orientations and different wavelengths, with the goal of achieving *r* = 0.92. The positions of the 1st order components of the illumination pattern at the PM are given by 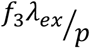. We machined the PM from a 1” diameter, ∼1 mm thick aluminum disk, creating 7 holes in the disk to selectively transmit only the zero order and first order components of the illumination pattern (each hole 0.5-1 mm in diameter, one centered and the six spaced at 60 degree intervals surrounding the central spot). The PM was finely rotated using a mount (Thorlabs, CRM1) to ensure maximal throughput of the illumination pattern.

Third, 3D SIM and 4-beam SIM both require the central, zero-order illumination beam to generate the interference pattern at the sample. The relative intensity of the zero-order beam can be controlled by modifying the duty cycle of the SLM pattern, defined by the fraction of pixels in the on-state (π phase retardance) in each period. We set the duty cycle of the SLM pattern to ∼30%, resulting in a zero-order intensity that was 70-75% of the first-order intensity. This ratio was chosen to emphasize the relatively weak amplitudes of the highest lateral spatial frequencies in the illumination pattern, as inspired by previous work^1,3^. In practice, the duty cycle was fine-tuned for each pattern orientation and laser wavelength to maintain the desired intensity ratio (33% and 31% duty cycle for 488 nm and 561 nm lasers, respectively).

With these considerations in mind, to generate SLM patterns with the appropriate orientation, period, duty cycle, and relative phase as required for 3D SIM, we adopted the following pattern finding algorithm (**Supplementary Fig. 4**): i) Pick a vector 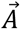 parallel to the desired pattern, where the pattern consists of stripes separated by periodicity *p*. 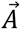 defines the direction (orientation) of the pattern. ii) Obtain the vector 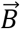 perpendicular to 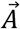 with length *p*. iii) For any pixel coordinate (*x, y*) on the SLM, compute the scalar projection 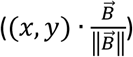 of vector (*x, y*) onto vector 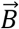. iv) Compute the modulo (MATLAB function ‘mod’) after dividing the projection by *p*. If the modulo is greater than the duty cycle multiplied with *p*, a gray level corresponding to 0 phase retardance is assigned to the current pixel. If the modulo is less than the duty cycle multiplied with *p*, a gray level corresponding to π phase retardance is assigned. v) Repeat the above procedure to find the patterns corresponding to the other phase steps simply by moving the origin of vector (*x, y*) along 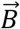 by the length of the desired phase step. The advantage of this method is that period and duty cycle may be independently tuned without affecting the pattern orientation. We thus found it very useful in fine-tuning the pattern parameters for e.g., multi-color imaging, without needing to change the physical properties of the PM.

Previous 3D SIM work^3,4^ used a binary ferroelectric SLM in which each pixel is set to a binary state, resulting in either 0 or π phase retardance. Here we used a nematic SLM, which allows a greater range of phase values. However, nematic SLMs must be calibrated to generate a look up table (LUT) that maps the input 0 - 255 values to the grayscale value that results in a linear 0 – 2π phase retardance. Prior to use, we calibrated the SLM for 488 nm and 561 nm wavelengths, following the manufacturer’s guidelines, subsequently selecting the LUT appropriate for the experiment at hand.

To ensure the illumination pattern has maximum contrast at the sample plane (s-polarization), the LCPR’s state was dynamically adjusted to keep the polarization of illumination beams parallel to vector 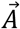 (orthogonal to vector 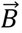) for each of the three pattern orientations. Fine alignment of the LCPR optical axis was accomplished using the following procedure: i) Place the LCPR in a rotation mount (Thorlabs, RSP2D) with one of its input axes roughly aligned with the polarization vector of the illumination post-SLM. ii) Place a linear polarizer after the LCPR with transmission axis in the same direction of vector 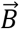. iii) Slowly rotate the LCPR until the transmitted intensity is minimized. Then fine tune the control voltage to the LCPR to further reduce the transmitted intensity. iv) Repeat step iii) several times to ensure that the LCPR is properly aligned for the current pattern orientation and repeat step ii) with polarizer axis rotated appropriately to match 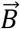 for the other two orientations. v) Remove the polarizer and record the LCPR input voltages for three pattern orientations, then apply the appropriate voltages to the LCPR during 3D SIM and 4-beam SIM experiments (**Supplementary Figs. 6, 7a**).

### 3D SIM system alignment

We implemented a series of checks to verify the alignment of our 3D SIM system. First, we minimized aberrations in the widefield point-spread function (PSF). We used the SLM to produce widefield illumination by setting all pixels to 0 phase, before acquiring a stack of a single 100 nm bead placed in center of the FOV. The stack was acquired with near-isotropic voxel size. When projecting the stack to visualize the axial views (XZ and YZ), we checked for symmetry of the PSF along each direction. By adjusting the four screws of the ASI stage insert and the correction collar, we minimized tilt and spherical aberrations, respectively.

Second, we checked the conjugation of the SLM plane to the sample, important to achieve imaging with high axial pattern modulation. To optimize this, we acquired a 4-μm 3D SIM stack on a single layer of 100 nm fluorescent beads with axial step size 20-40 nm, much smaller than the Nyquist sampling employed when taking biological data. We then used the ‘Illumination Pattern Focus’ calibration tool in *SIMcheck*^24^ to produce a projection of the axial cross-section of the beads layer along each direction (**Supplementary Fig. 15**). When the SLM surface is well aligned to the sample plane, axial views of beads at each direction appear symmetric, with approximately equal energy appearing above and below the central intensity maxima. Otherwise, axially projected images appear as a ‘zipper-like’ double layer, which indicates the detection camera is imperfectly positioned and should be translated axially. For our system, the SLM position and its corresponding axial interference pattern can be optimized only for a single illumination wavelength at once. We prioritized the 488 nm illumination wavelength and corresponding GFP emission band, leaving the camera position unchanged during 2-color 3D SIM imaging. However, as indicated below, we did translate the camera for 4-beam SIM, 2-color acquisition.

Finally, we evaluated the overall performance of the 3D SIM system by imaging 100 nm yellow green beads and passing the images through *SIMcheck* (**Supplementary Fig. 8**). The ‘Raw Fourier Projection’ output was used to check that the first-order illumination pattern components were evident in the frequency domain (**Supplementary Fig. 8a**). The **‘**Motion & Illumination Variation’ output was used to verify that the system remained stable to within the tolerance required for 3D SIM imaging (**Supplementary Fig. 8b**). A high ‘Modulation Contrast-to-Noise’ value demonstrated good modulation contrast of the illumination pattern (**Supplementary Fig. 8c**). Statistics from the ‘Reconstructed Intensity Histogram’ indicated a good signal-to-noise ratio of the reconstructed image (**Supplementary Fig. 8e**). Finally, FFTs corresponding to the lateral and axial dimensions of the reconstructed image, given by the **‘**Reconstructed Fourier Plots’ (**Supplementary Fig. 8f**) indicated that imaging performance approached the theoretical resolution limit.

### Mirror for 4-beam SIM, 4-beam SIM illumination pattern

A half-inch dielectric mirror (Thorlabs, BB05-E02) was glued to a lens tube (Thorlabs, SM05L05) with optical cement (Norland Products), and the entire assembly mounted to a home-made adaptor. A piezo tip/tilt scanner (Physik Instrumente, S316.10H) connected to the adaptor via set screws was used to precisely control the axial position of the mirror. In this work, only the Z axis of the piezo scanner was controlled, resulting in pure axial motion. This ‘piezo mirror’ assembly was then mounted on a kinematic mirror mount (Thorlabs, POLARIS-K2T) and a manual translation stage (Newport, 9067) to coarsely adjust the horizontal tilt angle and axial position of the mirror. Finally, the entire setup (**Supplementary Fig. 9a**) was bolted to the microscope stage and mounted opposite the sample, via half-inch posts, post holders and 90-degree right-angle clamps (Thorlabs, TR series, PH2 & RA90). By loosening the thumbscrews of the two post holders to remove the assembly, we switched between 3D SIM and 4-beam SIM mode. We required an imaging chamber with sufficient clear aperture to accommodate the half-inch diameter of the mirror. This motivated our choice of the magnetic imaging chamber (Warner Instruments, QR-40LP) with 19.7 mm aperture.

In creating the 4-beam SIM illumination pattern, we considered two key constraints: i) The mirror should be close enough (< 500 μm) to the coverslip to maintain high coherence between incoming and reflected beams, thereby producing high axial modulation depth. ii) The mirror surface should be perpendicular to the zero-order beam for a purely axial modulation. Fine alignment of the mirror and the reflected beam, necessary to satisfy these constraints, was accomplished using the following protocol: i) Set the SLM to widefield mode (all pixels to 0 phase) and focus on a dense single layer of 100 nm beads (or autofluorescence from a dirty coverslip). ii) Attach the mirror setup to the microscope stage with the piezo scanner set at its mid-point voltage. iii) Move the translation stage to lower the mirror towards the sample until it is submerged in the imaging medium. iv) Carefully fine-tune the manual translation stage attached to the mirror (move in ∼10 μm increments) and check the image of the beads layer. When the mirror hits the cover glass, the image of the beads will become out of focus. At this point, move the mirror in the opposite direction (towards the ceiling) by 300 μm. Now the mirror is coarsely positioned in the correct axial position. v) Acquire a 3D stack of images extending axially over 2 μm, with 20 nm step size. We found that multiple images in the stack provided valuable contextual information, helping us to finely align the system. Manually adjust the knobs on the kinematic mirror mount while monitoring the width and direction of the interference fringes. The closer the mirror is to the correct angle, the wider the fringes become. Perfect 2-beam interference with purely axial modulation is achieved when seeing a flat intensity profile (**Supplementary Fig. 9c, Supplementary Video 1**). vi) Repeat steps iv) and v) as necessary to ensure that the mirror is properly aligned. Finally, lock the position of the translation stage; at this point only small adjustments to step v) are required for same-day alignment when using the same imaging chamber. We note that the ‘side beams’ are also reflected from the mirror but illuminate areas well outside the 4-beam imaging field at the illumination angles and mirror-to-coverslip distance used in this work. Although we cleaned the mirror with ethanol between experiments, we found that we never had to replace it during this study.

### 4-beam SIM alignment

A successful SIM reconstruction requires that the axial phase of the illumination (relative position of the illumination pattern with respect to the detection focal plane) is the same for both PSF/OTF measurement and image acquisition. This is usually not a significant challenge in 3D SIM. However, in 4-beam SIM, the axial phase depends not only on the input illumination, but also sensitively on the relative position of the mirror with respect to the detection focal plane. The maxima in the axial direction of the interference pattern should ideally coincide with the detection focal plane^20^ (and be maintained at this position), typically requiring the position of piezo scanner (mirror) to be adjusted (and maintained) before commencing 4-beam SIM acquisition. We invented a bead-based alignment method to correct phase and drift by using the axial profile of a single 100 nm bead deposited on the coverslip (**Supplementary Fig. 12**): i) In widefield mode, obtain a single image of the sample. It is important that the beads are relatively sparse (e.g., 2 μL methanol solution with 1:20000 dilution of beads). ii) Select one fiducial bead as an approximation to the PSF, ensuring that there are no other beads within 50 pixels of it. Record its highest intensity coordinate (*x*_0_, *y*_0_). ii) If performing the phase alignment for the first time, acquire a 2 μm stack with 20 nm step size. Otherwise, record a 1 μm stack to reduce photobleaching and improve acquisition speed. iii) Crop the stack’s lateral dimensions to 50 × 50 pixels centered at (*x*_0_, *y*_0_) and apply a 2D Gaussian fit on the maximum intensity projection to estimate the PSF center (*x*_1_, *y*_1_) with subpixel precision. iv) Apply 2D Gaussian fits to each lateral plane in the cropped stack with predefined center (*x*_1_, *y*_1_). Compute the full width at half maximum (FWHM) along the x direction *FWHM*(*x*), the y direction *FWHM*(*y*) and their average *FWHM*(*average*) at each axial position in the stack. Interpolate *FWHM*(*average*) values within appropriate intervals (e.g., 300-500 nm for the yellow-green channel, 400-600 nm for the red channel) with 100 evenly spaced points, and then compute the averaged axial positions from the 100 points, yielding ‘PSF offset’. v) Derive the axial intensity profile of the bead by summing the intensity in each plane over a circular area centered at (*x*_1_, *y*_1_) with a radius of 3 pixels. Fit the axial intensity profile with a 1D Gaussian function to obtain the peak position of the axial interference pattern closest to the ‘PSF offset’, i.e., ‘SW peak’. vi) Compute the difference in position between ‘PSF offset’ and ‘SW peak’. If the difference is larger than a set tolerance (e.g., 10 nm), the difference is minimized using feedback, by adding ‘PSF offset’ and ‘SW peak’ to the current position of the piezo stage (moving the sample) and piezo scanner (moving the mirror), respectively. Steps ii) to vi) are repeated until the desired tolerance is achieved. Once the difference between ‘PSF offset’ and ‘SW peak’ is within the set tolerance, the axial intensity profile should exhibit side lobes of equal height above and below focus (**Supplementary Fig. 11b**) and 4-beam SIM imaging can commence using the current piezo scanner and piezo stage settings. After moving the stage to image another FOV, we usually waited ∼5 minutes before commencing imaging, finding that this settling time helped to stabilize the system.

As discussed above, the axial interference pattern of 3D SIM can be optimized only for a single illumination wavelength. If the camera position (detector plane) is adjusted correctly for the 488 nm illumination, the corresponding order 1 OTF fully overlaps in 4-beam SIM, which is sufficient for reconstruction (**Supplementary Fig. 15a**). However, when imaging red fluorescent beads excited with 561 nm illumination, gaps appear in the order 1 OTF if the detector plane is not moved (**Supplementary Fig. 15b**). Such missing frequency components cannot be restored during Wiener reconstruction. After translating the camera to the optimal position for 561 nm illumination, OTF overlap is restored (**Supplementary Fig. 15c**). Thus, to achieve acceptable 2-color 4-beam SIM reconstructions, we recorded the optimal positions for 488 nm and 561 nm illumination wavelengths and switched the camera positions between color acquisitions.

### 3- and 4-beam SIM data acquisition and OTF generation

Raw 3D SIM and 4-beam SIM data were collected by applying patterned illumination with five phases (2π/5 relative spacing) and three orientations (60° apart) for a total of 15 images per plane before changing focus (125 nm step size for 3D SIM, 60 nm step size for 4-beam SIM). Each image stack was thus saved in XYPAZ format, where ‘XY’, ‘P’, ‘A’ and ‘Z’ denote lateral plane, phase, orientation, and depth, respectively.

PSF data was acquired similarly except that only one pattern orientation (the first orientation) was acquired when imaging a single 100 nm fluorescent bead. For 4-beam SIM PSF measurement, phase alignment and drift correction were also applied immediately before acquisition. The axial range of the acquired PSF was 8 μm (4 μm below and above the bead center) and the image was cropped to a 256 × 256-pixel region centered on the bead. The OTFs required for parameter estimation and Wiener reconstruction were derived using the following procedure: i) Sum all five image volumes (corresponding to the five phases) to obtain an estimate of the widefield PSF. Use 3-point parabolic fitting around the pixel with maximum intensity to estimate the lateral center of the PSF more precisely. To estimate the axial center of the PSF, apply another 3-point (3D SIM) or 5-point (4-beam SIM) parabolic fitting on the axial view. ii) Subtract camera background, soften edges by multiplying each of the five image volumes with a squared sine function and convert them to the frequency domain by computing the 3D FFT. Multiply the FFT results with a separation matrix to obtain the real and imaginary parts of the five information bands. Multiply the bands with a phase factor determined by the PSF center to shift the PSF to the origin. iii) Divide the bands with the Fourier transform of a solid sphere with a diameter of the bead (100 nm) to compensate for the finite bead size. iv) Rotationally average along *k*_*z*_ axis to reduce noise, converting the bands into 2D data (*k*_*r*_/*k*_*z*_ view). v) The OTF support of the widefield microscope is a toroid with extent determined by the detection NA and emission wavelength. All values outside the OTF support were set to 0 to eliminate noise. In 3D SIM, order 0 and 2 OTFs are identical to the widefield OTF, but the order 1 OTF is composed of two overlapping widefield OTFs. In 4-beam SIM, only the order 2 OTF is equivalent to the widefield OTF, as the order 0 OTF consists of three separated widefield OTFs and the order 1 OTF consists of four overlapping widefield OTFs (**Supplementary Fig. 15**). vi) Normalize all OTFs to the maximum value (DC coordinate) of the order 0 OTF, setting this value to 1. OTFs must be measured for each objective and each laser wavelength, and can be subsequently applied to reconstruct SIM data acquired under the same conditions. When performing SIM reconstruction (see following section), the 2D OTFs must be converted back into a 3D form. We established this correspondence by assigning each voxel coordinate (*k*_*x*_, *k*_*y*_, *k*_*z*_) in the desired 3D OTF to a 2D coordinate 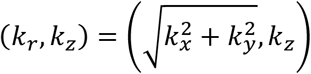, obtaining the value at each 2D coordinate by interpolating the values from the four nearest pixels. Code for deriving OTFs is available as **Supplementary Software**.

### 3- and 4-beam SIM image reconstruction

Raw 3D SIM and 4-beam SIM data were processed with custom-developed MATLAB (Mathworks, R2021b) software (available as **Supplementary Software**). We note that 4-beam SIM reconstruction is fundamentally no different than 3D SIM. Although additional axial information is present in 4-beam SIM data, such data contains the same number of lateral illumination frequency components and thus lateral information bands as in 3D SIM. The reconstruction algorithm we use is thus based on traditional 3D SIM reconstruction^1^, and can be subdivided as follows: i) Preprocessing. A constant camera background was first subtracted from the raw data, and the resulting image edges were softened by multiplying the images with a squared sine function. All images were normalized to have the same total intensity to compensate for fluctuations in illumination or bleaching between the different illumination phases, illumination orientations, or sample depths. ii) Frequency unmixing. For each orientation, five information bands (corresponding to m = 0, ±1, ±2 lateral illumination frequency components) can be estimated and separated based on the 3D FFTs of the five input volumes. iii) Parameter estimation. The precise position of the frequency components (corresponding to pattern line spacing and angle), the initial phase, and the corresponding modulation depth of order 2 (the highest order) were determined by computing the cross-correlation between the shifted order 2 and order 0 (DC) bands. The modulation depth is a useful indicator of cross-correlation performance as line spacing and angle are finely varied to find their optimal values. Good parameter estimation occurs when the modulation depth reaches a local maximum, maximizing the overlap between order 0 and the shifted order 2 information bands. Half the value of the order 2 illumination frequency component position was used for positioning order 1. The estimated illumination frequency component of order 1 was then used to determine the initial phase and modulation depth of order 1. iv) Wiener reconstruction. A generalized Wiener filter incorporating all estimated parameters shifted the bands to their correct positions in frequency space and combined all three orientations to fill in the ‘missing cone’ and enhance resolution. Using a Wiener parameter that is too high reduces resolution, while using a Wiener parameter that is too low artificially increases noise. A value of 0.001, used throughout this work, was empirically found to give a good balance between resolution and noise. After multiplying the generalized Wiener filter result with a triangular apodization function to suppress high-frequency edge artifacts, the final SIM images were derived by computing the inverse FFT and keeping non-negative real values.

### Hardware control

A PC computer (@Xi workstation, Intel Xeon CPU E5-1660 v4 @ 3.20 GHz, 16 threads, 64 GB memory, and 1 TB M.2 NVME SSD) was used to issue commands (waveforms) to a NI-DAQ card (National Instruments, PXI 6733, BNC 2110) housed in an external data acquisition (DAQ) card chassis (National Instruments, PXIe-1073), the internal pco.edge camera acquisition card and the internal SLM control card (**Supplementary Fig. 5**). A schematic describing the main waveforms (and their relative timing) is shown in **Supplementary Fig. 6**. The camera was set to external trigger with rolling shutter mode, receiving rising edges from a digital output (DO) on the DAQ card. The SLM was also set to external trigger mode, receiving falling edges from another DO port to change patterns in a predefined sequence. Stepwise waveforms from an analog output (AO) were used to drive the piezo stage for volumetric acquisition. Three AO ports (488 nm, 561 nm and blanking) drove the AOTF to individually control the intensity of each illumination wavelength. Three different voltages from an AO port were supplied to the LCPR to maximize modulation depth for the three pattern orientations. In 4-beam SIM, the piezo mirror position was controlled by an additional AO port during phase alignment and remained unchanged during subsequent 4-beam SIM acquisition. The x and y positions of the ASI stage, the z position of the ASI objective stepper motor (for coarse axial objective positioning) and the filter wheel selection were controlled through serial port commands.

Control software (Python 3.7) was based on our previous iSIM control software^60^, but modified significantly to control the SLM, LCPR and piezo scanner necessary for 3D SIM and 4-beam SIM. Timing diagrams (**Supplementary Figs. 6, 7**) show key use cases, including PSF/OTF data acquisition (**Supplementary Fig. 6a**), 3D SIM and 4-beam SIM acquisition at one lateral plane (**Supplementary Fig. 6b**), volumetric acquisition (**Supplementary Fig. 7a**) and axial alignment between illumination pattern and the detection focal plane (**Supplementary Fig. 7b**).

Volumetric acquisition time was determined by the laser exposure time per image, number of rows of pixels in the image, camera readout time (delay time between external trigger and laser exposure), SLM loading time, LCPR switching time and the number of z slices. The laser exposure time was set to 20 ms (50 ms in **Fig. 4e-k** and **Supplementary Fig. 19**), the raw image size was 1280 × 1080 pixels, camera readout time was 5.27 ms, SLM loading time was 5 ms and LCPR switching time was 10 ms. Thus, ∼25 ms is required per phase (single image), ∼130 ms for one orientation (5 images), and ∼390 ms for one plane (3 orientations x 5 phases = 15 images). For example, in the fixed U2OS cell imaging shown in **Fig. 2e-g**, when setting the z step size at 0.125 μm for 3D SIM and 0.06 μm z step size for 4-beam SIM, we collected a volume spanning 90 μm x 76 μm x 4 μm in ∼12.7 s and ∼26.5 s, respectively. We also employed a technique based on the ‘Transient Nematic Effect’ (T.N.E.) to shorten the temporal response of the LCPR by ∼2-fold. When changing the LCPR polarization state from low to high voltage, a very short duration (2 ms) high voltage spike is used to accelerate the molecular alignment parallel to the applied field. Voltage is then reduced to achieve the desired polarization (**Supplementary Fig. 6b**) in the remaining 8 ms of the 10 ms total switching time. Similarly, when changing from high to low voltage, a zero-voltage spike (2 ms) is applied before targeting the desired voltage (**Supplementary Fig.7a**). For two-color imaging with only one time point, 561 nm always preceded 488 nm to reduce photobleaching. Time-lapse imaging consisted of a series of repetitions of single-volume imaging with or without a delay between time points. **Supplementary Table 4** details the volume size and corresponding acquisition time for all the data presented in the paper.

### Registration, bleach correction of volumetric time-lapse data

The output values after deep learning are floating-point numbers between 0 and 1. To guarantee that all volumes have similar background intensity values for supplementary videos, outputs were converted to 16-bit unsigned integers, and then bleach corrected using the ImageJ plugin Bleach Correction (Image → Adjust → Bleach Correction → Histogram Matching). This plugin is available at https://imagej.net/plugins/bleach-correction. Adjacent time points were registered using a GPU-based 3D affine registration method^48^, available at https://github.com/eguomin/regDeconProject/tree/master/RegistrationFusion. This registration method is written in C++/CUDA to generate a DLL file, which is then called in MATLAB. The “translation only” registration mode was used.

### Segmentation and tracking in live microtubule data

To track filaments, raw image data was first manually segmented in Aivia 10.2 (Leica-Microsystems) to create mask channels. Mask channels were incorporated as a separate channel in Imaris 9.8.2 (Bitplane) and then converted to surface objects and overlaid over the input data.

Microtubule buckling and centrosome movements were manually tracked in Imaris by placing spots at the desired locations, with the position of the centrosome identified as the point of convergence of microtubule filaments.

### Simulation of mixed structures for axial resolution enhancement

We simulated 3D images (50 pairs for training, and 10 for testing) with mixed structures of dots, lines, and hollow spheres for validating the six-direction deep learning method that yields isotropic resolution from raw 3D SIM input (**Supplementary Fig. 16a**). Simulated structures were created in MATLAB (Mathworks, R2019b, with the Imaging Processing Toolbox). Each volume was composed of 3000 dots, 1800 lines and 600 hollow spheres, randomly located in a 300 × 300 × 300 grid (assuming a pixel size of 40 nm). Dots were generated with random intensity (2000-7000 counts); lines were generated with random angles in 3D (1-360 degree relative to axial axis, and 1-360 degree in lateral planes), random lengths (1-72 pixels), and a random intensity (100-500 counts); and hollow spheres were generated with random inner diameter (2-20 pixels), random thickness (1-2 pixels) and random intensity (10-200 counts). Structures were blurred with different 3D Gaussian functions. In the first dataset, sigma values were 1.3, 1.3, and 3.7 pixels in x, y, z dimensions, respectively, which represented the raw 3D SIM volumes with a resolution of 125 × 125 × 350 nm^3^ (FWHM value), assuming a pixel size of 40 nm. In the second set, sigma values were all 1.3 pixels, serving as the ground truth dataset with an isotropic resolution of 125 × 125 × 125 nm^3^ for the calculation of SSIM and PSNR.

For the six-direction deep learning method, we generated two sets of volumes based on the first dataset (i.e., the simulated raw 3D SIM input), one blurred with a 2D Gaussian function at each xz slice (sigma = 3.6 pixels in x dimension, and 0.5 pixels in z dimension), serving as the ‘input’ in the training; the other blurred with a 1D Gaussian function (sigma = 0.5 pixel) along the z dimension, serving as the ‘ground truth’ in the training. To study the effects of axial down-sampling factors (**Supplementary Fig. 16b**), we extracted xy planes from the first dataset with an axial interval of 2, 4, 6, and 8 (i.e., down-sampling factor) to create new volumes with 300 × 300 × 150 voxels, 300 × 300 × 75 voxels, 300 × 300 × 50 voxels, and 300 × 300 × 37 voxels, respectively. For each down-sampling factor, we interpolated the volumes back to 300 × 300 × 300 voxels and repeated the same Gaussian blurring process as for the first dataset (i.e., without any down-sampling) to generate ‘input’ and ‘ground truth’ datasets in the training.

For the prior isotropization method^35^, we created a 2D PSF by Gaussian blurring a dot along the y dimension (sigma = 3.6 pixels) and fed this into the content aware image restoration (CARE^35^) network pipeline, blurring the lateral views to resemble the lower resolution axial views and learning to reverse this blur to achieve isotropic resolution.

### Quantitative analysis

For the simulated datasets (**Supplementary Fig. 16a**), we selected 10 volumes to evaluate the structural similarity index measure (SSIM) and peak signal to noise ratio (PSNR) of the results we obtained using our six-direction deep learning method vs. the ground truth with MATLAB (Mathworks, R2019b), and then computed the mean value and standard deviation of these volumes (**Supplementary Fig. 16b**).

Lateral and axial resolution measures derived from fluorescent beads (**Fig. 1g, Supplementary Fig. 10d**) were estimated by computing the FWHM of line intensity profiles along xy and xz views. Statistical results (mean ± standard deviation) were obtained from *N* = 102, 100, 99 beads (1.35 NA) and 85, 81, 78 beads (1.27 NA) for widefield microscopy, 3D SIM, and 4-beam SIM, respectively. For rough resolution comparisons, we plotted the Fourier transforms of the images (**Fig. 1e, insets, Supplementary Fig. 10b, insets**).

### Neural networks for denoising and axial resolution improvement

We adapted and developed five neural networks including (1) content aware image restoration (CARE^35^) network that provides isotropic output (i.e., prior isotropic method); (2) six-direction deep learning method that improves axial resolution, which was implemented with the CARE network; (3) two-step 3D residual channel attention network (RCAN), i.e., first denoising raw, low SNR 3D SIM volumes, then predicting 3D SIM from the Wiener reconstruction of the denoised 3D SIM volumes (**Fig. 5a, Supplementary Fig. 20a**); (4) 15 input 3D RCAN that can directly predict a 3D SIM reconstruction from 15 raw low SNR 3D SIM volumes (**Supplementary Fig. 21b**); (5) 15 input DenseDeconNet for predicting 3D SIM reconstructions from 15 low SNR 3D SIM volumes (**Supplementary Fig. 21b**). Most model training and applications were performed on an NVIDIA TITAN RTX GPU (with 24 GB memory) installed on a local workstation. We also performed some CARE and RCAN training and model applications in Amazon Web Services (AWS), using a virtual machine with four NVIDIA Tesla V100 GPUs (each with 16 GB memory). We used Python version 3.7.1 for all neural networks, and Tensorflow framework version 2.4.1, 1.13.1, 1.14.0 for CARE, RCAN and DenseDeconNet, respectively.

CARE software was installed from GitHub (https://github.com/CSBDeep/CSBDeep). For simulated data training (**Supplementary Fig. 16a**), patches of size 64 × 64 × 64 voxels were randomly cropped from 50 3D volumes with size 300 × 300 × 300 voxels. For other 3D datasets (**Figs. 4-6, Supplementary Figs. 16-23**), 3D SIM volumes were first interpolated to 50 nm pixel size in all three dimensions. When using the prior method (**Supplementary Fig. 16**), the interpolated volumes, a 2D PSF (consisting of a point blurred with a 1D Gaussian function, sigma = 2.8 pixels along the y dimension), and the axial downsampling factor were fed into the isotropic CARE network using a patch size of 64 × 64 to create training pairs.

For the six-direction deep learning method, interpolated volumes were (1) blurred with a 1D Gaussian function (sigma = 1.0 pixel) along the z dimension to remove spurious sidelobes in the Fourier domain due to axial interpolation, serving as the ‘ground truth’ in the training or (2) blurred with a 2D Gaussian function at each xz plane (sigma = 2.8 pixels in the x dimension, degrading lateral resolution to the extent of the axial resolution, and 1.0 pixels in the z dimension as for the ground truth); downsampled along the x dimension to mimic the coarse axial sampling in real experiments; and finally upsampled to recover an isotropic pixel size, serving as the ‘input’ for training (**Supplementary Fig. 17a**). Patches of size 64 × 64 × nz (nz is the number of planes) were randomly cropped from the data, and 10% of the patches set aside for validation.

The trained network can recover resolution along the x axis from unseen ‘test’ data (**Supplementary Fig. 17b**), which was degraded with the same procedure (i.e., blurring, downsampling, and upsampling) as the ‘input’ data used to train the network. Digitally rotating the degraded data about the y axis and passing it through the trained network results in improved resolution along the lateral direction in the rotated space. By rotating the data back to the original frame, resolution can thus be improved along an arbitrary axis (**Supplementary Fig. 17c**). Last, by recording the maximum value at each spatial frequency taken over all rotations, a final prediction with isotropic resolution is obtained (**Supplementary Fig. 17d**).

Naïve rotation and interpolation in the xz plane will dramatically enlarge the size of each volume. For example, for raw input data with large xz aspect ratio spanning 800 × 800 × 80 voxels, as is typical for imaging single cells, a 60-degree rotation in the xz plane will produce a volume 800 × 800 × 733 voxels in size. This large size results in inefficient network prediction (for this example the prediction is useless over a region spanning 800 × 800 × 653 voxels). We thus cropped the input data into multiple subvolumes (e.g., 10 subvolumes of 80 × 800 × 80 voxels) prior to application of the CARE network. After rotation and interpolation, each subvolume spanned only 110 × 800 × 110 voxels, resulting in useless prediction over only 30 × 800 × 30 voxels, a 65-fold improvement in efficiency compared to the non-cropped case. This processing pipeline was implemented in Python v 3.7.1: (1) automatically cropping raw input data to multiple subvolumes with identical x and z dimensions, with 10 pixel overlap along the x axis; (2) rotating and bilinearly interpolating each subvolume with the built-in SciPy function ndimage.rotate; (3) applying the neural network model to each subvolume; (4) rotating back the subvolume predictions and cropping them to their original sizes; (5) stitching the subvolumes by averaging the overlap regions; (6) Fourier transforming the stitched volume for each rotation angle; (7) calculating the maximum value at each spatial frequency taken over all rotations; (8) inverse Fourier transforming the result and taking the absolute values as the final output.

All training for the prior method and six-direction method used 50 volumes without data augmentation (rotation, translation), and testing varied from 10-100 volumes (**Figs. 4-6, Supplementary Figs. 16, 18, 19, 20, 23**). The training learning rate was 2 × 10^4^, the number of epochs 100, the number of steps per epoch 200, and mean absolute error (MAE) was used as loss function. The training time for each model varied from ∼3− 7 hours. For example, it took ∼3 h for training the Jurkat T cells expressing EMTB 3x GFP datasets (**Fig. 6, Supplementary Videos. 13-16**), and ∼5.5 hours to apply the model to recover a 100 time-point dataset with size 420 × 420 × 80 voxels and 6 directions (total 600 volumes), including the processing time for cropping, image rotation, network prediction, rotation back, stitching, combination with max frequency method, and file reading/writing.

For the two-step denoising studies employing RCAN (**Figs. 5, 6, Supplementary Figs. 20-23**), we used our recently developed 3D RCAN model, appropriate for restoring image volumes (https://github.com/AiviaCommunity/3D-RCAN)^37^. For all 3D data, training patches of size 128 × 128 × nz (nz is the number of acquisition planes without axial interpolation) were randomly cropped from the data. To train the first denoising network, we used patches derived from 750 matched volumes (50 acquisitions x 3 orientations x 5 phases) acquired at low and high illumination intensity (e.g., ∼0.8 W/cm^2^ vs. 8 W/cm^2^ for **Supplementary Fig. 20**). The trained network was used to denoise raw, low SNR data (5 phases x 3 orientations), and the denoised output used for 3D SIM reconstructions as described above. To train the second denoising network, we used patches derived from 50 matched volumes (3D SIM reconstructions derived from high SNR raw data as high SNR ‘ground truth’; and the 3D SIM reconstructions derived from the denoised data as the matched ‘input’ data). The output of this second model may then be axially interpolated, blurred, downsampled, upsampled, and fed into the six-direction model that enhances axial resolution, generating a final denoised reconstruction with isotropic resolution (**Figs. 5, 6, Supplementary Figs. 20, 23**). For training the RCAN networks, the learning rate was 2 × 10^−4^, the number of epochs for training 200, the number of residual blocks 5, the number of residual groups 5, the number of channels 32, the steps per epoch 400, and mean absolute error (MAE) was used as the loss function.

For some studies (**Supplementary Figs. 21, 22**), we extended our 3D RCAN method to handle 15 input volumes. Multiple volumetric inputs with the same shape were concatenated into a single volume with multiple channels and fed to the model. We followed ref.^61^ and used subpixel convolution^62^ to generate up-sampled super-resolution outputs. A fixed number of volumetric inputs are paired with one ground truth volume for model training and the same number of volumetric inputs are expected for prediction. For this multiple input training, we used the same network parameters as in single-input RCAN training.

For the studies using DenseDeconNet (**Supplementary Fig. 21**), we similarly extended our previously published single-input neural network (https://github.com/eguomin/regDeconProject/tree/master/DeepLearning) for 15-inputs. The input data was 15 raw low SNR 3D SIM volumes, and the ground-truth data consisted of high SNR 3D SIM reconstructions. For all 3D data, patches of size 64 × 64 × 64 were randomly cropped from the data. All 15 low SNR data were concatenated along the channel-axis to form the input. The percentile-based normalization was adopted to normalize the input and ground-truth (the low percentile was 1.0 and the high percentile was 99.8). In the training, the objective function incorporated three terms: the mean square error (MSE), the structural similarity (SSIM) index, and the minimum value of the output (MIN); the parameter to control the MIN term was set to 1. The training epoch was 200 with 400 steps per epoch, the learning rate was 0.01, decay rate was 0.985, decay step was 400.

### Sample preparation

#### General considerations

Coverslips were cleaned and coated with Poly-L-Lysine and (particularly for 4-beam SIM) fluorescent beads. Unless otherwise noted, we used the following protocol. High precision, #1.5 coverslips (Thorlabs, CG15XH) were cleaned by immersion in 75% ethanol overnight and air dried before use. Approximately 50 μL of Poly-L-lysine solution (Sigma, P8920) was applied to the center of the coverslips in a biosafety cabinet. After air drying for 15 minutes at room temperature (RT), coverslips were rinsed in pure ethanol and air dried until use. Orange, red, or yellow-green FluoSpheres (Invitrogen, F8800, F8801, F8803, all 0.1 μm diameter) were selected depending on the application, dissolved in pure methanol at 1:20000 dilution, and 2 μL of the solution applied to the center of a poly-L-lysine coated coverslip.

For 4-beam SIM experiments using the 1.35 NA silicone oil lens, we performed imaging in an iodixanol solution index-matched to the refractive index of the silicone oil (1.406) to minimize aberrations. Iodixanol solution (Sigma, D1556) consisted of 45.6% iodixanol in water. We verified its refractive index as 1.406 using a refractometer (American Optical).

#### Bacteria

Vegetatively growing *Bacillus subtilis* strain PY79^63^ was grown in Luria-Bertani Broth (KD Medical, BLE-3030) for 2 h at 37 °C, shaking at 250 rpm. To visualize localization of DivIVA-GFP, strain KR541 (*amyE::P*_*hyperspank*_*-divIVA-gfp cat*)^64^ was grown in casein hydrolysate media^65^ (KD Medical, CUS-0803) containing 1 mM final concentration of IPTG to induce DivIVA-GFP production for 2 h at 37 °C, shaking at 250 rpm. To induce sporulation by the resuspension method, an overnight culture of strain CVO1000 (*amyE::spoVM-gfp cat*)^66^ grown at 22 °C was first sub-cultured to a final OD_600nm_ of 0.1 in casein hydrolysate media for 2 h at 37 °C, shaking at 250 rpm. Cells were harvested by centrifugation at 14,000 × g, resuspended in equal volume of Sterlini-Mandelstam media^65^ (KD Medical, CUS-0822), and grown for 4 h at 37 °C, shaking at 250 rpm. Prior to imaging, 1 ml of culture was removed, centrifuged at 14,000 × g, and the cell pellet resuspended in PBS.

For some experiments, we stained bacterial membranes. Bacteria were diluted in 1mL of PBS and stained with CellBrite Fix 488 (Biotium, 30090-T) or CellBrite Fix 555 (Biotium, 30088-T) for 5 minutes at room temperature (RT) at 1X working concentration according to the manufacturer’s guidelines. Stained bacteria were washed 3 times with 1X PBS, each time centrifuging the solution at 3000 RPM for 3 minutes. Finally, bacteria were concentrated in 100 μL of 1X PBS and 2 μL placed on the center of a Poly-L-lysine and beaded coated coverslip for imaging.

#### Mouse LSECs

C57BL/6 Mouse Primary Liver Sinusoidal Endothelial Cells (Cell Biologics, C57-6017) were cultured in culture medium (Complete Mouse Endothelial Cell Medium, containing 10% Fetal Bovine Serum, and Endothelial Cell Growth Supplement (Cell Biologics, M1168, 6912, and 1166 respectively) according to the manufacturer’s instructions. Prior to cell deposition, glass coverslips (Thorlabs, CG15XH) were sequentially washed with 0.1 M sodium hydroxide, 0.1 M hydrochloric acid, and acetone, coated with 200 μl Poly-L-lysine solution (Sigma, Cat. P8920), allowed to dry for 5 minutes, and washed once with pure methanol. Equal volumes of red (Invitrogen, F8801) and yellow-green (Invitrogen, F8803) fluorescent microspheres were diluted 1:20000 in methanol, and 2 μL of the bead dilution was added to the center of the coverslip. After the methanol dried, the coverslips were placed in an aqueous solution containing 1% rat tail collagen (Cell Biologics, 6953) for 1 hour and placed in culture medium. After counting, 50,000 cells were seeded per 3.5 cm dish and incubated at 37 °C in 5% CO_2_ for 16 hours. Cultured cells were fixed with 4% paraformaldehyde in PBS for 20 minutes, washed in PBS, incubated with Alexa Fluor™ 568 Phalloidin (Thermofisher, A12380, 7unit/1ml) in PBS for 2h at room temperature, washed in PBS, fixed again in 4% paraformaldehyde, washed with water, stained with CellBrite Fix 488 (Biotium, 30090-T, 1:250 dilution) in PBS for 10 minutes at room temperature, and finally washed and stored in PBS prior to imaging. Imaging was performed within six hours of staining.

#### Mouse embryonic fibroblasts

Mouse embryonic fibroblasts (MEFs, gift of Oliver Daumke’s lab) were cultivated in DMEM (Gibco, 119950409) supplemented with 10% fetal bovine serum (Atlanta Biologicals, S10350) and 1% Penicillin-Streptomycin (Gibco, 15070063) at 37 °C in 5% CO_2_. For imaging experiments the cells were seeded on fibronectin (Sigma, F1141) coated glass coverslips (Thorlabs, CG15XH), and transfected with Caveolin1-EGFP plasmid (gift of Richard Lundmark) using Lipofectamine3000 (Invitrogen, L3000-001). After 48 h the cells were washed with PBS (Gibco, 10010023) followed by fixation with 4% paraformaldehyde/PBS (Electron Microscopy Science, 15700) for 20 min. Next, the cells were washed three times with PBS, treated with 3% bovine serum albumin (BSA, Sigma, A9418)/ 0.1% Triton X-100 (Sigma, X100)/PBS for 30 min, and followed by incubation with 3% BSA/PBS for 45 min. Caveolae were stained with rabbit polyclonal antibody against Cavin1 (abcam, 76919, 1:100 in 3% BSA/PBS) for 1 h. Afterwards, MEFs were washed extensively and GFP-booster (tagged with Alexa488, ChromoTek, gb2AF488-50, 1:500 in 3%BSA/PBS) and anti-rabbit-Alexa568 (Invitrogen, A11011, 1:500) were applied for 1 h. To remove any unbound antibodies cells were washed 5 times with PBS and stored at 4 °C in PBS until imaging.

#### Jurkat T cells

E6-1 Jurkat cells were cultured in RPMI 1640 supplemented with 10% fetal bovine serum (FBS) and 1% Penn-Strep antibiotics. For transient transfections we used the Neon (Thermo Fisher Scientific) electroporation system 2 days before the experiment. 2 × 10^5^ cells were resuspended in 10 μL of R-buffer with 0.5–2 μg of the EMTB 3x EGFP plasmid (a gift from William Bennett, Addgene plasmid 26741). Cells were exposed to three pulses of amplitude 1325 V and duration 10 ms in the electroporator. Cells were then transferred to 500 μL of RPMI 1640 supplemented with 10% FBS and kept in the incubator at 37 °C.

Glass coverslips (Thorlabs, CG15XH) were incubated in poly-l-lysine (PLL, Sigma Aldrich, P8920-100ML) at 0.1% W/V for 10 min. PLL was washed with 70% ethanol and the coverslips left to dry. T-cell activating antibody coating was performed by incubating the coverslips in a 10 μg/ml solution of anti-CD3 antibody (Hit-3a, Thermo Fisher Scientific, 16-0039-85) for 2 h at 37 °C. Excess anti-CD3 was removed by washing with L-15 imaging media (Fisher Scientific, 21-083-027 immediately prior to cell plating.

For fixed cell experiments, EMTB-3xEGFP expressing Jurkat cells were plated on anti-CD3 coated coverslips for 7 minutes in L-15 media. Cells were rinsed with 1X PBS 3 times at room temperature, covered with 100% methanol cooled to -20 C for 3 minutes, and finally washed 3 times with 1X PBS.

For experiments with live cells, a small volume (typically 500 μL) containing 3×10^5^ cells were centrifuged for 5 minutes at 250 RCF. The supernatant was removed, the cells resuspended in 120 μL of L-15 imaging media and approximately 40 μL (equivalently 1×10^5^ cells) of L-15 media with cells was added to the anti-CD3 coated coverslip. Cells were allowed to settle for 2 minutes before commencing imaging.

#### U2OS cells

U2OS cells (ATCC, HTB-96) were cultured in DMEM media (Lonza, 12-604F) supplemented with 10% Fetal Bovine Serum (Thermo Fisher, A4766801) at 37 °C and 5% CO_2_ on Poly-L-lysine and optionally beaded (for 4-beam SIM) coverslips placed in 6 well plates (Corning, 3506).

We used the following buffers for immunolabeling vimentin and microtubules: (i) PEM buffer (in 1X PBS): 80 mM PIPES sodium salt (Sigma, P2949), 5 mM EGTA (Sigma, E3889), 2 mM MgCl_2_ (Quality Biological, 351-033-721), adjusted with sodium hydroxide to pH 6.8. (ii) Pre-extraction solution: PEM buffer supplemented with 0.3% Triton X-100 (Sigma, 93443) and 0.125% Glutaraldehyde (Sigma, G5882). (iii) Microtubule fixation buffer: Pre-extraction solution supplemented with 2% Paraformaldehyde (Electron Microscopy Sciences, 15710).

To immunolabel microtubules, U2OS cells were treated with pre-extraction buffer at 37 °C for 30 s, then the buffer was quickly replaced with microtubule fixation buffer at 37 °C for 15 minutes. Fixed cells were rinsed 3 times with 1 mL of 1X PBS (Gibco, 10010-023), quenched in 1 mL of 0.1% sodium borohydride/PBS solution (Sigma, 213462) for 7 minutes at RT, and blocked in 100% FBS (Sigma, 12103C) at 37°C for 1 hour. Microtubules were incubated with primary mouse-α-tubulin antibody (Invitrogen, 32-2500, 1:100) in 1X PBS supplemented with 10% FBS overnight at 4 °C, washed 3 times (1 minute incubation each time) in 1X PBS, and labeled with secondary donkey-α-mouse Alexa Fluor 488 antibody (Jackson Immuno Research, 715-547-003, 1:200 dilution) in 1X PBS supplemented with 10% FBS for 1 hour at RT.

Vimentin was immunolabeled with primary rabbit-α-vimentin (Abcam, 92547, 1:100) in 1X PBS supplemented with 10% FBS overnight at 4 °C, washed 3 times (1 minute incubation each time), and labeled with secondary donkey-α-mouse Alexa Fluor 594 (Jackson Immuno Research, 711-587-003, 1:200 dilution) in 1X PBS supplemented with 10% FBS for 1 hour at RT. Samples were rinsed 3 times in 1X PBS before imaging. For dual-color samples, we performed microtubule and vimentin immunolabeling in parallel on the same sample.

For immunolabeling Tomm20, U2OS cells were fixed with 2% paraformaldehyde and 0.125% Glutaraldehyde in 1X PBS for 15 min at RT. Cells were rinsed 3 times with 1X PBS, permeabilized by 0.1% Triton X-100/PBS (Sigma, 93443) for 1 min at RT, rinsed 3 times with 1X PBS, incubated with primary Rabbit-α-Tomm20 (Abcam, ab186735, 1:100 dilution in 1X PBS) for 1 hour at RT, washed in 1X PBS (1 minute incubation each time) 3 times, stained in with secondary antibody Donkey-α-Rabbit Alexa 488 (Jackson Immuno Research, 711-547-003, 1:200 in 1X PBS) for 1 hour at RT, and finally washed 3 times (1 minute incubation each time) prior to imaging.

To label actin, U2OS cells were similarly fixed, permeabilized, and washed, then incubated with Phalloidin Alexa Fluor 488 (Invitrogen, 2090563, 1:50 dilution in 1X PBS) or Phalloidin Alexa Fluor 568 (Invitrogen, 1800130, 1:50 dilution) for 1 hour at RT, rinsed 3 times with 1X PBS, and incubated in 1X PBS or Iodixanol solution before imaging.

We stained mitochondria and internal membranes with synthetic dyes for live cell imaging. In the former case, U2OS cells were incubated with MitoTracker Green FM (Invitrogen, M7514, 100 nM in 1X PBS) for 15 minutes at 37 °C and rinsed 3 times with 1X PBS immediately prior to imaging. In the latter case, U2OS cells were incubated in DMEM media with Potomac Gold (kindly provided by Luke Lavis, 200 nM in DMEM media supplemented with 10% FBS) for 30 minutes at 37 °C. Immediately prior to imaging, cells were rinsed 3 times with 1X PBS.

We also transfected cells with plasmids to mark organelles for live imaging experiments. Cell cultures were transfected using xTreme gene HP DNA Transfection Reagent (Sigma, 6366236001). The transfection mixture contained 100 μL 1X PBS, 2 μL Transfection Reagent, and 1 μg plasmid DNA. To label outer mitochondrial membranes, cell cultures were transfected at 50% confluency with mEmerald-Tomm20-C-10 plasmid DNA (Addgene, 54281), and imaged one day after transfection. We also created a stable cell line to mark lysosomes. Cells were transfected with LAMP1-EGFP plasmid DNA (gift of George Patterson’s lab) at 80-85% confluency. The next day, cells were incubated with fresh media for another 24 hours. Transfected cell cultures were screened by G418 selection agent (Corning, 30-234-CR, 750 μg/mL) for 2 weeks. After screening, GFP positive cells were sorted into a 48-well plate automatically (BD, FACS Aria III) so that each well contained only one cell. Sorted cells were cultured with 750 μg/mL G418 for further selection. Clones that managed to survive and expand were transferred to another dish for further expansion, and a small aliquot used to check GFP signal and cell morphology.

For some dual-color experiments, U2OS cells expressing LAMP1-EGFP were incubated in 1X PBS with Lysotracker DND99 (Invitrogen, L7528, 50 nM) for 5 minutes at 37 °C. Prior to imaging, cells were rinsed 3 times with 1X PBS.

## Software availability

The custom software used in this study are available upon request, with most software and test data publicly available https://github.com/eexuesong/SIMreconProject; https://github.com/CSBDeep/CSBDeep and https://github.com/AiviaCommunity/3D-RCAN.

## Data availability

The data that support the findings of this study are included in **Supplementary Figs. 1–23** and **Supplementary Videos 1–16**, and some representative source data for the figures (**Figs. 1d, 2a, 2e, 2h, 3a, 3c, 3h, 3b, 3e, 4b, 4e, 5b, 5e, 6**) will be publicly available on Zenodo after publication. Other datasets are available from the corresponding author upon reasonable request. Source data are provided with this paper.

## References

1 Gustafsson, M. G. L. et al. Three-Dimensional Resolution Doubling in Wide-Field Fluorescence Microscopy by Structured Illumination. Biophys. J. 94, 4957–4970 (2008).

2 Schermelleh, L. et al. Super-resolution microscopy demystified. Nat Cell Biol 21, 72–84 (2019).

3 Shao, L., Kner, P., Rego, E. H. & Gustafsson, M. G. L. Super-resolution 3D microscopy of live whole cells using structured illumination. Nat. Methods 8, 1044–1046 (2011).

4 Fiolka, R., Shao, L., Rego, E. H., Davidson, M. W. & Gustafsson, M. G. L. Time-lapse two-color 3D imaging of live cells with doubled resolution using structured illumination. Proc Natl Acad Sci USA 109, 5311–5315 (2012).

5 Schermelleh, L. et al. Subdiffraction Multicolor Imaging of the Nuclear Periphery with 3D Structured Illumination Microscopy. Science 320, 1332–1336 (2008).

6 Strauss, M. P. et al. 3D-SIM Super Resolution Microscopy Reveals a Bead-Like Arrangement for FtsZ and the Division Machinery: Implications for Triggering Cytokinesis. PLoS Biol 10, e1001389 (2012).

7 Rowlett, V. W. & Margolin, W. 3D-SIM Super-resolution of FtsZ and Its Membrane Tethers in Escherichia coli Cells. Biophysical Journal 107, L17–L20 (2014).

8 Bisson-Filho, A. W. et al. Treadmilling by FtsZ filaments drives peptidoglycan synthesis and bacterial cell division. Science 355, 739–743 (2017).

9 Lesterlin, C., Ball, G., Schermelleh, L. & Sherratt, D. RecA bundles mediate homology pairing between distant sisters during DNA break repair.. Nature 506, 249–253 (2014).

10 Baddeley, D. et al. Measurement of replication structures at the nanometer scale using super-resolution light microscopy. Nucleic Acids Research 38, e8 (2009).

11 Regev-Rudzki, N. et al. Cell-Cell Communication between Malaria-Infected Red Blood Cells via Exosome-like Vesicles. Cell 153, 1120–1133 (2013).

12 Riglar, D. T. et al. Spatial association with PTEX complexes defines regions for effector export into Plasmodium falciparum-infected erythrocytes. Nature Communications 4, 1415 (2013).

13 Kwon, H.-K., Chen, H.-M., Mathis, D. & Benoist, C. Different molecular complexes that mediate transcriptional induction and repression by FoxP3. Nature Immunology 18, 1238–1248 (2017).

14 Rodermund, L. et al. Time-resolved structured illumination microscopy reveals key principles of Xist RNA spreading. Science 372, eabe7500 (2021).

15 Brown, A. C. N. et al. Remodelling of Cortical Actin Where Lytic Granules Dock at Natural Killer Cell Immune Synapses Revealed by Super-Resolution Microscopy. PLoS Biol 9, e1001152 (2011).

16 Wang, C.-J. R., Carlton, P. M., Golubovskaya, I. N. & Cande, W. Z. Interlock Formation and Coiling of Meiotic Chromosome Axes During Synapsis Genetics 183, 905–915 (2009).

17 Cogger, V. C. et al. Three-dimensional structured illumination microscopy of liver sinusoidal endothelial cell fenestrations. J Struct Biol 171, 382–388 (2010).

18 Sonnen, K. F., Schermelleh, L., Leonhardt, H. & Nigg, E. A. 3D-structured illumination microscopy provides novel insight into architecture of human centrosomes. Biol Open 1, 965–976 (2012).

19 Stephan, T. et al. MICOS assembly controls mitochondrial inner membrane remodeling and crista junction redistribution to mediate cristae formation. EMBO J. 39 (2020).

20 Shao, L. et al. I5S: Wide-Field Light Microscopy with 100-nm-Scale Resolution in Three Dimensions. Biophysical Journal 94, 4971–4983 (2008).

21 Manton, J. D., Strohl, F., Fiolka, R., Kaminski, C. F. & Rees, E. J. Concepts for structured illumination microscopy with extended axial resolution through mirrored illumination kBiomedical Optics Express 11, 2098–2108 (2020).

22 Lanni, F. in Applications of Fluorescence in the Biomedical Sciences 505–521 (1986).

23 Bailey, B., Farkas, D. L., Taylor, D. L. & Lanni, F. Enhancement of axial resolution in fluorescence microscopy by standing-wave excitation. Nature 366, 44–48 (1993).

24 Ball, G. et al. SIMcheck: a Toolbox for Successful Super-Resolution Structured Illumination Microscopy. Sci. Reports 5, 15915 (2015).

25 Demmerle, J. et al. Strategic and practical guidelines for successful structured illumination microscopy. Nature Protocols 12, 988–1010 (2017).

26 Boothe, T. et al. A tunable refractive index matching medium for live imaging cells, tissues and model organisms. eLIFE 6, e27240 (2017).

27 Arigovindan, M., Sedat, J. W. & Agard, D. A. Effects of depth dependent spherical aberrations in 3D structured illumination microscopy. Optics Express 20, 6527–6541 (2012).

28 Bratton, B. P. & Shaevitz, J. W. Simple Experimental Methods for Determining the Apparent Focal Shift in a Microscope System. PLoS One 10, e0134616 (2015).

29 Eswaramoorthy, P. et al. Cellular architecture mediates DivIVA ultrastructure and regulates min activity in Bacillus subtilis.. MBio. 2, e00257–00211 (2011).

30 Ramamurthi, K. S., Lecuyer, S., Stone, H. A. & Losick, R. Geometric Cue for Protein Localization in a Bacterium. Science 323, 1354–1357 (2009).

31 Peluso, E. A., Updegrove, T. B., Chen, J., Shroff, H. & Ramamurthi, K. S. A 2-dimensional ratchet model describes assembly initiation of a specialized bacterial cell surface. Proc Natl Acad Sci U S A 116, 21789–21799 (2019).

32 Gan, Z. et al. Vimentin Intermediate Filaments Template Microtubule Networks to Enhance Persistence in Cell Polarity and Directed Migration. Cell Systems 3, 252–263 (2016).

33 Spahn, C. K. et al. A toolbox for multiplexed super-resolution imaging of the E. coli nucleoid and membrane using novel PAINT labels. Sci. Reports 8, 14768 (2018).

34 Laissue, P. P., Alghamdi, R. A., Tomancak, P. & Reynaud, E. G. S. H. Assessing phototoxicity in live fluorescence imaging. Nat Methods 14, 657–661 (2017).

35 Weigert, M. et al. Content-aware image restoration: pushing the limits of fluorescence microscopy. Nat Methods 15, 1090–1097 (2018).

36 Wang, H. et al. Deep learning enables cross-modality super-resolution in fluorescence microscopy. Nat Methods 16, 103–110 (2019).

37 Chen, J. et al. Three-dimensional residual channel attention networks denoise and sharpen fluorescence microscopy image volumes. Nature Methods 18, 678–687, doi:https://doi.org/10.1101/2020.08.27.270439 (2021).

38 Qiao, C. et al. Evaluation and development of deep neural networks for image super-resolution in optical microscopy. Nature Methods 18, 194–202 (2021).

39 Ouyang, W., Aristov, A., Lelek, M., Hao, X. & Zimmer, C. Deep Learning Massively Accelerates Super-Resolution Localization Microscopy. Nature Biotechnol. 36, 460–468 (2018).

40 Weigert, M., Royer, L., Jug, F. & Myers, G. Isotropic reconstruction of 3D fluorescence microscopy images using convolutional neural networks. International Conference on Medical Image Computing and Computer-Assisted Intervention, 126-134 (2017).

41 Wu, Y. et al. Multiview confocal super-resolution microscopy. Nature 600, 279–284, doi:https://doi.org/10.1101/2021.05.21.445200 (2021).

42 Krüger, J.-R., Keller-Findeisen, J., Geisler, C. & Egner, A. Tomographic STED microscopy. Biomed Opt Express 11, 3139–3163 (2020).

43 Matthaeus, C. & Taraska, J. W. Energy and Dynamics of Caveolae Trafficking. Front Cell Dev Biol 8, 614472 (2021).

44 Chu, K. et al. Image reconstruction for structured-illumination microscopy with low signal level. Optics Express 22, 8687–8702 (2014).

45 Jin, L. et al. Deep learning enables structured illumination microscopy with low light levels and enhanced speed. Nat Commun. 11, 1934 (2020).

46 Christensen, C. N., Ward, E. N., Lu, M., Lio, P. & Kaminski, C. F. ML-SIM: universal reconstruction of structured illumination microscopy images using transfer learning. Biomedical Optics Express 12, 2720–2733 (2021).

47 Qiao, C. et al. 3D Structured Illumination Microscopy via Channel Attention Generative Adversarial Network. IEEE Journal of Selected Topics in Quantum Electronics 27, 6801711 (2021).

48 Guo, M. et al. Rapid image deconvolution and multiview fusion for optical microscopy. Nature Biotechnol. 38, 1337–1346, doi:https://doi.org/10.1038/s41587-020-0560-x (2020).

49 Rey-Suarez, I., Rogers, N., Kerr, S., Shroff, H. & Upadhyaya, A. Actomyosin dynamics modulate microtubule deformation and growth during T cell activation. Molecular Biology of the Cell, mbcE20100685 (2021).

50 Murugesan, S. et al. Formin-generated actomyosin arcs propel T cell receptor microcluster movement at the immune synapse. Journal of Cell Biology 215, 383–399 (2016).

51 Yi, J. et al. Centrosome repositioning in T cells is biphasic and driven by microtubule end-on capture-shrinkage. J Cell Biol. 202, 779–792 (2013).

52 Gros, O. J., Damstra, H. G. J., Kapitein, L. C., Akhmanova, A. & Berger, F. Dynein self-organizes while translocating the centrosome in T-cells. Molecular Biology of the Cell 32, 855–868 (2021).

53 Lanni, F. Feedback-stabilized focal plane control for light microscopes. Review of Scientific Instruments 64 (1993).

54 Li, Y. et al. Incorporating the image formation into deep learning improves network performance in deconvolution applications. bioRxiv (2022).

55 Bodén, A. et al. Volumetric live cell imaging with three-dimensional parallelized RESOLFT microscopy. Nature Biotechnology 39, 609–618 (2021).

56 Cao, B., Coelho, S., Li, J., Wang, G. & Pertsinidis, A. Volumetric interferometric lattice light-sheet imaging. Nature Biotechnology 39, 1385–1393 (2021).

57 Li, D. et al. Extended-resolution structured illumination imaging of endocytic and cytoskeletal dynamics. Science 349, aab3500 (2015).

58 Hoffman, D. P. et al. Correlative three-dimensional super-resolution and block-face electron microscopy of whole vitreously frozen cells. Science 367, eaaz5357 (2020).

59 Gustafsson, M. G. L., Agard, D. A. & Sedat, J. W. I5M: 3D widefield light microscopy with better than 100 nm axial resolution. J. Microsc. 195, 10–16 (1999).

60 York, A. G. et al. Instant super-resolution imaging in live cells and embryos via analog image processing. Nat Methods 10, 1122–1126 (2013).

61 Zhang, Y. et al. Image Super-Resolution Using Very Deep Residual Channel Attention Networks. European Conference on Computer Vision, 286–301 (2018).

62 Shi, W. et al. Real-Time Single Image and Video Super-Resolution Using an Efficient Sub-Pixel Convolutional Neural Network. arXiv, 1609.05158 (2016).

63 Youngman, P., Perkins, J. B. & Losick, R. Construction of a cloning site near one end of Tn917 into which foreign DNA may be inserted without affecting transposition in Bacillus subtilis or expression of the transposon-borne erm gene. Plasmid 12, 1–9 (1984).

64 Ramamurthi, K. S. & Losick, R. Negative membrane curvature as a cue for subcellular localization of a bacterial protein. Proc Natl Acad Sci U S A 106, 13541–13545 (2009).

65 Sterlini, J. M. & Mandelstam, J. Commitment to sporulation in Bacillus subtilis and its relationship to development of actinomycin resistance. Biochem. J. 113, 29–37 (1969).

66 van Ooij, C. & Losick, R. Subcellular localization of a small sporulation protein in Bacillus subtilis. J Bacteriol. 185, 1391–1398. (2003).

