## Supplementary Material for "Three-dimensional structured illumination microscopy with enhanced axial resolution"

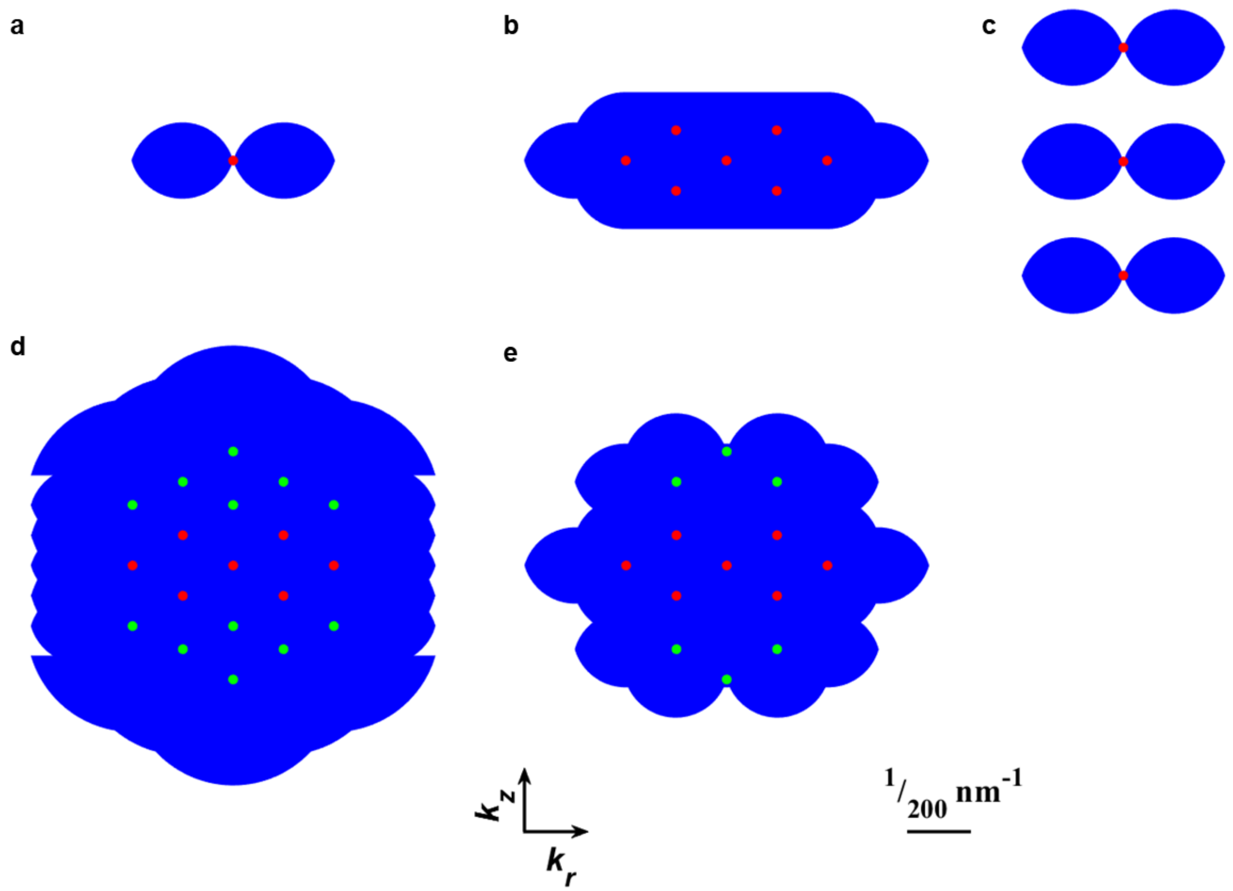


**Supplementary Fig. 1, Support of optical transfer functions assuming a silicone oil objective lens.** Axial cross sections through support are shown for **a**) widefield, **b**) 3D SIM, **c)** standing wave, **d**) $I^5S$, and **e**) 4-beam, mirror-based SIM systems. Dots indicate illumination spatial frequency components, with green dots in **d, e**) indicating additional frequency components that are not present in 3D SIM. Supports were simulated with the following parameters: emission NA = 1.35; refractive index of immersion medium = 1.406; excitation wavelength: 488 nm; emission wavelength: 525 nm; side beams in **b, d, e**) at 92% of pupil radius.

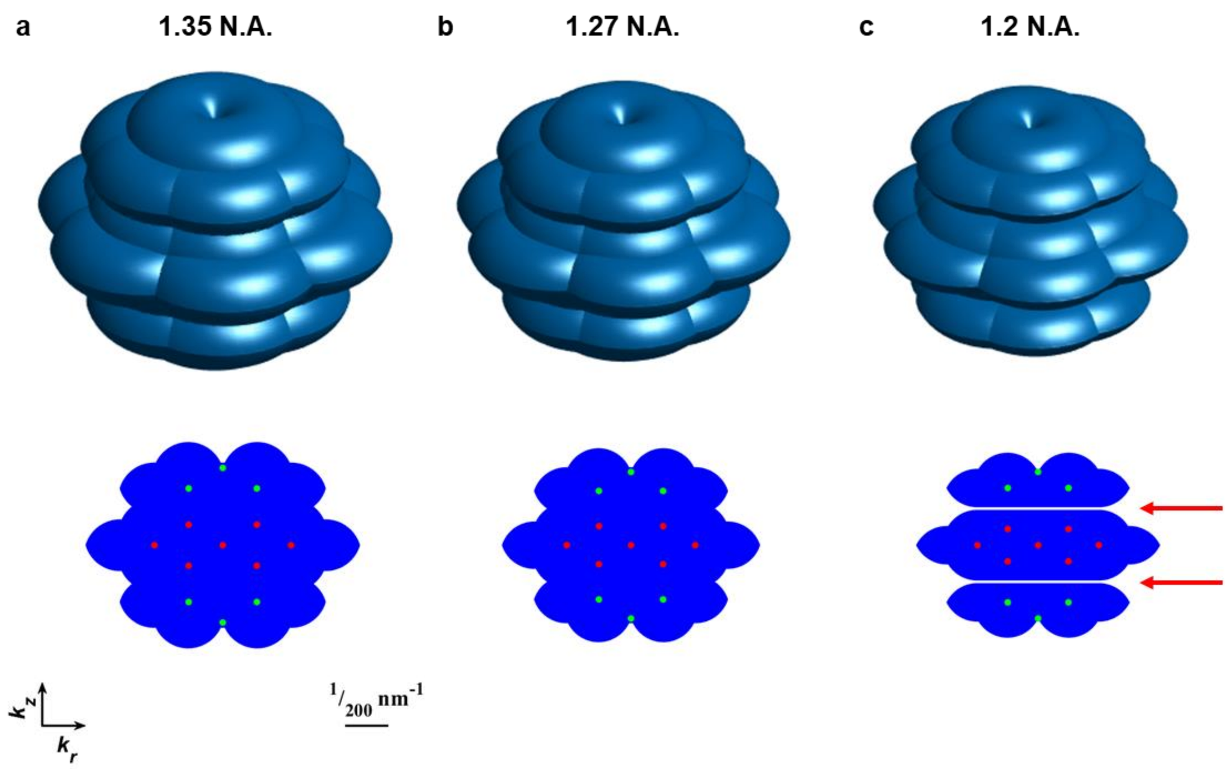


**Supplementary Fig. 2, Support of 4-beam SIM optical transfer functions for commercially available objective lenses. a**) 1.35 NA silicone oil, **b**) 1.27 NA water immersion lens, **c**) 1.2 NA water immersion lens. Top row: 3D projection views; bottom row: axial cross sections as in **Supplementary Fig. 1**. Red arrows in **c**) highlight 'gaps' in spatial frequency support for the 1.2 NA lens case, absent when using the 1.35 NA or 1.27 NA lenses. Simulation parameters: emission NA = 1.35, 1.27, 1.2; refractive index of immersion medium = 1.406 in **a**), 1.33 in **b, c**); excitation wavelength: 488 nm; emission wavelength: 525 nm; side beams at 92% of pupil radius.

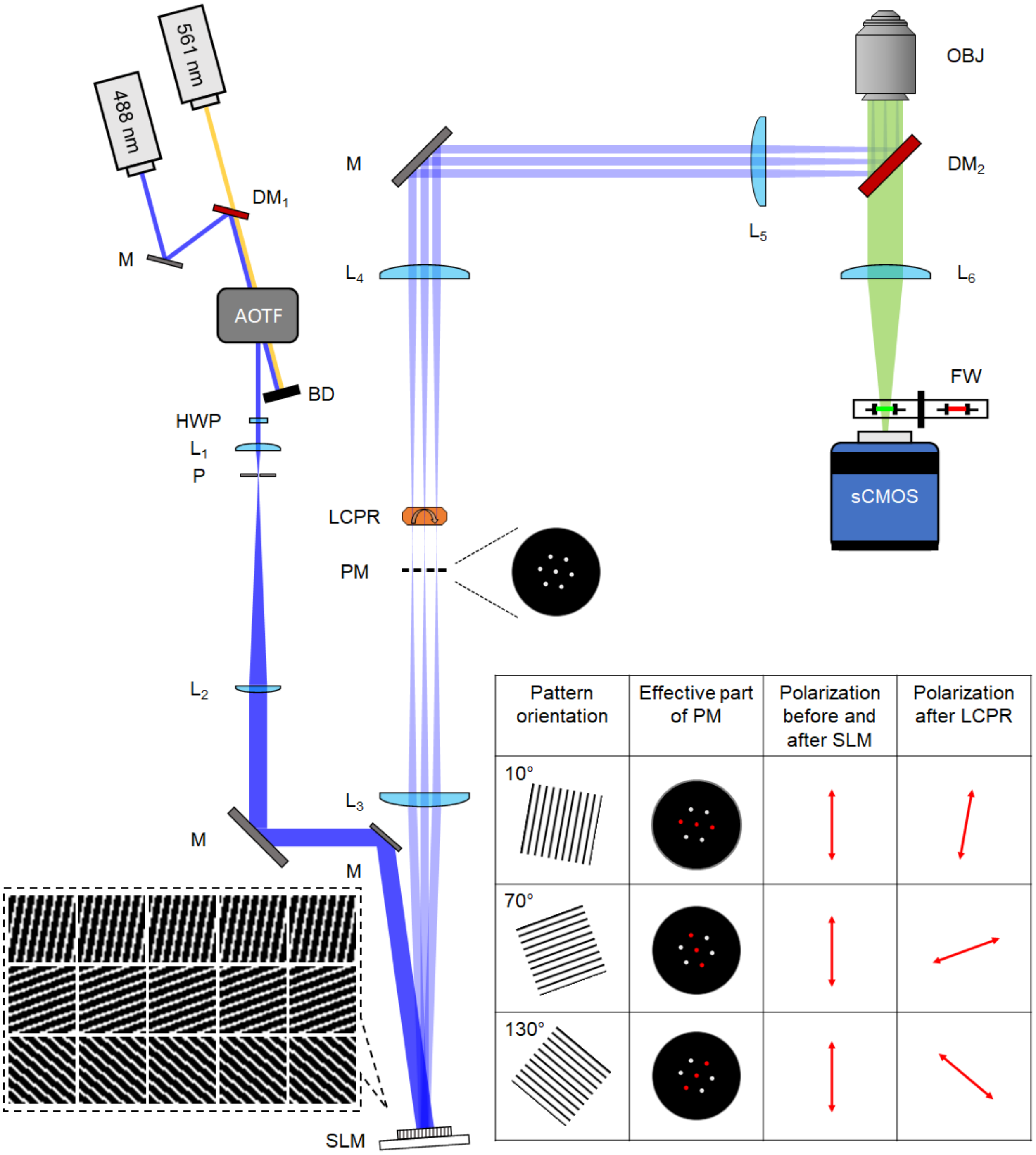


**Supplementary Fig. 3, Optical layout of home-built 3D SIM system used in this work**. The output from linearly polarized, 488 nm and 561 nm diode lasers are combined via a dichroic mirror (DM1) and passed through an acousto-optic tunable filter (AOTF) for rapid shuttering and intensity control. The first-order beam is selected (the zero-order beam is shunted to a beam dump, BD), and spatially filtered (focused via lens L1 onto pinhole P then collimated and expanded via lens L2) before being redirected onto a phase-only spatial light modulator (SLM) at near normal incidence. A half wave plate (HWP) positioned prior to the spatial filter is used to adjust the direction of linear polarization, aligning it for maximum phase modulation by the SLM and thereby ensuring high contrast for the 15 patterns used in 3D SIM (shown in left inset). Lens L3 is positioned one focal length after the SLM, producing a Fourier image of the illumination pattern at its focus. A pinhole mask (PM) placed at this plane serves to filter out unwanted illumination orders. A telescope (lens pair L4 and L5 placed in 4*f* configuration) relays the image of the illuminated mask to the back focal plane of the objective lens (OBJ), which produces the illumination pattern (image of the SLM) at the sample. A liquid crystal polarization rotator (LCPR) is used for rapid rotation of the polarization state post-SLM, producing mostly *s* polarized illumination at the sample and thus high illumination pattern contrast there (right inset shows relationship between pattern orientation, illumination state of the mask, and polarization rotation after LCPR; note images show plane perpendicular to beam propagation). Fluorescence is isolated post-objective via a dichroic mirror (DM2) and imaged to a scientific complementary metal-oxide-semiconductor detector (sCMOS) mounted on a translation stage via tube lens L6. Emission filters mounted in a filter wheel (FW) serve to further isolate fluorescence and select appropriate spectral bands. See **Methods** for further information.

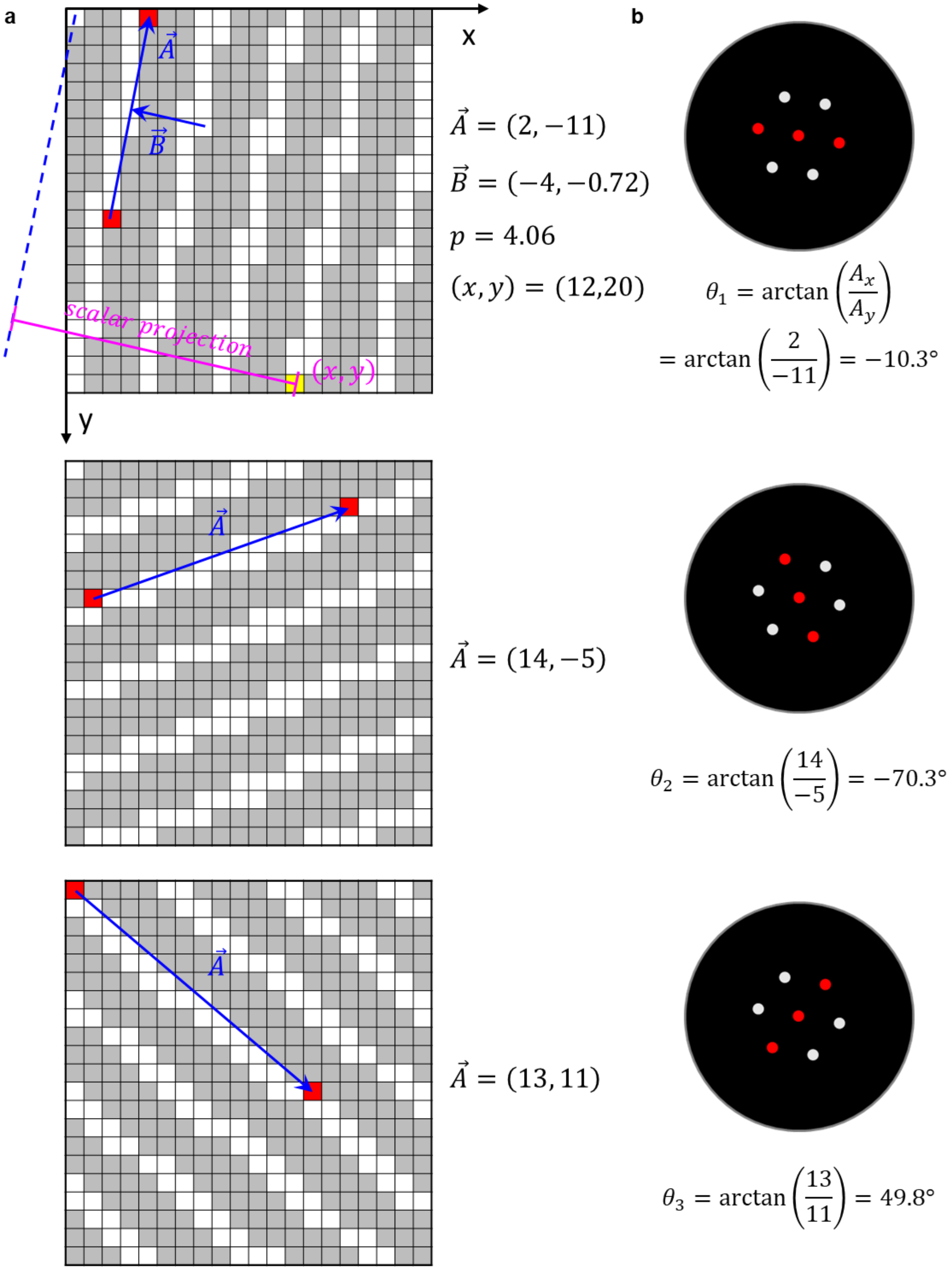
a
b
x
y
$\vec{A}$
$\vec{B}$
scalar projection
$(x, y)$
$\vec{A} = (2, -11)$
$\vec{B} = (-4, -0.72)$
$p = 4.06$
$(x, y) = (12,20)$
$\theta_1 = \arctan\left(\frac{A_x}{A_y}\right)$
$= \arctan\left(\frac{2}{-11}\right) = -10.3°$
$\vec{A} = (14, -5)$
$\theta_2 = \arctan\left(\frac{14}{-5}\right) = -70.3°$
$\vec{A} = (13, 11)$
$\theta_3 = \arctan\left(\frac{13}{11}\right) = 49.8°$

**Supplementary Fig. 4, SLM pattern generation with tunable period and duty cycle. a**) Example SLM patterns with 4.06-pixel period and 33% duty cycle used for 488 nm excitation with 1.27 NA objective. Top panel shows first orientation and defines relevant quantities; other panels show second and third orientations. **b**) The angles and corresponding components (red) transmitted through the PM for the 3 pattern directions. In **a**), the pattern orientation is specified by the blue-colored vector $\vec{A}$ (defined between the two pixels colored in red) and the periodicity $p$ by another blue-colored vector $\vec{B}$. The blue dashed line is the extension along the direction of vector $\vec{A}$ through the origin, drawn for illustration purposes only. For an arbitrary pixel coordinate $(x, y)$ on the SLM (pixel colored in yellow), the magenta projection represents the scalar projection of vector $(x, y)$ onto vector $\vec{B}$, which is -15.4 when $(x, y) = (12,20)$. The modulo after dividing the scalar projection by the magnitude of $\vec{B}$ serves as the criterion for whether the pixel is set to 0 (gray) or π (white) retardance. For example, the magnitude of $\vec{B}$ is 4.06, the modulo after dividing -15.4 by 4.06 is 0.86 pixels, which is smaller than 1.34 pixels (33% duty cycle multiplied with 4.06-pixel period), so (12, 20) was assigned to the on-state (π phase retardance) as shown in the figure.

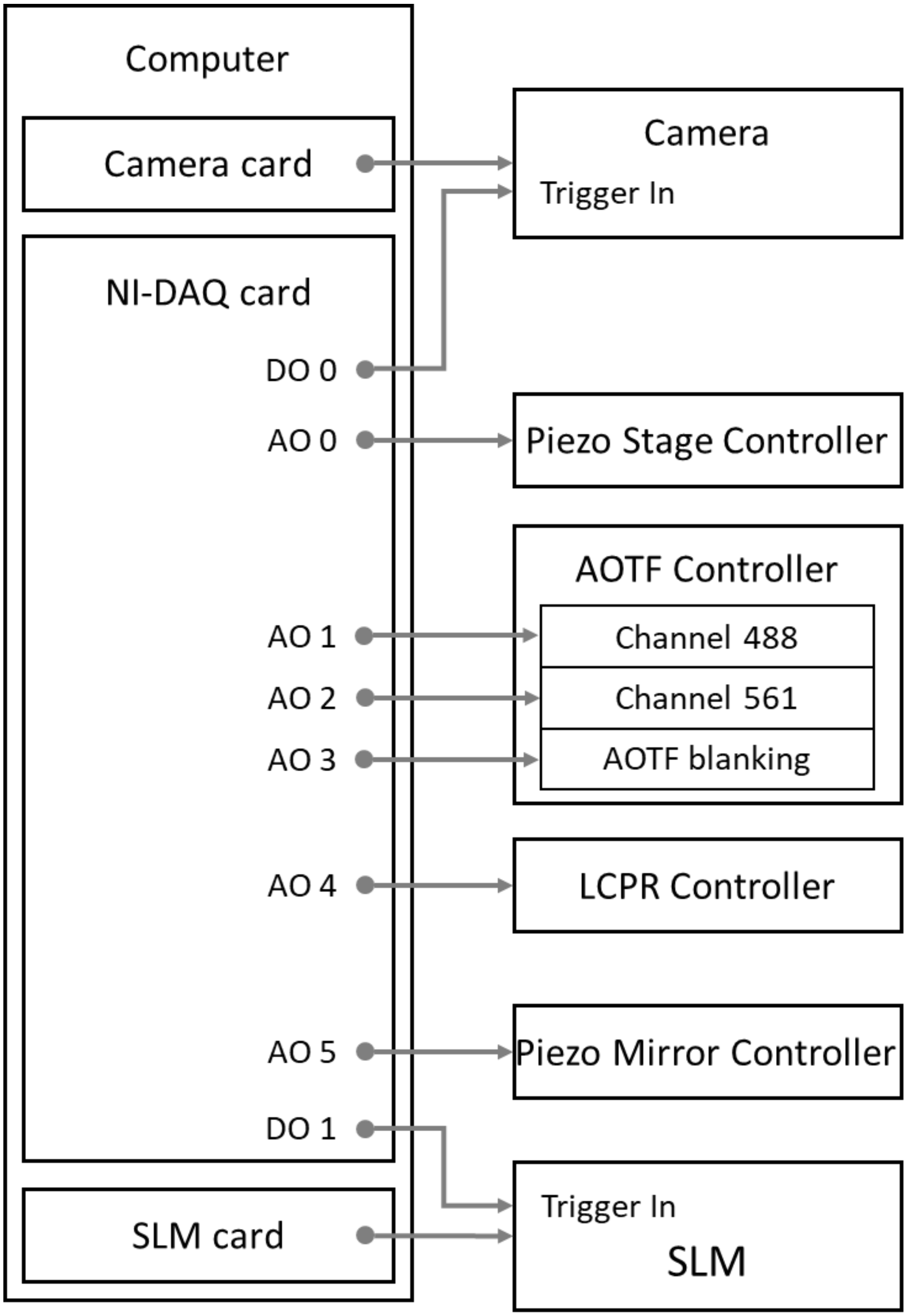


**Supplementary Fig. 5, Hardware control schematic.** DO: digital output. AO: analog output. DAQ: data acquisition. AOTF: acousto-optic tunable filter. SLM: spatial light modulation. LCPR: liquid crystal polarization rotator. Camera acquisition card, NI-DAQ card and SLM control card all reside in the workstation shown by the left box; commands are then issued to devices as indicated. See also **Methods, Supplementary Fig. 6, 7**.

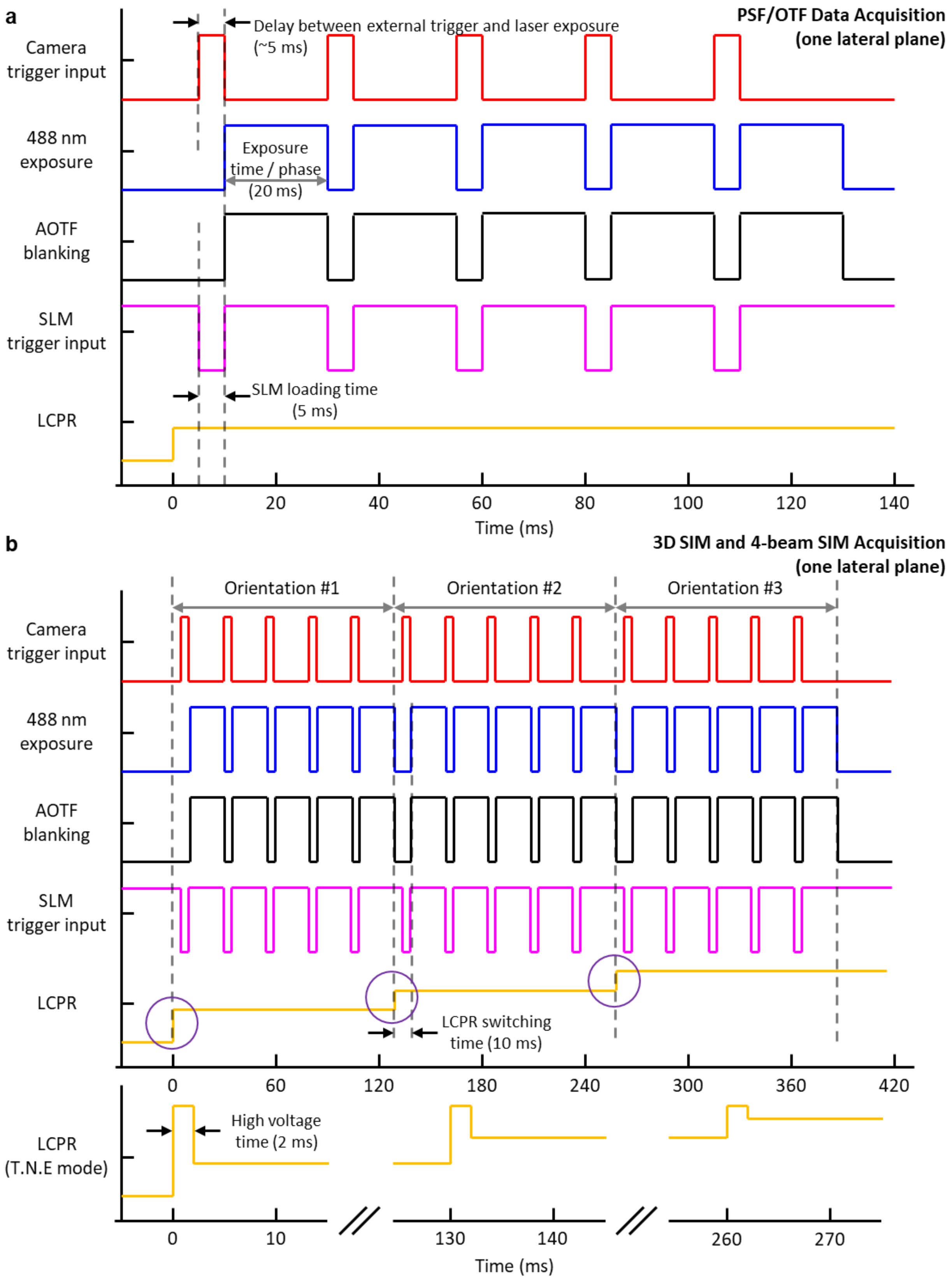
a
PSF/OTF Data Acquisition
(one lateral plane)
Delay between external trigger and laser exposure
(~5 ms)
Camera
trigger input
488 nm
exposure
Exposure
time / phase
(20 ms)
AOTF
blanking
SLM
trigger input
SLM loading time
(5 ms)
LCPR
0
20
40
60
80
100
120
140
Time (ms)
b
3D SIM and 4-beam SIM Acquisition
(one lateral plane)
Orientation #1
Orientation #2
Orientation #3
Camera
trigger input
488 nm
exposure
AOTF
blanking
SLM
trigger input
LCPR
LCPR switching
time (10 ms)
0
60
120
180
240
300
360
420
LCPR
(T.N.E mode)
High voltage
time (2 ms)
0
10
130
140
260
270
Time (ms)

**Supplementary Fig. 6, Timing diagrams for hardware control, single plane acquisition.** Timing diagrams showing control waveforms for **a**) PSF/OTF data acquisition and **b**) 3D SIM and 4-beam SIM acquisition at one lateral plane. The 'Transient Nematic Effect' (T.N.E.) was used to improve the response time of the LCPR, as shown in **b**). See also **Methods** for detailed explanation.

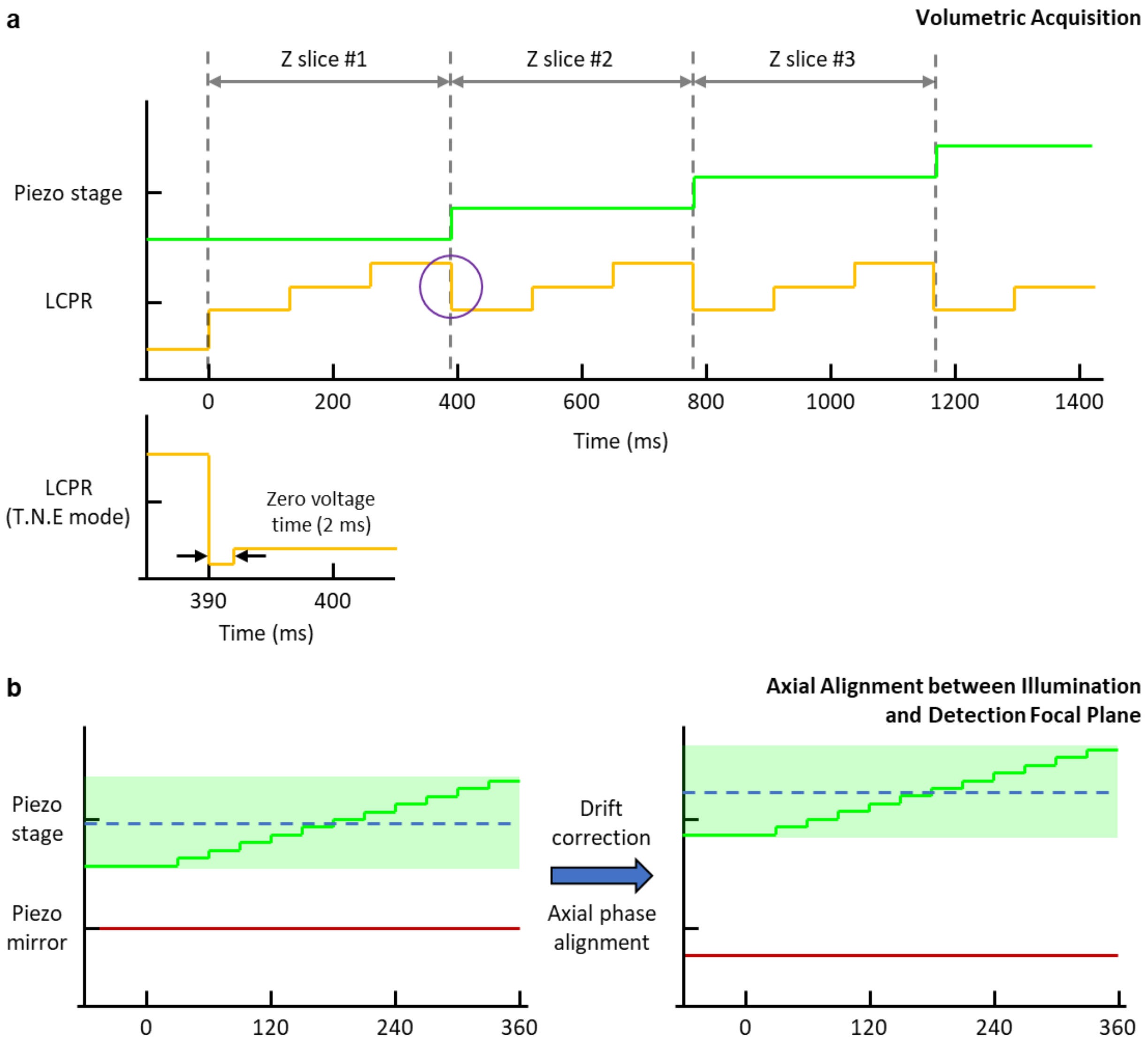


**Supplementary Fig. 7, Timing diagrams for hardware control, volume acquisition.** Timing diagrams showing control waveforms for **a**) 'naïve' volumetric acquisition (used for 3D SIM acquisition) and **b**) acquisition after axial phase alignment between illumination and detection focal plane (e.g., for 4-beam SIM).

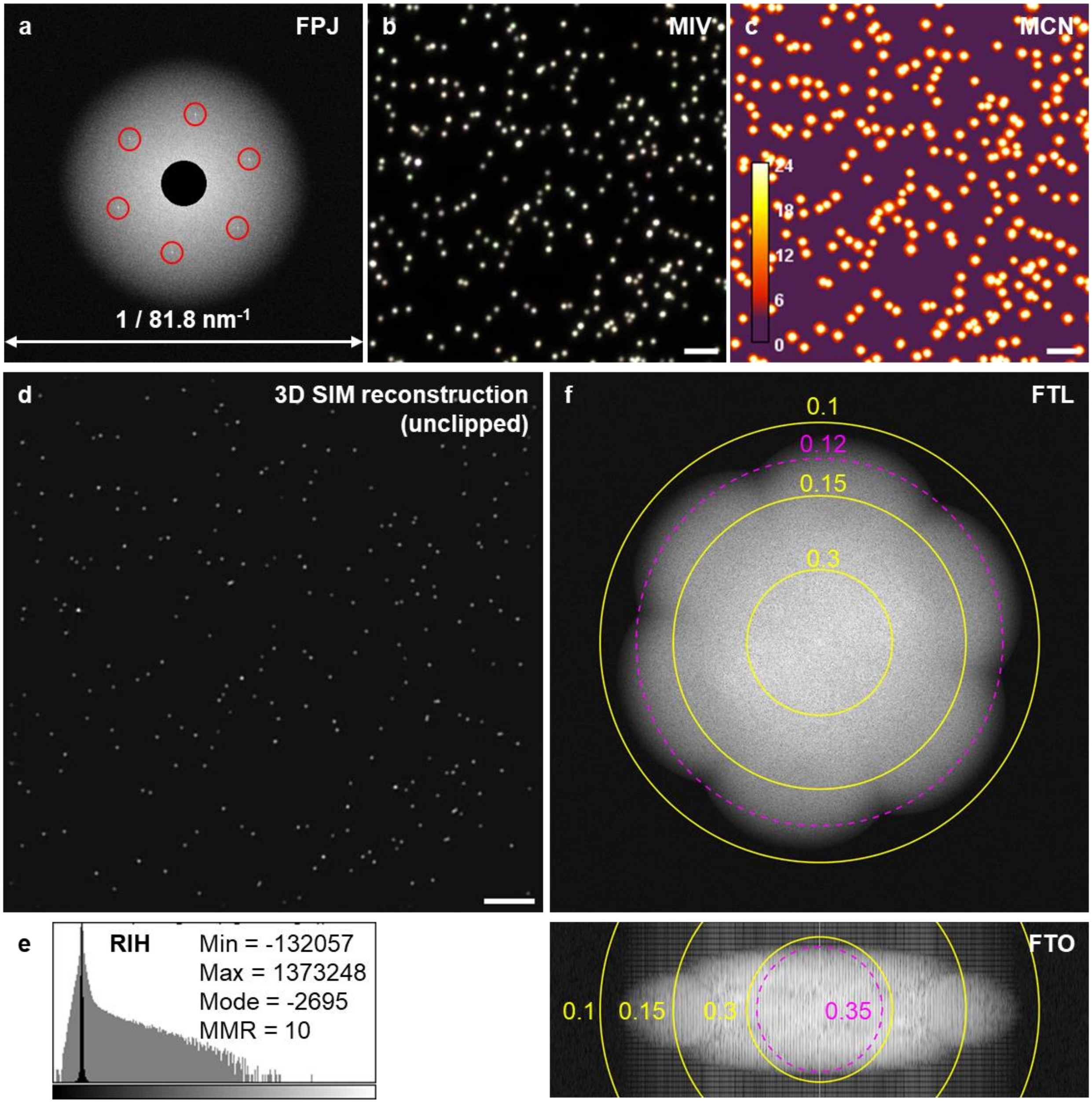


**Supplementary Fig. 8, SIMcheck output for 100 nm yellow-green beads imaged with 1.27 NA water objective. a**) 'Raw Fourier Projection (FPJ)' of raw 3D SIM input data. The FPJ highlights spots corresponding to the first-order frequencies in the illumination pattern for each orientation. **b**) 'Motion & Illumination Variation (MIV)' of raw SIM data. Each orientation is phase-averaged, normalized, and pseudo-colored with cyan, magenta and yellow. The gray- to white-only MIV here demonstrates that there is not significant movement of the beads during SIM acquisition. **c**) 'Modulation Contrast-to-Noise (MCN)' of raw SIM data. This function estimates the local modulation contrast. All beads have MCN values larger than 18, which demonstrates excellent modulation contrast. **d**) The reconstructed 3D SIM image (single lateral plane shown) with negative values preserved (unclipped). **e**) 'Reconstructed Intensity Histogram (RIH)' of data corresponding to **d**). Linear-scaled (black) and log-scaled (gray) intensity histograms show the contribution of floating-point values of the reconstructed data. The intensity minima, maxima, mode and the minimum-to-maximum ratio (MMR) are also shown in the figure. MMR computes the feature intensity relative to the reconstructed noise and intensity dips generated in the reconstruction process. An MMR value of 10 indicates good signal-to-noise ratio. **f**) 'Reconstructed Fourier Plots' of data corresponding to **d**). 'Fourier Transform Lateral (FTL)' and 'Fourier Transform Orthogonal (FTO)' are log-scaled projections of the 3D FFT of the reconstructed data. Both projections are shown superimposed with the corresponding dimensions in real space (yellow rings and values in microns). The theoretical frequency/resolution limit are also indicated (magenta dashed circles). Scale bars: 2 µm.

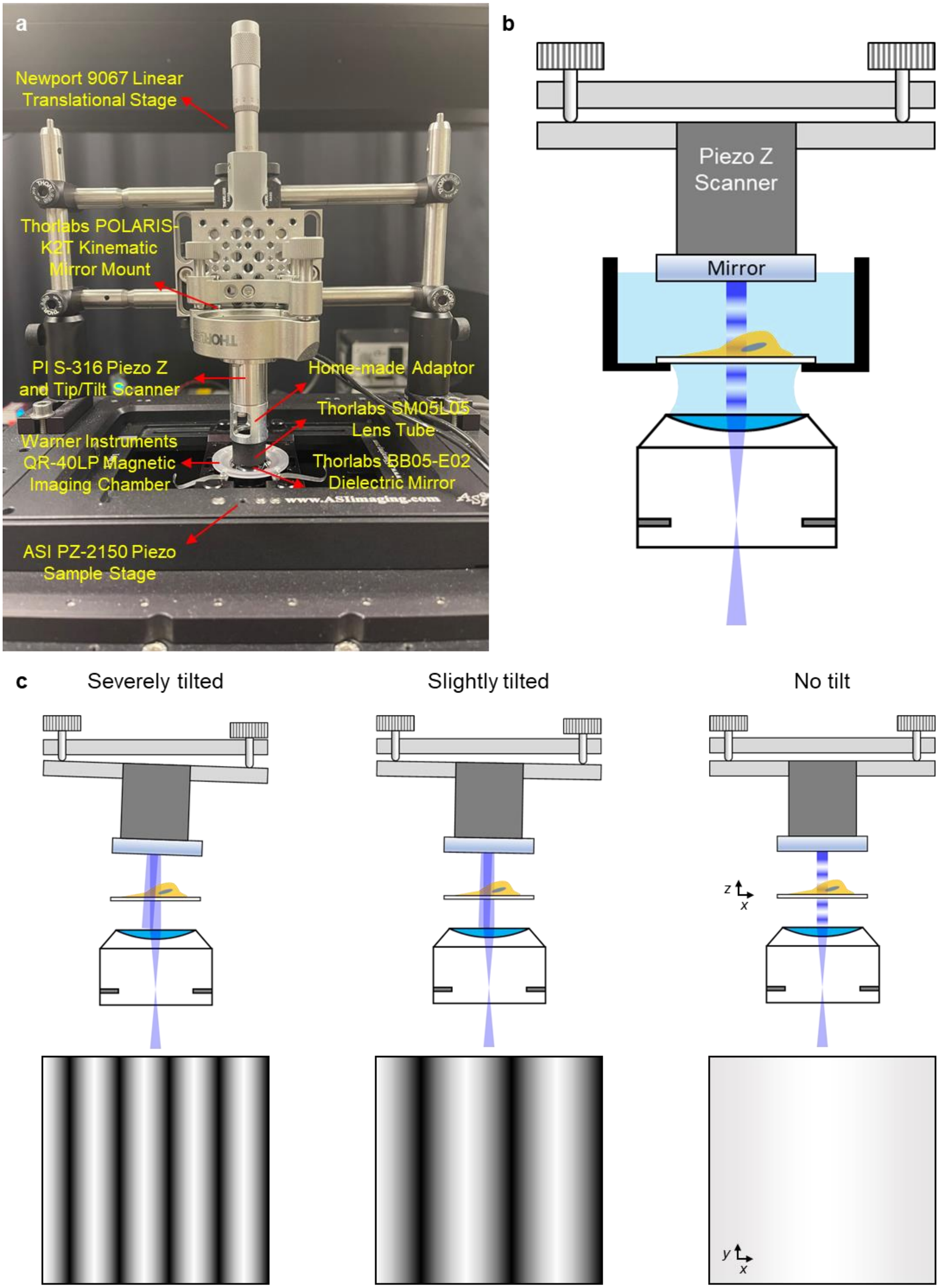
a
Newport 9067 Linear Translational Stage
Thorlabs POLARIS-K2T Kinematic Mirror Mount
PI S-316 Piezo Z and Tip/Tilt Scanner
Home-made Adaptor
Thorlabs SM05L05 Lens Tube
Warner Instruments QR-40LP Magnetic Imaging Chamber
Thorlabs BB05-E02 Dielectric Mirror
ASI PZ-2150 Piezo Sample Stage
b
Piezo Z Scanner
Mirror
c
Severely tilted
Slightly tilted
No tilt
z
x
y
x

**Supplementary Fig. 9, Mounting and aligning reflective mirror for 4-beam SIM. a**) Photograph of mounting scheme, indicating hardware mounts with vendor information. PI: Physik Instrumente. **b**) Schematic to accompany **a**), showing kinematic mirror mount, Piezo Z scanner, mirror, sample, and objective lens. Schematic is not to scale. **c**) Alignment of the reflected beam is achieved by manually adjusting the knobs on the kinematic mirror mount until tilt is minimized. This can be achieved by monitoring the autofluorescence from a dirty coverslip immersed in liquid; tilt is minimized when a flat intensity profile is achieved when translating the sample through focus, yielding a purely axial modulation. Top images show reflected beam at different tilts of mirror; bottom images show corresponding planar images of autofluorescence when lateral modulation is present (left, middle) vs. mostly axial modulation (right). See also **Supplementary Video 1** and **Methods** for more information.

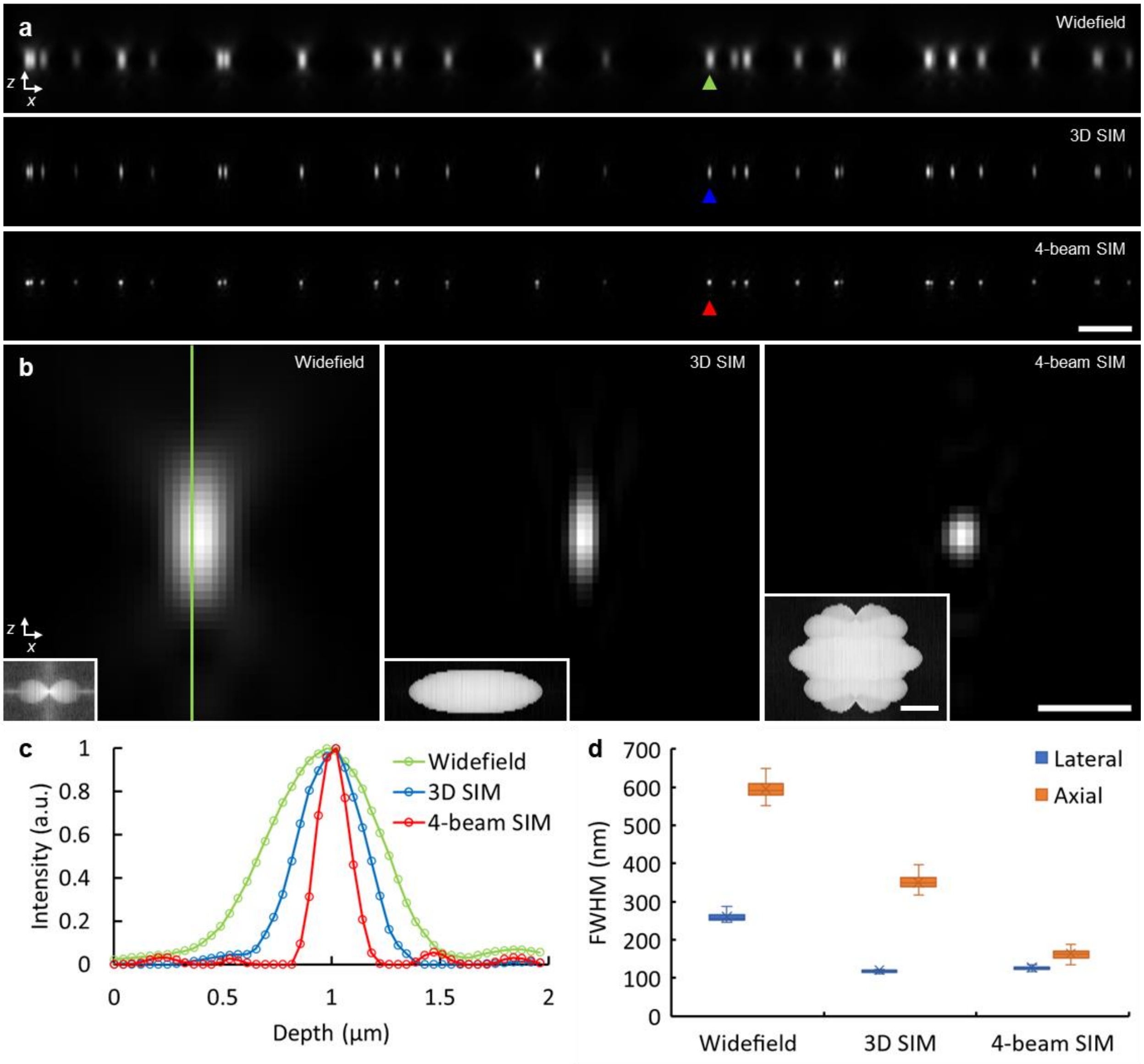


**Supplementary Fig. 10, Estimating spatial resolution for 1.27 NA water lens**. **a**) xz cross sectional views of 100 nm yellow-green beads, as viewed with widefield microscopy (top), 3D SIM (middle), and 4-beam SIM (bottom). Scale bar: 2 μm. **b**) Higher magnification views of bead marked by green, blue, and red arrowheads in **a**). Insets show magnitude of corresponding Fourier transform. Scale bar: 500 nm, 200 $nm^{-1}$ (inset). **c**) Line profiles taken along vertical line shown in **b**). **d**) Full width at half maximum (FWHM) analysis from *N* = 85, 81, 78 beads, showing lateral (blue) and axial (orange) values for the three methods. See also **Fig. 1**, **Supplementary Table 1**.

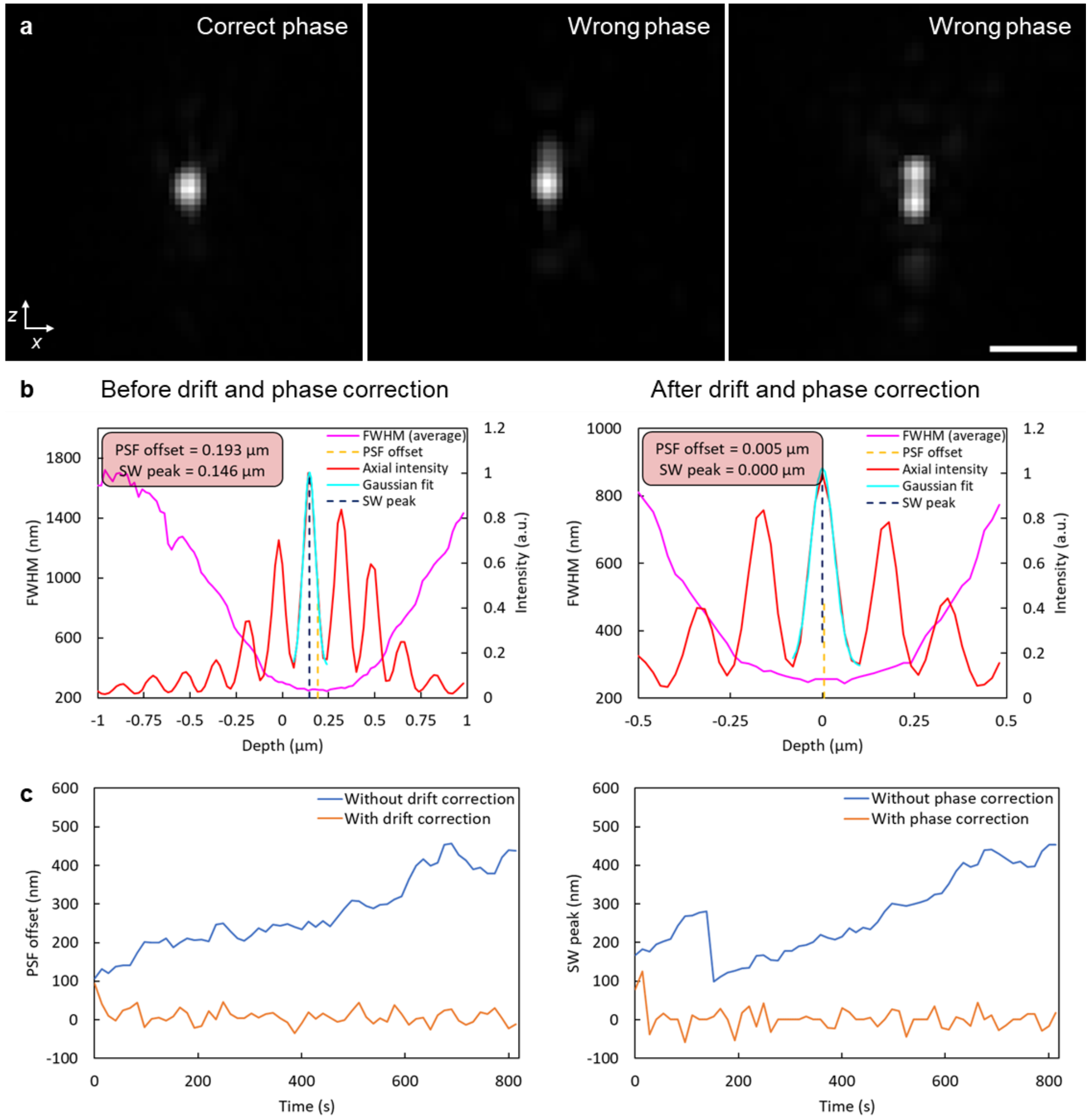


**Supplementary Fig. 11, Drift and relative phase correction for 4-beam SIM. a**) Image of reconstructed bead after 4-beam SIM acquisition with correct (left) and incorrect (middle, right) relative phase between detection plane and illumination pattern maxima. When the phase difference is minimized (left) the bead appears nearly isotropic; progressively larger phase differences culminate in the appearance of a doubled bead (right). Scale bar: 500 nm. **b**) Axial position of standing wave illumination (red, 'Axial intensity') and center of bead (magenta, 'FWHM (average)') before (left) and after (right) drift and relative phase correction. Before correction, the peak standing wave illumination maxima (dark blue dashed line, 'SW peak') and bead position (orange dashed line, 'PSF offset') are offset from the focal plane position and from each other. After correction, bead and illumination maxima are within 5 nm of each other and the focal plane position. **c**) Examples of temporally varying axial bead position drift (left) and illumination pattern maxima drift (right) with (orange) and without (blue) active correction. Excluding the first three timepoints (when the system is settling), positional standard deviation after active correction is s = 18 nm for bead, 20 nm for illumination pattern maxima vs. 91 nm and 103 nm without active correction.

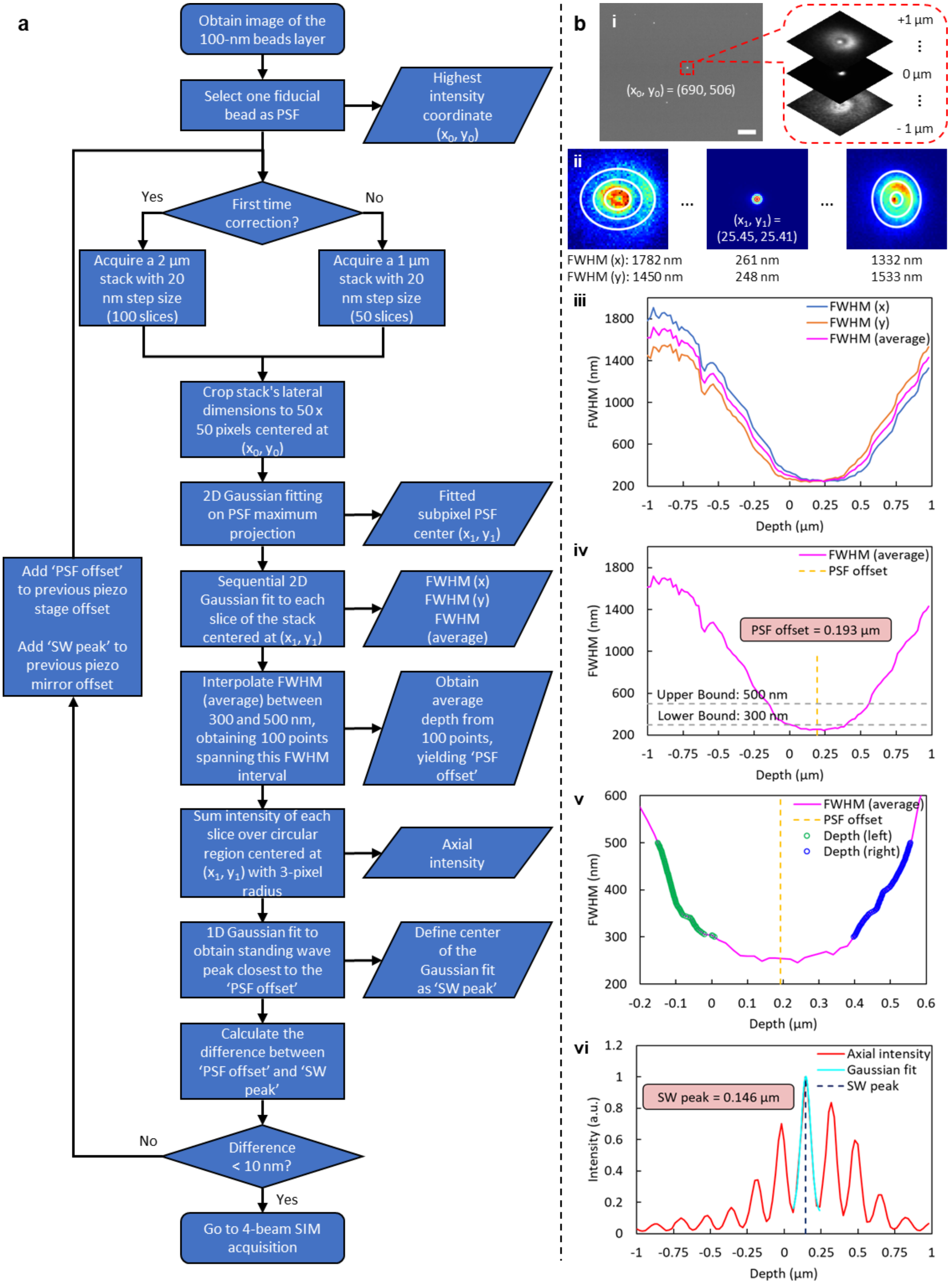

a
Obtain image of the 100-nm beads layer
Select one fiducial bead as PSF
Highest intensity coordinate ($x_0$, $y_0$)
First time correction?
Yes
No
Acquire a 2 µm stack with 20 nm step size (100 slices)
Acquire a 1 µm stack with 20 nm step size (50 slices)
Crop stack's lateral dimensions to 50 x 50 pixels centered at ($x_0$, $y_0$)
2D Gaussian fitting on PSF maximum projection
Fitted subpixel PSF center ($x_1$, $y_1$)
Sequential 2D Gaussian fit to each slice of the stack centered at ($x_1$, $y_1$)
FWHM (x) FWHM (y) FWHM (average)
Interpolate FWHM (average) between 300 and 500 nm, obtaining 100 points spanning this FWHM interval
Obtain average depth from 100 points, yielding 'PSF offset'
Sum intensity of each slice over circular region centered at ($x_1$, $y_1$) with 3-pixel radius
Axial intensity
1D Gaussian fit to obtain standing wave peak closest to the 'PSF offset'
Define center of the Gaussian fit as 'SW peak'
Calculate the difference between 'PSF offset' and 'SW peak'
Difference < 10 nm?
No
Yes
Add 'PSF offset' to previous piezo stage offset
Add 'SW peak' to previous piezo mirror offset
Go to 4-beam SIM acquisition
b
i
($x_0$, $y_0$) = (690, 506)
+1 µm
0 µm
- 1 µm
ii
($x_1$, $y_1$) = (25.45, 25.41)
FWHM (x): 1782 nm
FWHM (y): 1450 nm
261 nm
248 nm
1332 nm
1533 nm
iii
FWHM (x)
FWHM (y)
FWHM (average)
FWHM (nm)
Depth (µm)
iv
FWHM (average)
PSF offset
PSF offset = 0.193 µm
Upper Bound: 500 nm
Lower Bound: 300 nm
FWHM (nm)
Depth (µm)
v
FWHM (average)
PSF offset
Depth (left)
Depth (right)
FWHM (nm)
Depth (µm)
vi
Axial intensity
Gaussian fit
SW peak
SW peak = 0.146 µm
Intensity (a.u.)
Depth (µm)

**Supplementary Fig. 12, Bead-based alignment protocol. a**) Flowchart showing procedure for updating position of piezo mirror and piezo stage. FWHM: full width at half maximum. See **Methods** for detailed explanation of all steps. **b**) Schematic images and graphs to accompany **a**). *i)* xy image (left, Scale bar: 10 µm) and cropped stack (right) of indicated bead. *ii)* Images of selected planes in stack, showing astigmatism and example FWHM(x) and FWHM(y) values. *iii)* FWHM(x), FWHM(y), and average of the two curves. *iv, v)* PSF offset is computed by interpolating FWHM(average) and considering 100 interpolated values spanning e.g., from FWHM = 500 nm to FWHM = 300 nm. The midpoints between left and right values for all 100 values are averaged, yielding PSF offset. *vi)* A Gaussian fit on the maxima of the standing wave axial intensity profile is used to find the peak of the standing wave intensity nearest to the PSF offset.

**a**

$$Fluorescence\ Loss = \frac{Average\ Intensity\ (2nd\ Image)\ -\ Background}{Average\ Intensity\ (1st\ Image)\ -\ Background}$$

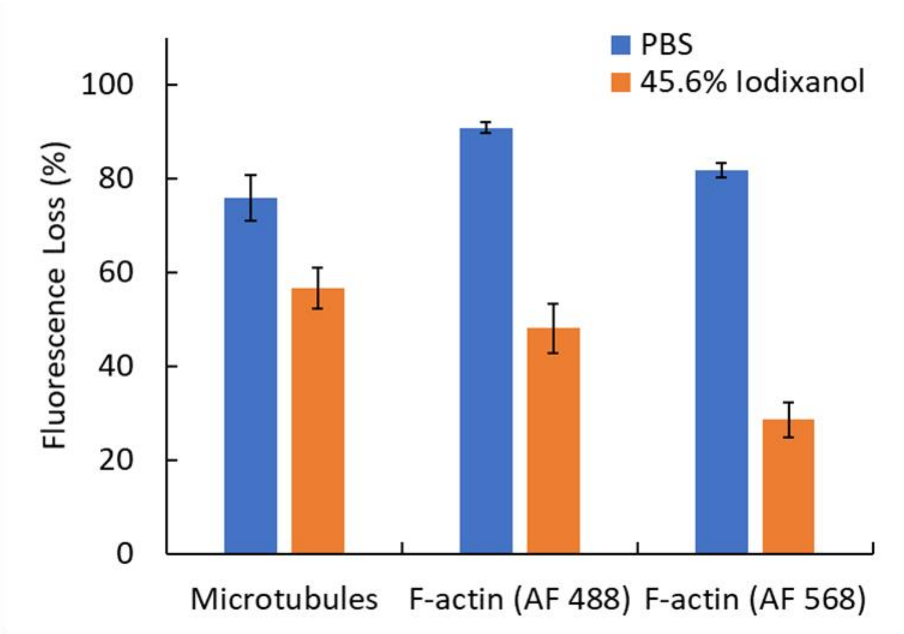


**b**

1st 3D SIM measurement | 2nd 3D SIM measurement

PBS

Iodixanol

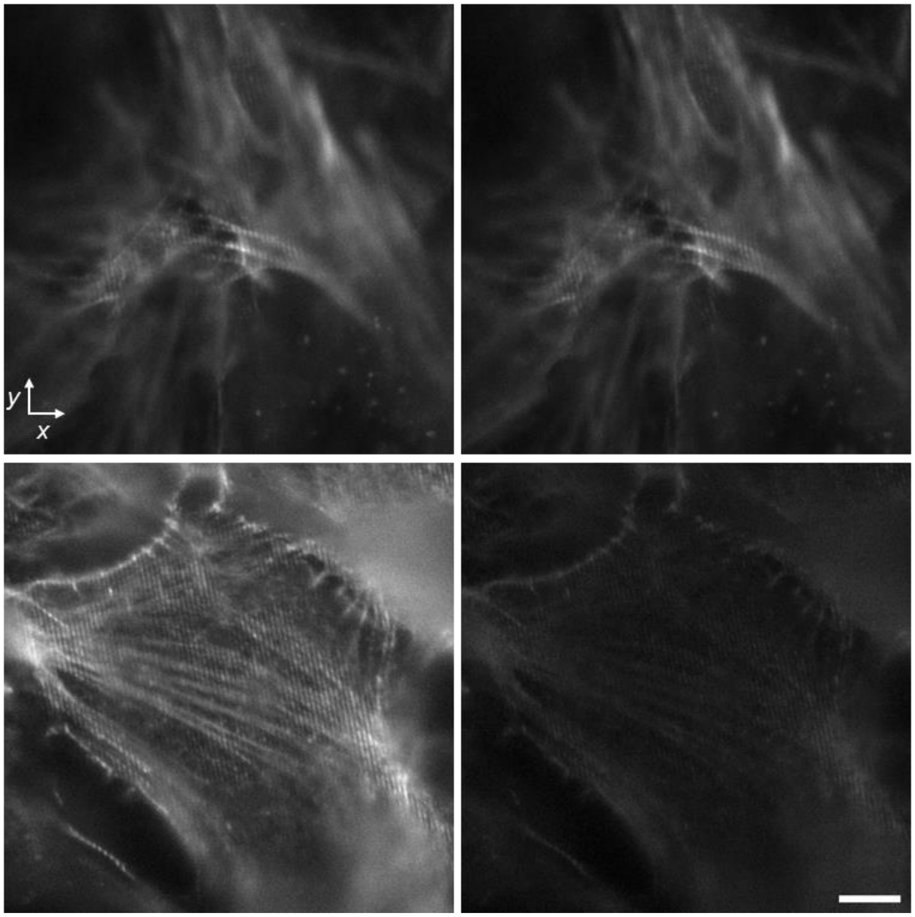

**Supplementary Fig. 13, More bleaching in iodixanol than PBS. a**) Comparing loss in fluorescence after 3D SIM imaging twice in PBS (blue) vs. 45.6% iodixanol in water (orange). Imaging in iodixanol results in more bleaching in immunolabeled microtubules (left, labeled with Alexa Fluor (AF) 488) and phalloidin-stained F-actin (middle, labeled with AF 488; right, labeled with AF 568). Quantification was performed by selecting areas within images of fixed cells and analyzing according to the indicated ratiometric equation. *N* = 18 measurements for microtubules (gathered from 2 cells), 18 measurements for AF 488 labeled actin (from 2 cells) and 18 measurements for AF 568 labeled actin (from 3 cells). **b**) Example raw frames after first 3D SIM measurement (left) and second 3D SIM measurement (right) of Alexa Fluor 488 labeled F-actin, in PBS (top) and iodixanol (bottom). Scale bar: 5 µm.

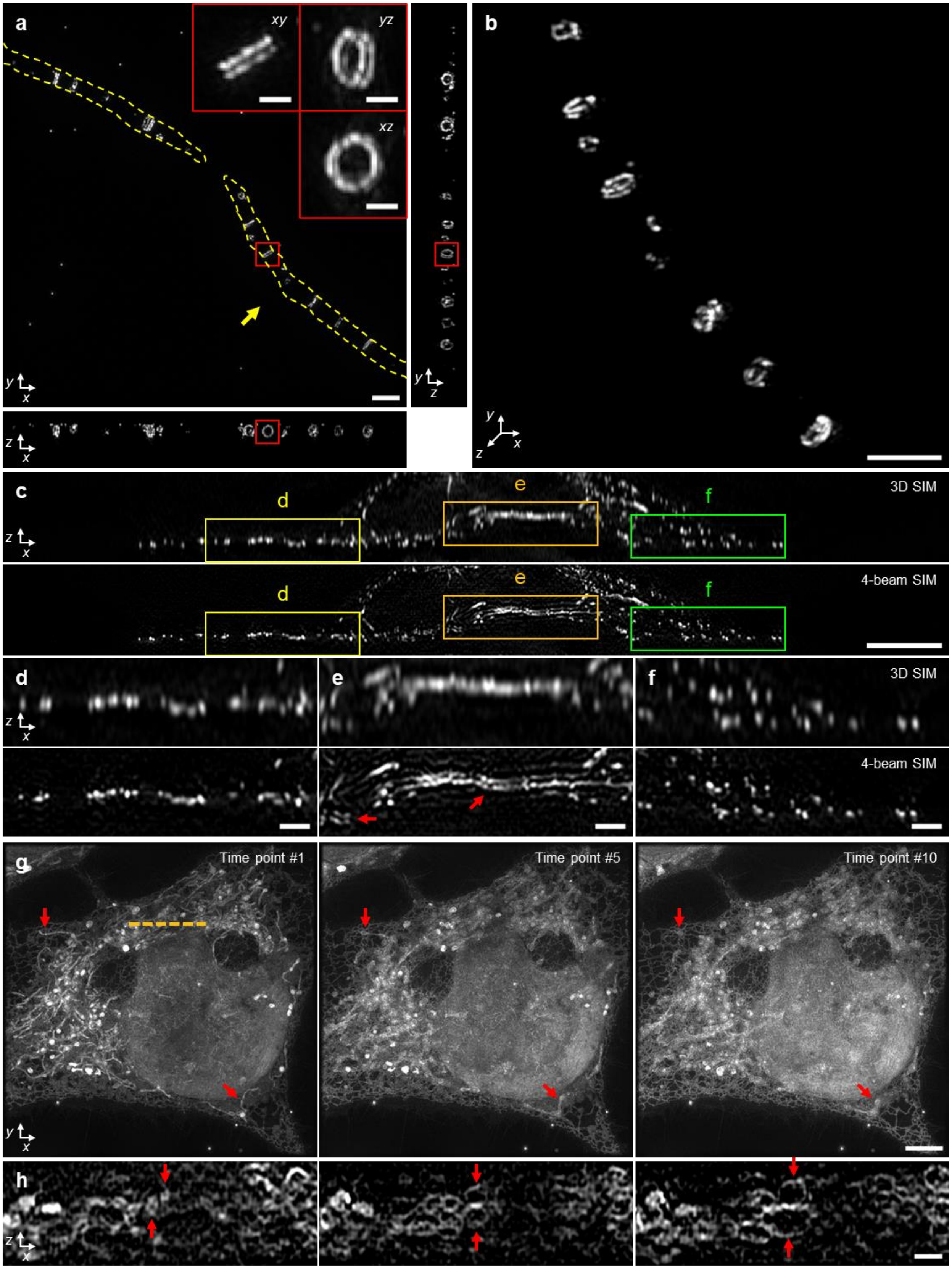
a
xy
yz
xz
y
x
y
z
z
x
b
y
z
x
c
d
e
f
3D SIM
z
x
d
e
f
4-beam SIM
d
e
f
3D SIM
z
x
4-beam SIM
g
Time point #1
Time point #5
Time point #10
y
x
h
z
x

**Supplementary Fig. 14, Additional samples imaged with 4-beam SIM, 1.27 NA water immersion lens.** **a**) Live, vegetative *B. subtilis* with GFP-DivIVA marker. Yellow dashed lines have been added to better delineate cell boundaries. Three maximum intensity projections are shown in lateral and axial views. Insets show higher magnification views of red rectangular regions, indicating clear double rings. **b**) Three-dimensional maximum intensity projection view of the same cell marked by yellow arrow in **a**). See also **Supplementary Video 5**. **c**) Axial views of Alexa Fluor 488 immunolabeled microtubules in fixed U2OS cells, as viewed in 3D SIM (top) and 4-beam SIM (bottom). Higher magnification views of yellow **d**), orange **e**), and green **f**) regions are also shown, with red arrows marking 'doubling'/'ringing' artifacts of microtubules at the nuclear boundary, that likely arise due to refractive index mismatch at nuclear boundary. **g, h**) Live U2OS cell stained with 200 nM Potomac Gold and imaged in 4-beam SIM in lateral (**g**), maximum intensity projection) and axial (**h**)**,** single plane corresponding to orange dashed line in **g**)). Selected time points from a 4D acquisition are shown. Red arrows mark swelling mitochondria, indicating photodamage. See also **Supplementary Video 7**. Scale bars: 2 μm (500 nm for insets) **a, b**); 5 μm **c, g**); 1 μm **d-f, h**).

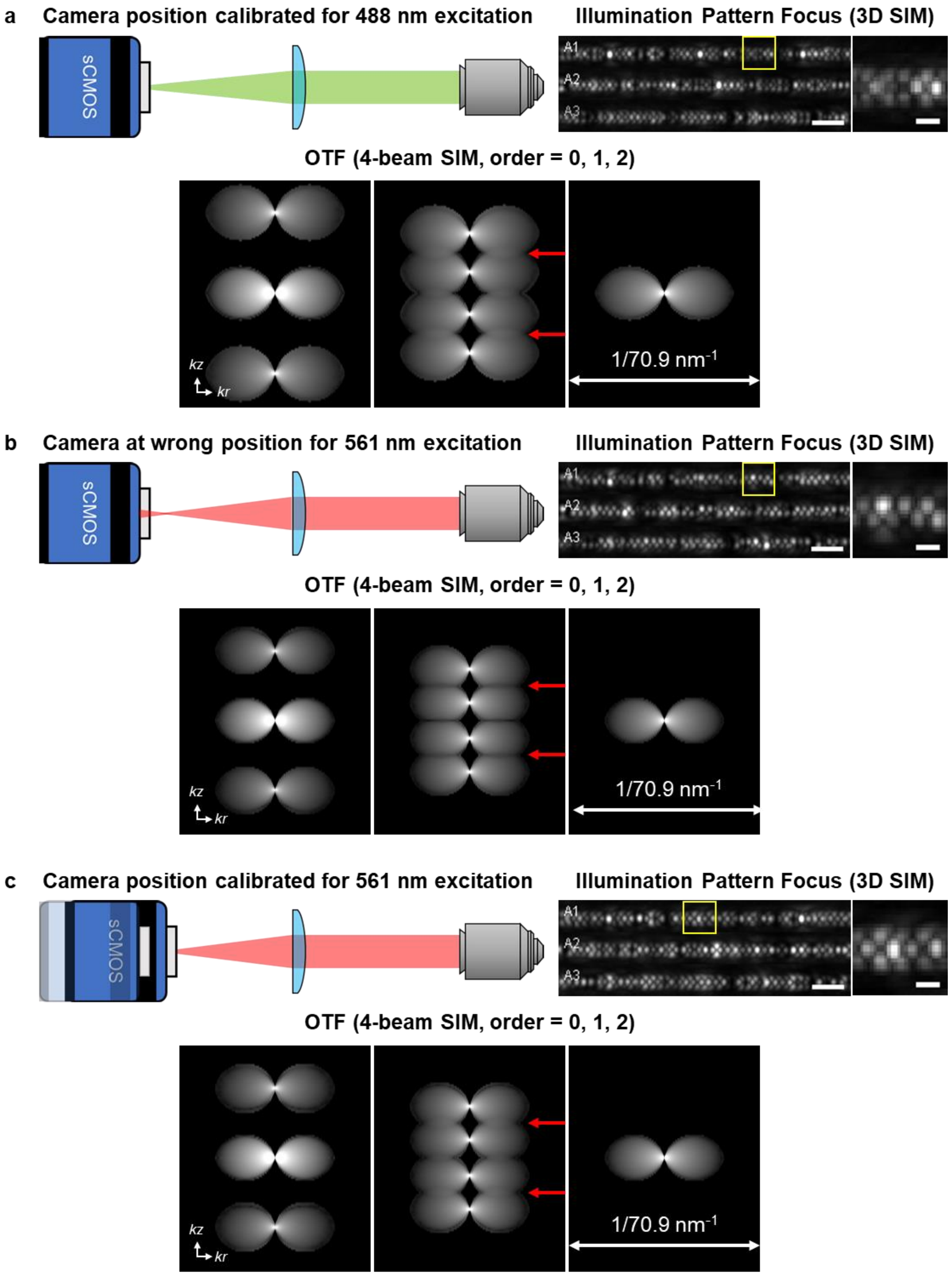
a
Camera position calibrated for 488 nm excitation
sCMOS
Illumination Pattern Focus (3D SIM)
A1
A2
A3
OTF (4-beam SIM, order = 0, 1, 2)
kz
kr
1/70.9 nm-1
b
Camera at wrong position for 561 nm excitation
sCMOS
Illumination Pattern Focus (3D SIM)
A1
A2
A3
OTF (4-beam SIM, order = 0, 1, 2)
kz
kr
1/70.9 nm-1
c
Camera position calibrated for 561 nm excitation
sCMOS
Illumination Pattern Focus (3D SIM)
A1
A2
A3
OTF (4-beam SIM, order = 0, 1, 2)
kz
kr
1/70.9 nm-1

**Supplementary Fig. 15, Adjusting the relative phase between the detector plane and illumination pattern. a**) When the camera position (detector plane) is adjusted correctly for the 488 nm illumination pattern, order 1 optical transfer functions (OTFs) overlap (red arrows), producing high quality reconstructions. In practice, we adjust the camera position while monitoring a thin layer of 100 nm yellow-green beads, acquiring 3D SIM data, and passing it through the SIMCheck software. When the illumination pattern is aligned, axial views of beads at each orientation (A1, A2, A3) appear symmetric, with approximately equal energy appearing above and below the central intensity maxima (right, 'Illumination Pattern Focus 3D SIM'). **b**) When imaging red fluorescent beads excited with 561 nm illumination, if the detector plane is not moved, the beads appear asymmetric, and gaps appear in the order 1 OTF. **c**) Translating the camera restores OTF overlap and beads appear symmetric again. Scale bars: 2 µm lower magnification views, 500 nm higher magnification views of yellow rectangular regions.

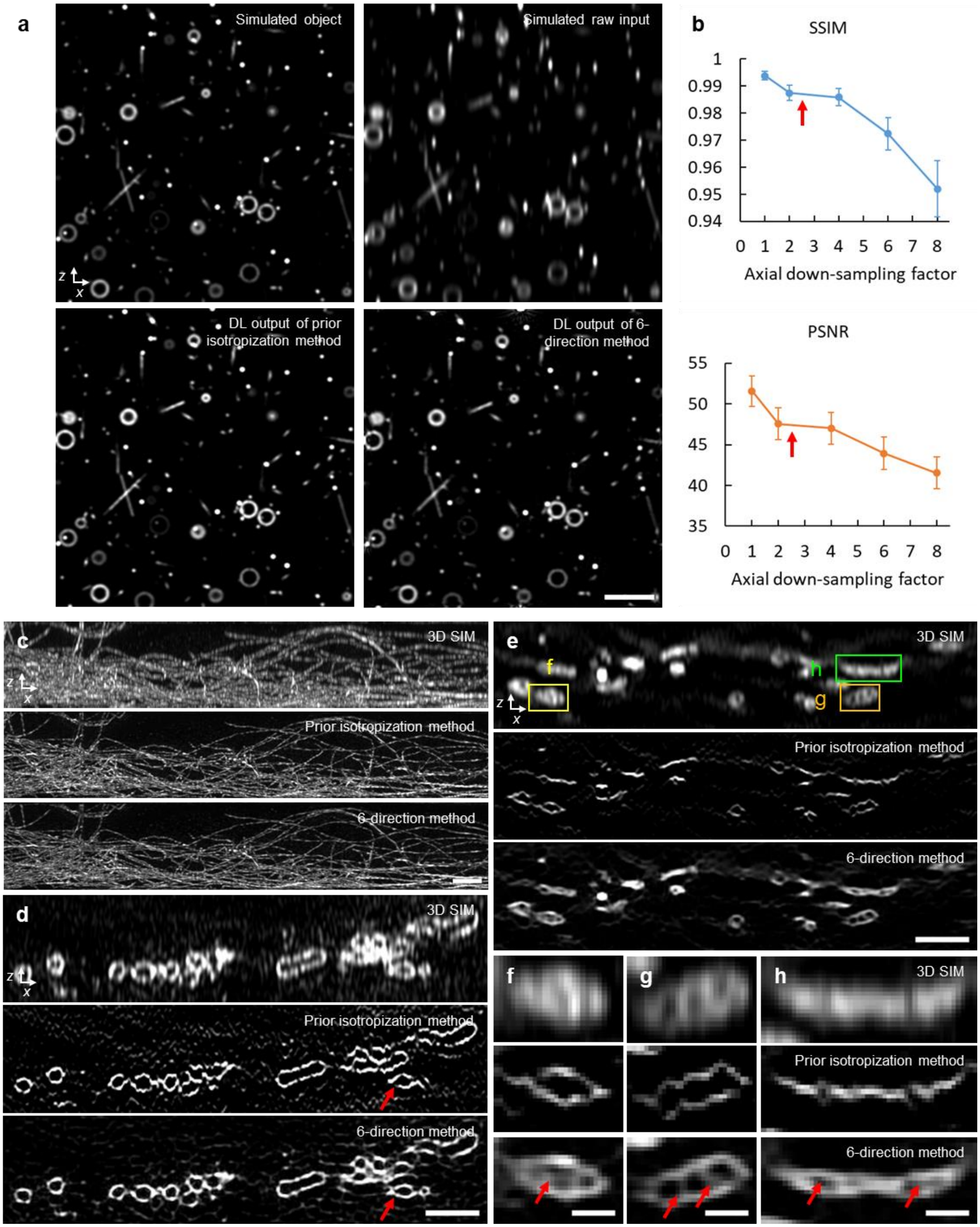


**Supplementary Fig. 16, Comparing methods for network-based isotropization. a**) Simulated objects consisting of lines, points, and hollow spheres (upper left) were blurred and downsampled to simulate 3D SIM data (upper right). Both prior network (lower left, trained only on lateral xy views) and current network (lower right, which considers six rotated axial views of the sample) produce high quality predictions resembling the ground truth. **b**) Structural similarity index (SSIM) and peak signal-to-noise-ratio (PSNR) as a function of axial downsampling factor, in the six-direction method. Red arrows show downsampling factor used in this work (2.5). **c**) Axial views of immunolabeled microtubules in fixed U2OS cells, as viewed in 3D SIM (top), output of prior network (middle), and current network (bottom). Note close visual similarity between network outputs. **d**) As for **c**), but images are of immunolabeled Tomm20 in fixed U2OS cells, marking the outer mitochondrial membrane. Red arrows highlight 'breaks' in the membrane that occur in previous network output. **e**) As for **c, d**), but images show MitoTracker Green FM label in live U2OS cells. **f-h**) Higher magnification views of rectangular regions in **e**), illustrating features resolved in current work but absent (**f, g**) or distorted (**h**) in others (red arrows). Scale bars: 2 µm **a, c-e**); 500 nm **f-h**).

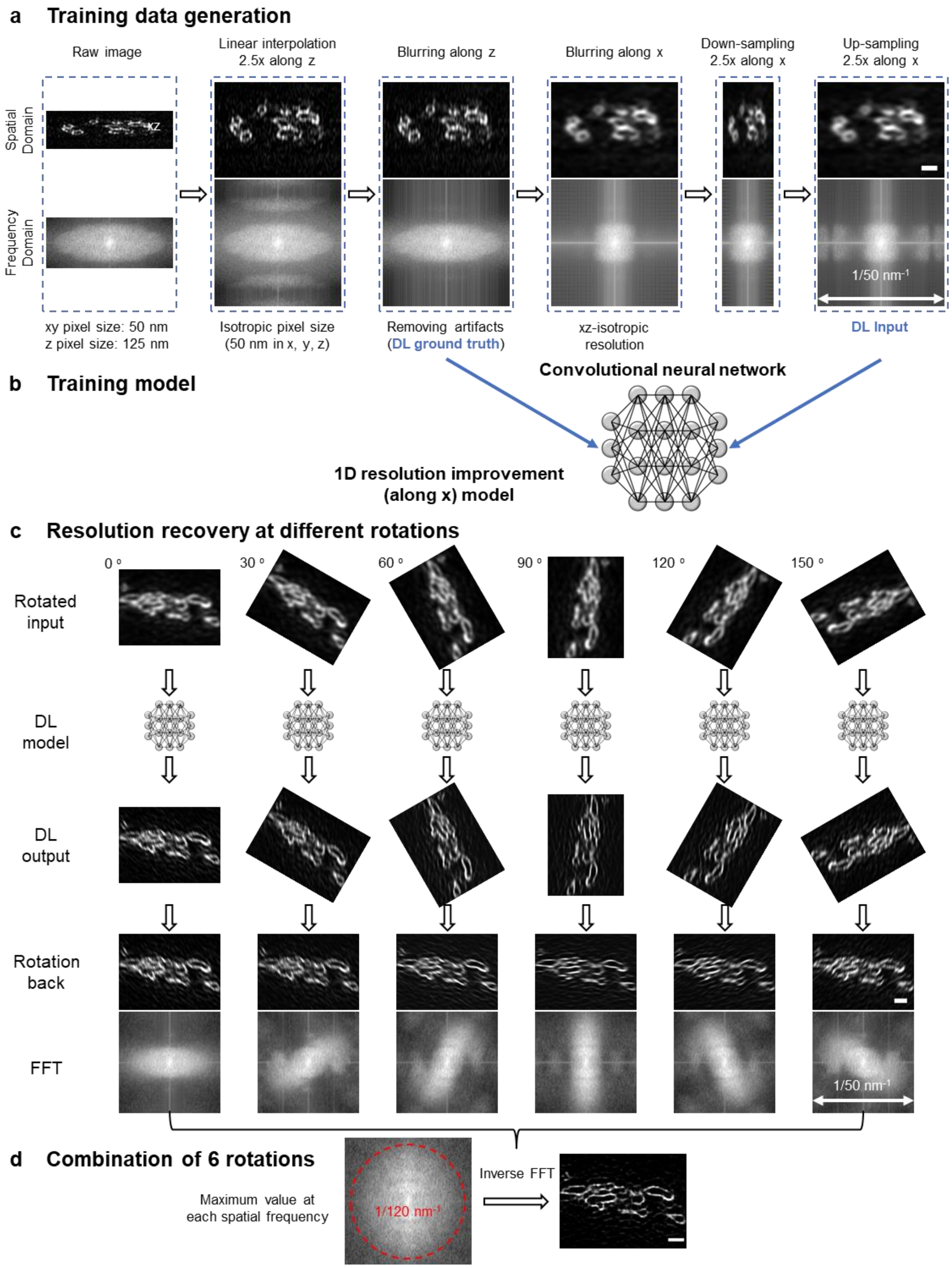

a Training data generation
Raw image
Linear interpolation 2.5x along z
Blurring along z
Blurring along x
Down-sampling 2.5x along x
Up-sampling 2.5x along x
Spatial Domain
Frequency Domain
xz
1/50 nm⁻¹
xy pixel size: 50 nm
z pixel size: 125 nm
Isotropic pixel size (50 nm in x, y, z)
Removing artifacts (DL ground truth)
xz-isotropic resolution
DL Input
b Training model
Convolutional neural network
1D resolution improvement (along x) model
c Resolution recovery at different rotations
0 °
30 °
60 °
90 °
120 °
150 °
Rotated input
DL model
DL output
Rotation back
FFT
1/50 nm⁻¹
d Combination of 6 rotations
Maximum value at each spatial frequency
1/120 nm⁻¹
Inverse FFT

**Supplementary Fig. 17, An improved deep learning method for enhancing axial resolution in 3D SIM data. a**) Training data generation. 3D SIM data are interpolated 2.5-fold along the axial (z) dimension to generate isotropic pixels, slightly blurred in the axial direction to remove spurious sidelobes in the Fourier domain, blurred laterally (along x) to generate data with isotropic resolution, downsampled along x to simulate the acquisition of coarsely acquired axial information, and finally upsampled again to recover an isotropic pixel size. A convolutional neural network **b**) is then trained to recover higher resolution lateral information from the degraded data as indicated. Digitally rotating 3D SIM data about the y axis and passing it through the trained network improves resolution along the lateral coordinate in the rotated space; rotating the data back to the original frame then improves resolution along the direction of rotation **c**). **d**) Recording the maximum value (taken over all rotated outputs in **c**)) at each spatial frequency generates a final prediction with isotropic ~120 nm resolution **d**). All real space scale bars 1 µm; in this example 3D SIM images show Alexa Fluor 488 immunolabeled Tomm20, marking mitochondria, in fixed U2OS cell. See also **Fig. 4a**, **Methods**.

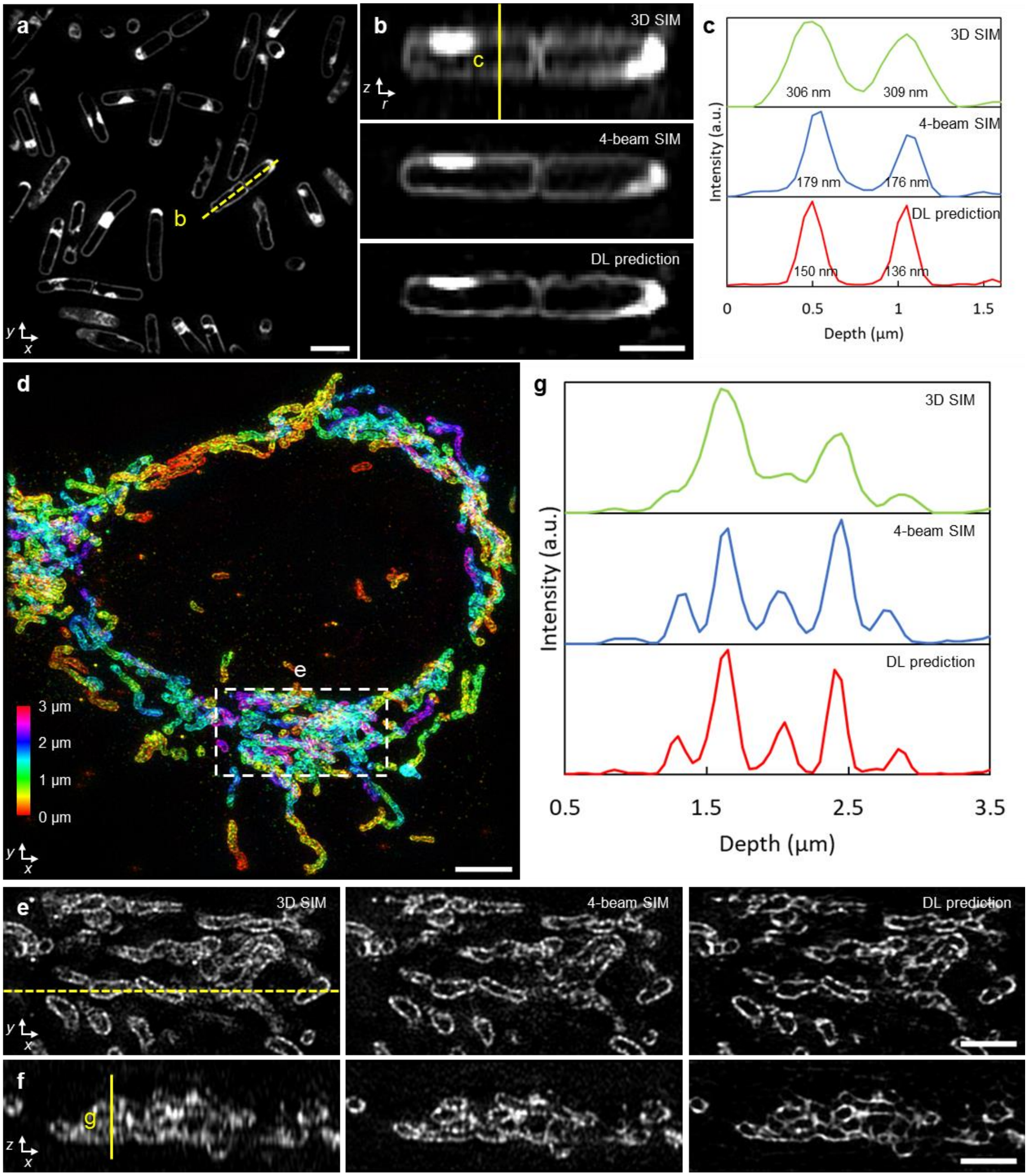


**Supplementary Fig. 18, Images of additional samples, comparing deep learning isotropization output to 3D SIM and 4-beam SIM. a**) DL prediction images of live *B. subtilis* stained for membrane (CellBrite Fix 488). **b**) Axial views along dashed yellow line shown in **a**), comparing 3D SIM (top), 4-beam SIM (middle), and deep learning (DL) prediction based on 3D SIM input. **c**) Line profiles corresponding to yellow vertical line in **b**). Full width at half maximum (FWHM) values under each peak are indicated. **d**) Depth-color coded images of immunolabeled Tomm20 in fixed U2OS cells. Maximum intensity projection of DL prediction is shown. **e**) Higher magnification views of dashed region in **d**), with lateral (top) and corresponding axial views **f**) (corresponding to dashed yellow line in lateral view) comparing 3D SIM (left), 4-beam SIM (middle) and DL prediction (right). **g**) Line profiles along yellow vertical line in **f**). Scale bars: 2 µm **a, e, f**), 1 µm **b**), 4 µm **d**).

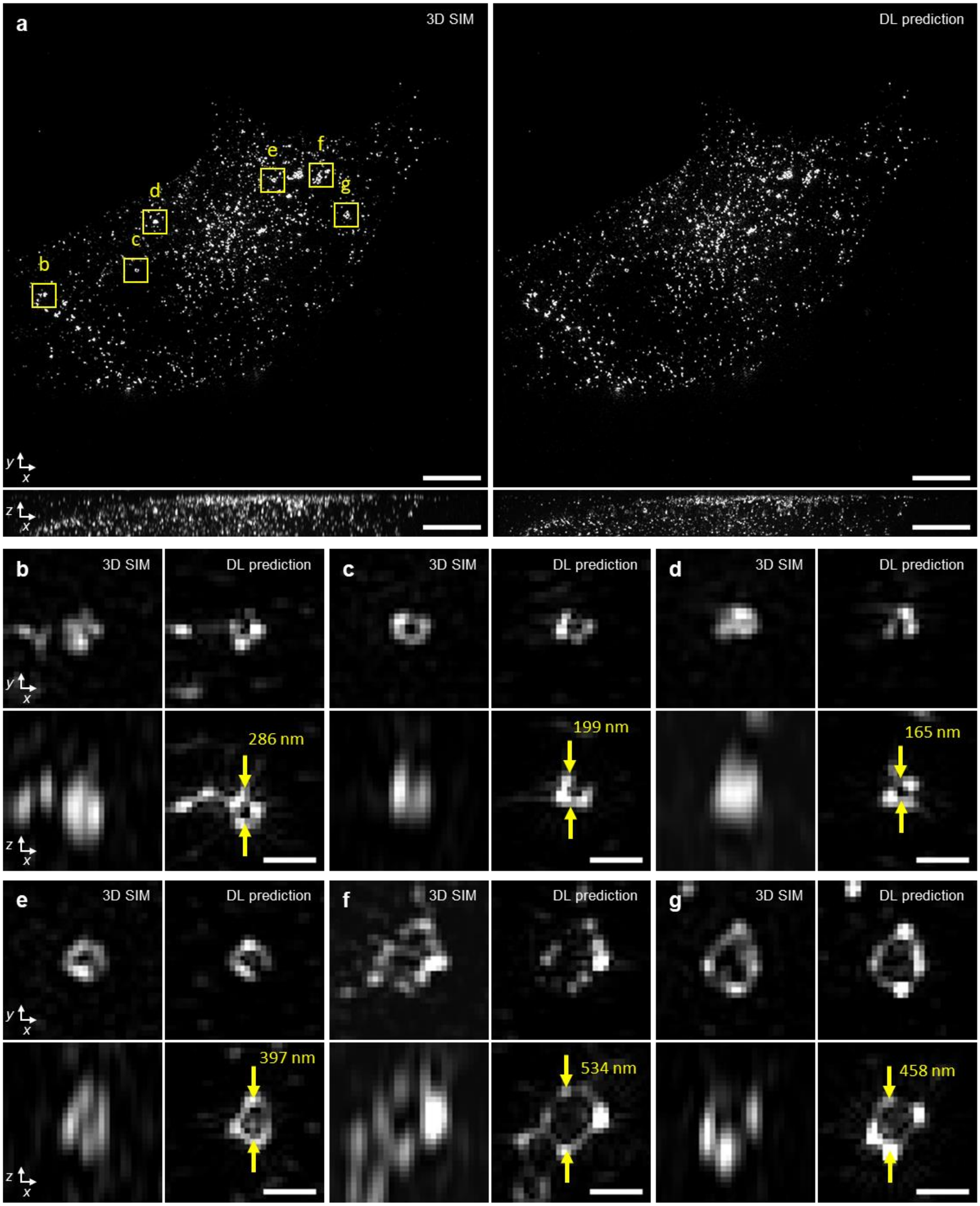


**Supplementary Fig. 19, Caveolin-1-EGFP decorates ring structures of variable diameter in fixed mouse embryonic fibroblasts. a**) 3D SIM images (left) and accompanying deep learning prediction (right). Lateral (top) and axial (bottom) maximum intensity projections are shown. **b-g**) Higher magnification views of yellow rectangular regions in **a**), highlighting ring structures of variable diameter. Apparent ring diameters are marked on axial views. Scale bars: 5 µm **a**); 500 nm **b-f**).

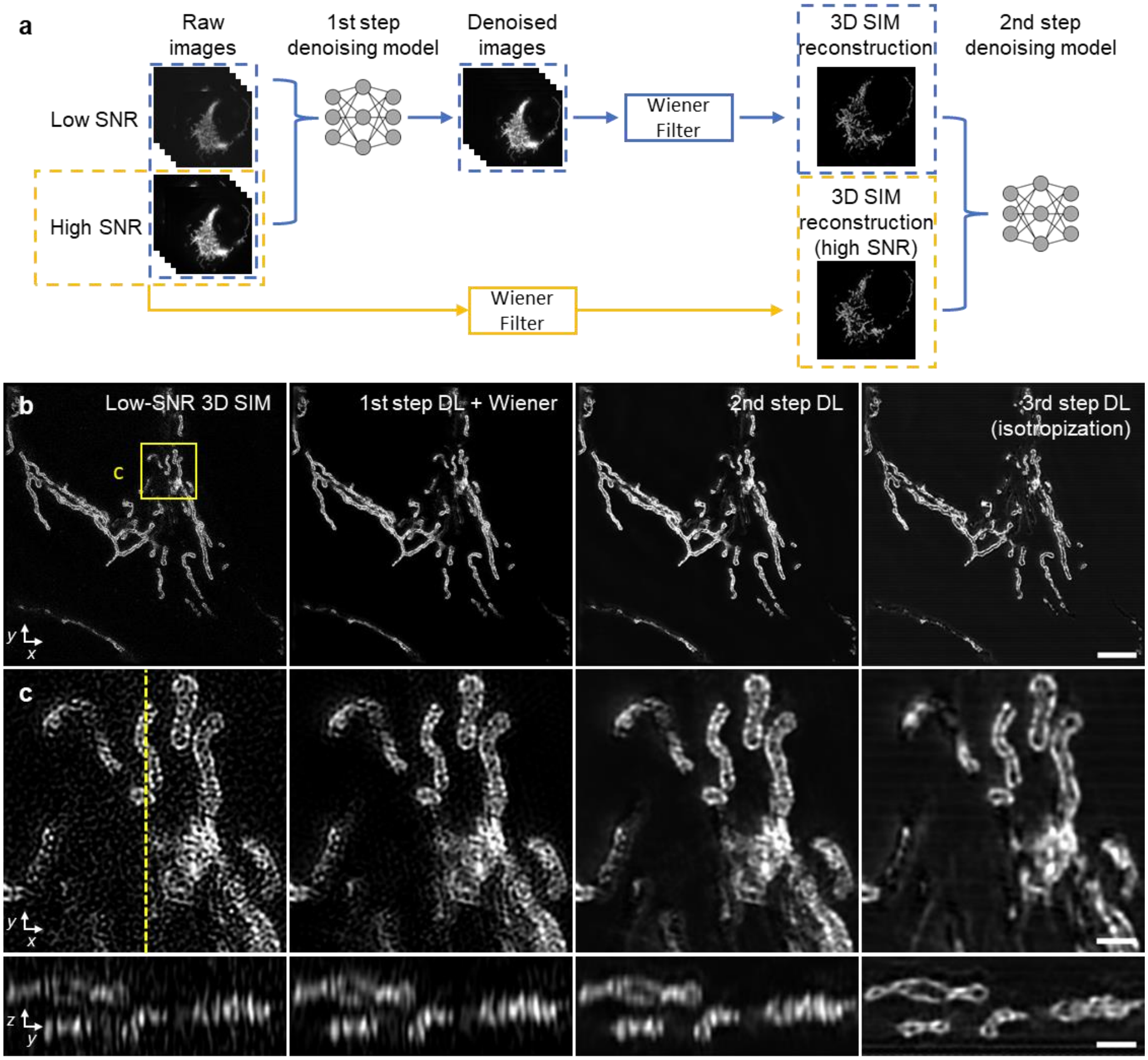


**Supplementary Fig. 20, Multi-step deep learning for denoising and resolution isotropy. a**) Training procedure. Matched sets of raw data (5 phases x 3 orientations) are collected at high and low signal-to-noise ratio (SNR) and are used to train a denoising model (1$^{st}$ step denoising) to denoise low SNR data. A second denoising model (2$^{nd}$ step denoising) is trained using the 3D SIM reconstructions derived from generalized Wiener filters that (1) combine the denoised images from the 1$^{st}$ denoising step or (2) combine the high SNR raw data. The output of this second model may then be fed into a third model that enhances axial resolution, generating a final denoised reconstruction with isotropic resolution. See also **Fig. 5a, Supplementary Figs. 21, 22**. **b**) Comparative images (single planes) of immunolabeled Tomm20 in a fixed U2OS cell, as observed after (left to right): direct 3D SIM reconstruction from the low SNR raw data; after application of the 1$^{st}$ denoising model and Wiener filter; after application of the 2$^{nd}$ denoising model; and after the isotropization model. **c)** *Top*: Higher magnification views of yellow rectangular region in **b**), emphasizing progressive improvement in SNR; *Bottom*: axial view corresponding to yellow dashed line in top view. Scale bars: 5 μm **b**), 1 μm **c**).

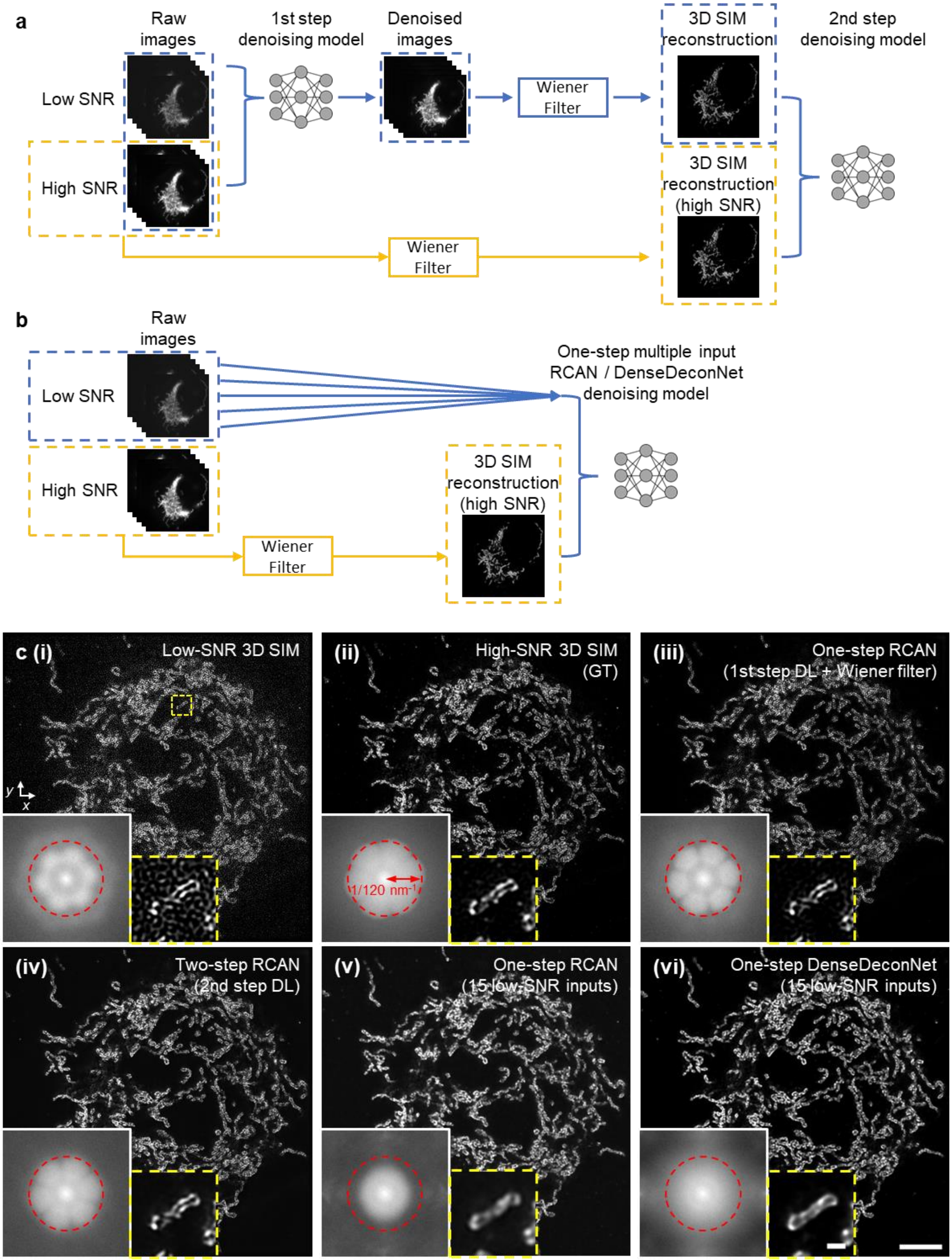

a
Raw images
1st step denoising model
Denoised images
3D SIM reconstruction
2nd step denoising model
Low SNR
High SNR
Wiener Filter
3D SIM reconstruction (high SNR)
Wiener Filter
b
Raw images
One-step multiple input RCAN / DenseDeconNet denoising model
Low SNR
High SNR
3D SIM reconstruction (high SNR)
Wiener Filter
c (i)
Low-SNR 3D SIM
(ii)
High-SNR 3D SIM (GT)
1/120 nm⁻¹
(iii)
One-step RCAN (1st step DL + Wiener filter)
(iv)
Two-step RCAN (2nd step DL)
(v)
One-step RCAN (15 low-SNR inputs)
(vi)
One-step DenseDeconNet (15 low-SNR inputs)

**Supplementary Fig. 21, Comparing different denoising methods. a**) Two step denoising method incorporating Wiener filter, reproduced from **Supplementary Fig. 20a**. This is the method used in the main text, see also **Fig. 5a**. **b**) multiple-input one-step denoising, trained using raw data (5 phases x 3 orientations = 15 inputs) and high SNR 3D SIM ground truth reconstruction. The model is either a multiple input RCAN or a multiple input DenseDeconNet. **c**) Comparative images of immunolabeled Tomm20 in a fixed U2OS cell, as reconstructed i) directly from the raw, input using a generalized Wiener filter; ii) using high SNR input data, i.e., ground truth (GT); iii) one-step RCAN denoising followed by application of the Wiener filter; iv) as in iii), but followed by the 2$^{nd}$ denoising step, i.e., two-step RCAN denoising incorporating the Wiener filter; v) one-step RCAN using 15 low SNR inputs, as shown in **b**); vi) one-step DenseDeconNet using 15 low SNR inputs, as shown in **b**). Insets show Fourier transforms of the images and higher magnification views of mitochondria in dashed yellow box in **c** (**i**). Structural similarity index measure (SSIM) and peak signal-to-noise ratios are also provided in **Supplementary Table 2**, assuming high SNR 3D SIM reconstruction **c** (**ii**) as ground truth. Scale bars: 5 µm, 500 nm (inset).

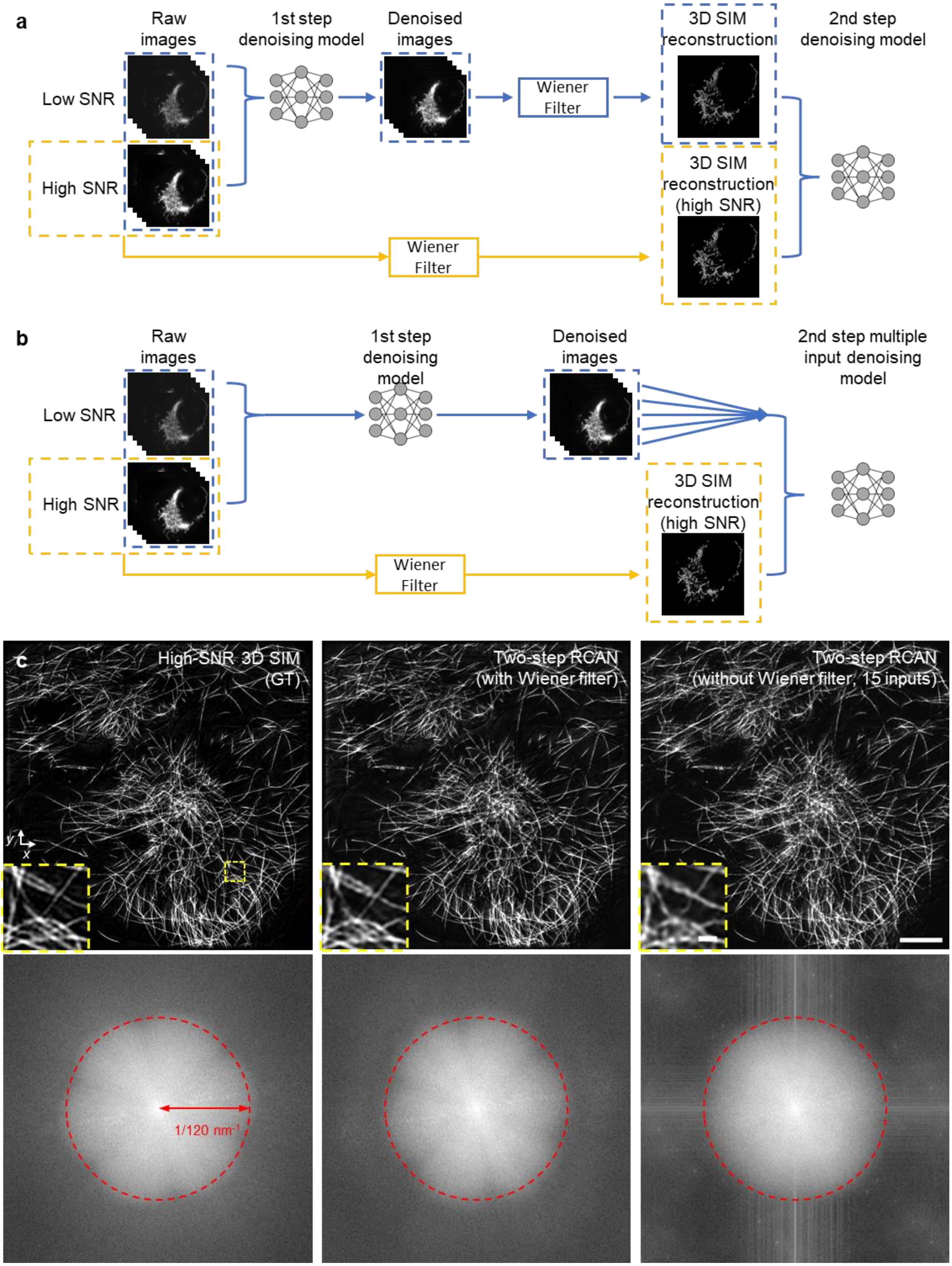
a
Raw images
1st step denoising model
Denoised images
3D SIM reconstruction
2nd step denoising model
Low SNR
High SNR
Wiener Filter
3D SIM reconstruction (high SNR)
Wiener Filter
b
Raw images
1st step denoising model
Denoised images
2nd step multiple input denoising model
Low SNR
High SNR
3D SIM reconstruction (high SNR)
Wiener Filter
c
High-SNR 3D SIM (GT)
Two-step RCAN (with Wiener filter)
Two-step RCAN (without Wiener filter, 15 inputs)
y
x
1/120 nm-1

**Supplementary Fig. 22, Incorporating the Wiener filter between denoising steps improves the performance of two-step denoising. a**) Two step denoising method incorporating Wiener filter, reproduced from **Supplementary Fig. 20a**. This is the method used in the main text, see also **Fig. 5a**. **b**) As in **a**), but without applying the Wiener filter to the output of the first step, instead using the 15 denoised images as input into the 2$^{nd}$ step. **c**) *Top:* Comparative images of immunolabeled microtubules in fixed U2OS cells, as observed after (left to right): performing 3D SIM reconstruction on high SNR raw input, i.e., the ground truth (GT); after applying two-step RCAN with intermediate Wiener filter, i.e., process illustrated in **a**); after applying 2$^{nd}$ RCAN model to set of 15 denoised input images without Wiener filter, i.e., process illustrated in **b**). Insets show higher magnification view of immunolabeled microtubules in yellow dashed rectangular region. *Bottom*: Corresponding Fourier transforms. Note that although Fourier transforms are similar, insets reveal detail lost without Wiener filter. See also the corresponding structural similarity index measure (SSIM) and peak signal-to-noise ratios in **Supplementary Table 3**. Scale bars: 5 µm, 500 nm (inset).

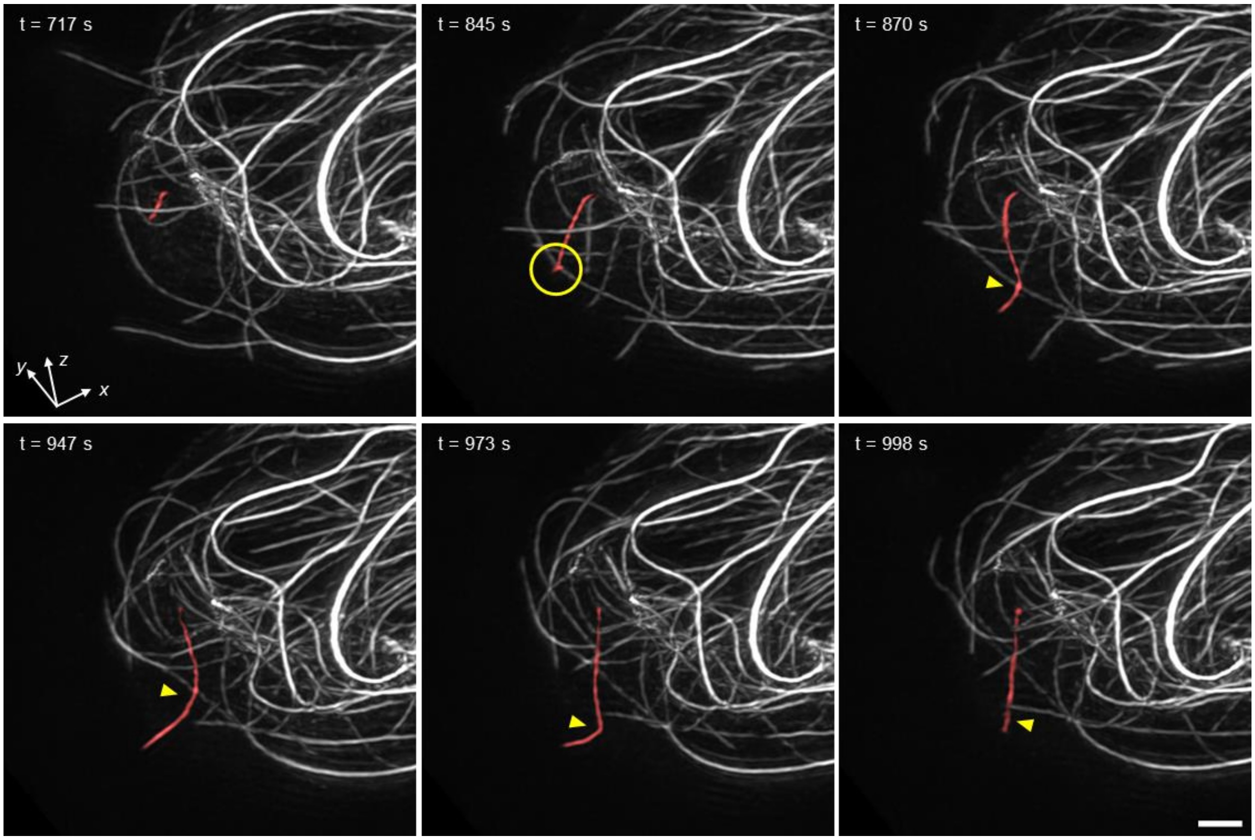


**Supplementary Fig. 23, Local sliding and buckling of microtubules within a living immune cell.** Selected volumetric reconstructions of Jurkat T cell expressing EMTB 3x GFP are shown from 50 time point series (volumes recorded every 25.6 s), in perspective views. Target microtubule (red, t = 717 s) grows and subsequently contacts another microtubule (t = 845 s, position marked with yellow circle). The target microtubule then appears to slide and buckle at the attachment point (t = 870 s, 947 s, yellow arrowhead). A second buckling event occurs (t = 973 s) before the microtubule appears to depolymerize (t= 998 s). See also **Supplementary Video 16**. Scale bars: 2 µm.

**Supplementary Table 1, FWHMs of 100 nm beads acquired with 1.35 NA silicone oil immersion and 1.27 NA water immersion objectives.** N: number of beads used for each statistic, std: standard deviation.

| Figure # | NA | FWHM (nm) | Widefield | | | 3D SIM | | | 4-beam SIM | | |
|---|---|---|---|---|---|---|---|---|---|---|---|
| Fig. 1g | 1.35 | | Mean | std | N | Mean | std | N | Mean | std | N |
| | | Lateral | 267.5 | 15.7 | 102 | 118.6 | 10.9 | 100 | 123.5 | 12.1 | 99 |
| | | Axial | 581.4 | 22.5 | | 300.6 | 12.9 | | 163.0 | 12.9 | |
| Supplementary Fig. 10d | 1.27 | | Mean | std | N | Mean | std | N | Mean | std | N |
| | | Lateral | 260.4 | 16.4 | 85 | 120.4 | 11.5 | 81 | 127.6 | 9.8 | 78 |
| | | Axial | 595.1 | 22.2 | | 350.3 | 15.1 | | 164.4 | 25.0 | |

**Supplementary Table 2, Quantitative comparisons amongst denoising methods, as assayed on images of immunolabeled Tomm20 in fixed U2OS cells.** 3D SIM reconstructions derived from high SNR input were taken as the ground truth. std: standard deviation. PSNR values are reported in dB. N: number of 2D slices used for each statistic, std: standard deviation. See also **Supplementary Fig. 21**. Entries with highest (best) values are bolded.

| **Figure #** | **Methods** | **SSIM** | | | **PSNR** | | |
|---|---|---|---|---|---|---|---|
| Supplementary Fig. 21c | | Mean | std | N | Mean | std | N |
| | **Low-SNR 3D SIM** | 0.71 | 0.11 | 11 | 30.51 | 2.97 | 11 |
| | **One-step RCAN (1st step DL + Wiener filter)** | 0.83 | 0.06 | 11 | 32.78 | 3.29 | 11 |
| | **Two-step RCAN (2nd step DL)** | **0.86** | 0.03 | 11 | **34.14** | 3.18 | 11 |
| | **One-step RCAN (15 low-SNR inputs)** | 0.81 | 0.04 | 11 | 32.75 | 3.24 | 11 |
| | **One-step DenseDeconNet (15 low-SNR inputs)** | 0.85 | 0.04 | 11 | 33.14 | 3.26 | 11 |

**Supplementary Table 3, Quantitative comparisons of two-step RCAN with and without intermediate Wiener filter, as assayed on images of immunolabeled microtubules in fixed U2OS cells.** 3D SIM reconstructions derived from high SNR input were taken as the ground truth. std: standard deviation. PSNR values are reported in dB. See also **Supplementary Fig. 22.** N: number of 2D slices used for each statistic, std: standard deviation. Entries with highest values are bolded.

| **Figure #** | **Methods** | **SSIM** | | | **PSNR** | | |
|---|---|---|---|---|---|---|---|
| Supplementary Fig. 22c | | Mean | std | N | Mean | std | N |
| | **Two-step RCAN (with Wiener filter)** | **0.84** | 0.03 | 14 | **33.09** | 1.63 | 14 |
| | **Two-step RCAN (without Wiener filter)** | 0.78 | 0.04 | 14 | 31.45 | 1.41 | 14 |

**Supplementary Table 4, Data acquisition and processing parameters for data acquired with the SIM platform.** All raw images have lateral dimensions of 1280 x 1080 pixels and are cropped into square regions for further processing. The volume size W x H x D represent voxel size in x, y, and z dimensions, C is the number of colors. The volume acquisition time includes all colors and 15 phases. In the "Deep learning?" column, A means axial resolution improvement and DN means denoising.

| Figure # | Sample/structure (excitation wavelength) | Objective NA | Imaging mode | Laser Intensity (W/cm$^2$) | volume size W x H x D x C (z step size) | Volume acquisition time | Volume or subvolume # (time interval) | Volume size after reconstruction (sample size, μm$^3$) | Deep learning ? |
|---|---|---|---|---|---|---|---|---|---|
| Fig. 1d, e | 100 nm yellow-green beads (488 nm) | 1.35 | 4-beam SIM | 15 | 512 x 512 x 67 x 1 (60 nm) | 26.5 s | 1 | 1024 x 1024 x 67 (~36 x 36 x 4) | No |
| | | | 3D SIM | 15 | 512 x 512 x 32 x 1 (125 nm) | 12.7 s | 1 | 1024 x 1024 x 32 (~36 x 36 x 4) | |
| Supplementary Fig. 8 | 100 nm yellow-green beads (488 nm) | 1.27 | 3D SIM | 5 | 256 x 256 x 32 x 1 (125 nm) | 12.7 s | 1 | 512 x 512 x 32 (~21 x 21 x 4) | No |
| Supplementary Fig. 10 | 100 nm yellow-green beads (488 nm) | 1.27 | 4-beam SIM | 15 | 512 x 512 x 67 x 1 (60 nm) | 26.5 s | 1 | 1024 x 1024 x 67 (~42 x 42 x 4) | No |
| | | | 3D SIM | 15 | 512 x 512 x 32 x 1 (125 nm) | 12.7 s | 1 | 1024 x 1024 x 32 (~42 x 42 x 4) | |
| Fig. 2a-c Supplementary Video 2 | Live vegetative *B. subtilis*, membranes (488 nm) | 1.35 | 4-beam SIM | 4 | 256 x 256 x 67 x 1 (60 nm) | 26.5 s | 1 | 512 x 512 x 67 (~18 x 18 x 4) | No |
| | | | 3D SIM | 15 | 256 x 256 x 32 x 1 (125 nm) | 12.7 s | 1 | 512 x 512 x 32 (~18 x 18 x 4) | |
| Fig. 2e-g Supplementary Video 3 | Fixed U2OS cell, outer mitochondrial membranes (488 nm) | 1.35 | 4-beam SIM | 4 | 512 x 512 x 67 x 1 (60 nm) | 26.5 s | 1 | 1024 x 1024 x 67 (~36 x 36 x 4) | No |
| | | | 3D SIM | 15 | 512 x 512 x 32 x 1 (125 nm) | 12.7 s | 1 | 1024 x 1024 x 32 (~36 x 36 x 4) | |

| | | | | | | | | | |
|---|---|---|---|---|---|---|---|---|---|
| Fig. 2h, i Supplementary Video 4 | Live U2OS cell, inner mitochondrial membranes (488 nm) | 1.35 | 4-beam SIM | 5 | 512 x 512 x 67 x 1 (60 nm) | 26.5 s | 1 | 1024 x 1024 x 32 (~36 x 36 x 4) | No |
| Supplementary Fig. 13b | Fixed U2Os cell, F-actins (488 nm) | 1.35 | 3D SIM | 4 | 512 x 512 x 64 x 1 (125 nm) | 25.3 s | 1 | 1024 x 1024 x 64 (~36 x 36 x 8) | No |
| Fig. 3a, b | Live, sporulating *B. subtilis*, spores (488 nm) and membranes (561 nm) | 1.27 | 4-beam SIM | 25 (488 nm)<br>13 (561 nm) | 256 x 256 x 67 x 2 (60 nm) | 53 s | 1 | 512 x 512 x 67 (~21 x 21 x 4) | No |
| Fig. 3c-g | Fixed U2OS cell, microtubules (488 nm) and vimentin (561 nm) | 1.27 | 4-beam SIM | 20 (488 nm)<br>15 (561 nm) | 960 x 960 x 100 x 2 (60 nm) | 79 s | 1 | 1920 x 1920 x 100 (~78 x 78 x 6) | No |
| Fig. 3h-n Supplementary Video 6 | Fixed mouse liver sinusoidal endothelial cell, membranes (488 nm) and actins (561 nm) | 1.27 | 4-beam SIM | 4.5 (488 nm)<br>25 (561 nm) | 1024 x 1024 x 67 x 2 (60 nm) | 53 s | 1 | 2048 x 2048 x 67 (~84 x 84 x 4) | No |
| Supplementary Fig. 14a, b Supplementary Video 5 | Live, vegetative *B. subtilis*, DivIVA-GFP (488 nm) | 1.27 | 4-beam SIM | 25 | 352 x 352 x 67 x 1 (60 nm) | 26.5 s | 1 | 704 x 704 x 67 (~29 x 291 x 4) | No |
| Supplementary Fig. 14c-d | Fixed U2OS cell, microtubules (488 nm) | 1.27 | 4-beam SIM | 15 | 768 x 768 x 100 x 1 (60 nm) | 39.5 | 1 | 1536 x 1536 x 100 (~63 x 63 x 6) | No |
| | | | 3D SIM | 25 | 768 x 768 x 48 x 1 (125 nm) | 19 s | 1 | 1536 x 1536 x 48 (~63 x 63 x 6) | |
| Supplementary Fig. 14g, h Supplementary Video 7 | Live U2OS cell, membranes (561 nm) | 1.27 | 4-beam SIM | 25 | 512 x 512 x 67 x 1 (60 nm) | 26.5 s | 10 (19 s) | 1024 x 1024 x 67 (~42 x 42 x 4) | No |

| | | | | | | | | | |
|---|---|---|---|---|---|---|---|---|---|
| Supplementary Fig. 16c | Fixed U2OS cell, microtubules (488 nm) | 1.27 | 3D SIM | 4.5 | 512 x 512 x 48 x 1 (125 nm) | 19 s | 1 | 1024 x 1024 x 48 (~42 x 42 x 6) | Yes (A) |
| Supplementary Fig. 16d | Fixed U2OS cell, outer mitochondrial membranes (488 nm) | 1.35 | 3D SIM | 5 | 512 x 512 x 64 x 1 (125 nm) | 25.3 s | 1 | 1024 x 1024 x 64 (~36 x 36 x 8) | Yes (A) |
| Supplementary Fig. 16e-h | Live U2OS cell, inner mitochondrial membranes (488 nm) | 1.35 | 3D SIM | 5 | 512 x 512 x 64 x 1 (125 nm) | 25.3 s | 1 | 1024 x 1024 x 64 (~36 x 36 x 8) | Yes (A) |
| Fig. 4b, c | Fixed U2OS cell, microtubules (488 nm) | 1.27 | 4-beam SIM | 15 | 832 x 832 x 100 x 1 (60 nm) | 39.5 s | 1 | 1664 x 1664 x 100 (~68 x 68 x 6) | Yes (A) |
| | | | 3D SIM | 25 | 832 x 832 x 48 x 1 (125 nm) | 19 s | 1 | 1664 x 1664 x 48 (~68 x 68 x 6) | |
| Fig. 4e-k | Fixed mouse embryonic fibroblast, Caveolin-1 (488 nm) and Cavin-1 (561 nm) | 1.27 | 3D SIM | 25 (488 nm) 21 (561 nm) | 512 x 512 x 32 (125 nm) | 54.5 s | 1 | 1024 x 1024 x 32 (~42 x 42 x 4) | Yes (A) |
| Supplementary Fig. 18a, b | Live vegetative *B. subtilis*, membranes (488 nm) | 1.35 | 4-beam SIM | 4 | 256 x 256 x 67 x 1 (60 nm) | 26.5 s | 1 | 512 x 512 x 67 (~18 x 18 x 4) | Yes (A) |
| | | | 3D SIM | 15 | 256 x 256 x 32 x 1 (125 nm) | 12.7 s | 1 | 512 x 512 x 32 (~18 x 18 x 4) | |
| Supplementary Fig. 18d-f | Fixed U2OS cell, outer mitochondrial membranes (488 nm) | 1.35 | 4-beam SIM | 4 | 512 x 512 x 67 x 1 (60 nm) | 26.5 s | 1 | 1024 x 1024 x 67 (~36 x 36 x 4) | Yes (A) |
| | | | 3D SIM | 15 | 512 x 512 x 32 x 1 (125 nm) | 12.7 s | 1 | 1024 x 1024 x 32 (~36 x 36 x 4) | |
| Supplementary Fig. 19 | Fixed mouse embryonic fibroblast, Caveolin-1 (488 nm) | 1.27 | 3D SIM | 25 | 512 x 512 x 32 (125 nm) | 27.1 s | 1 | 1024 x 1024 x 32 (~42 x 42 x 4) | Yes (A) |

| | | | | | | | | | |
|---|---|---|---|---|---|---|---|---|---|
| Fig. 5b-d Supplementary Video 8 | Live U2OS cell, outer mitochondrial membranes (488 nm) | 1.27 | 3D SIM | 0.5 | 512 x 512 x 32 x 1 (125 nm) | 12.7 s | 50 (9.7 s) | 1024 x 1024 x 32 (~42 x 42 x 4) | Yes (DN + A) |
| Fig. 5e-k Supplementary Video 10, 12 | Live U2OS cell, lysosomal membranes (488 nm) and interior of lysosomes (561 nm) | 1.27 | 3D SIM | 0.8 (488 nm) 0.3 (561 nm) | 512 x 512 x 32 x 2 (125 nm) | 25.8 s | 60 | 1024 x 1024 x 32 (~42 x 42 x 4) | Yes (DN + A) |
| Supplementary Video 9 | Live U2OS cell, lysosomal membranes (488 nm) | 1.27 | 3D SIM | 2 | 576 x 576 x 32 x 1 (125 nm) | 12.7 s | 50 (4.8 s) | 1152 x 1152 x 32 (~47 x 47 x 4) | Yes (DN + A) |
| Supplementary Video 11 | Live U2OS cell, lysosomal membranes (488 nm) and interior of lysosomes (561 nm) | 1.27 | 3D SIM | 0.5 (488 nm) 0.3 (561 nm) | 448 x 448 x 24 x 2 (125 nm) | 19 s | 60 | 896 x 896 x 24 (~37 x 37 x 3) | Yes (DN + A) |
| Supplementary Fig. 20b, c | Fixed U2OS cell, outer mitochondrial membranes (488 nm) | 1.35 | 3D SIM | 0.8 | 512 x 512 x 32 x 1 (125 nm) | 12.7 s | 1 | 1024 x 1024 x 32 (~36 x 36 x 4) | Yes (DN + A) |
| Supplementary Fig. 21 c | Fixed U2OS cell, outer mitochondrial membranes (488 nm) | 1.35 | 3D SIM | 0.8 | 512 x 512 x 64 x 1 (125 nm) | 25.3 s | 1 | 1024 x 1024 x 64 (~36 x 36 x 8) | Yes (DN) |
| Supplementary Fig. 22 c | Fixed U2OS cell, microtubules (488 nm) | 1.35 | 3D SIM | 0.4 | 512 x 512 x 48 x 1 (125 nm) | 19 s | 1 | 1024 x 1024 x 48 (~36 x 36 x 6) | Yes (DN) |
| Fig. 6 Supplementary Video 13-15 | Live Jurkat T cell, microtubules (488 nm) | 1.27 | 3D SIM | 0.8 | 256 x 256 x 32 x 1 (125 nm) | 12.8 s | 100 | 512 x 512 x 32 (~21 x 21 x 4) | Yes (DN + A) |

| | | | | | | | | | |
|---|---|---|---|---|---|---|---|---|---|
| Supplementary Fig. 23 Supplementary Video 16 | Live Jurkat T cell, microtubules (488 nm) | 1.27 | 3D SIM | 0.5 | 256 x 256 x 32 x 1 (125 nm) | 12.8 s | 50 (12.8 s) | 512 x 512 x 32 (~21 x 21 x 4) | Yes (DN + A) |

**Legends for Supplementary Videos**

**Supplementary Video 1**: Images of autofluorescence from dirty coverslip during z stack with lateral illumination modulation (left, before mirror alignment) vs. mostly axial modulation (right, after mirror alignment). See also **Supplementary Fig. 9c**.

**Supplementary Video 2**: 3D projections of live vegetative *B. subtilis* stained with CellBrite Fix 488, marking membranes. Widefield (top), 3D-SIM (middle), and 4-beam SIM reconstructions (bottom) are compared. See also **Fig. 2b**.

**Supplementary Video 3**: 3D projections of fixed U2OS cell labeled with Tomm20 primary and rabbit-Alexa Fluor 488 secondary antibodies, marking outer mitochondrial membranes. Widefield (left), 3D-SIM (middle), and 4-beam SIM (right) reconstructions are compared. See also **Fig. 2f, g**.

**Supplementary Video 4**: Live U2OS cell stained with MitoTracker Green FM. First movie segment shows lateral views as a function of z (i.e., 'z stack'). Second movie segment shows the axial view as a function of lateral coordinate. See also **Fig. 2h, i**.

**Supplementary Video 5**: 4-beam SIM imaging of live, vegetative *B. subtilis* with GFP-DivIVA marker, projection view. See also **Supplementary Fig. 14b**.

**Supplementary Video 6**: 4-beam SIM imaging of fixed mouse liver sinusoidal endothelial cell with CellBrite Fix 488 label, marking membrane (green) and Alexa Fluor 568 Phalloidin, marking actin filaments (red). See also **Fig. 3h**.

**Supplementary Video 7**: Time-lapse imaging of live U2OS cell stained with 200 nM Potomac Gold in 4-beam SIM. Yellow arrow and rectangle highlight phototoxicity in one region. Red arrows highlight morphological changes of mitochondria between the 1$^{st}$ time point and the 10$^{th}$ time point. See also **Supplementary Fig. 14g, h**.

**Supplementary Video 8**: Time-lapse imaging of live U2OS cell expressing Tomm20-GFP marker. First movie segment shows 3D projections of mitochondrial dynamics after two-step denoising and isotropic prediction. Second movie segment shows higher magnification views of yellow rectangular region in the first movie segment. Raw low-SNR 3D SIM reconstructions (top left), 1$^{st}$ step denoising results (top right), 2$^{nd}$ step denoising results (bottom left) and 3$^{rd}$ step isotropization (bottom right) are compared. Third movie segment shows axial planes indicated by yellow lines in the second movie segment. See also **Fig. 5b-d**.

**Supplementary Video 9**: Time-lapse imaging of live U2OS cell expressing LAMP1-EGFP marker. First movie segment shows 3D projections of lysosomal dynamics after two-step denoising and isotropic prediction. Second movie segment shows higher magnification views of yellow rectangular region in the first movie segment. Raw low-SNR 3D SIM reconstructions (top left), 1$^{st}$ step denoising results (top right), 2$^{nd}$ step denoising results (bottom left) and 3$^{rd}$ step isotropization (bottom right) are compared. Third movie segment shows axial planes indicated by yellow lines in the second movie segment.

**Supplementary Video 10**: Time-lapse imaging of live U2OS cell expressing lysosomal marker LAMP1-GFP (green) and additionally labeled with LysoTracker Red to mark the lysosome interior (red). 3D projections of raw low-SNR 3D SIM reconstructions (left) and two-step denoising and isotropization predictions (right) are compared. See also **Fig. 5e**.

**Supplementary Video 11**: Time-lapse imaging of additional live U2OS cell #2 expressing lysosomal marker LAMP1-GFP (green) and additionally labeled with LysoTracker Red to mark the lysosome interior (red). 3D projections of raw low-SNR 3D SIM reconstructions (left) and two-step denoising and isotropization predictions (right) are compared.

**Supplementary Video 12**: Higher magnification view of maximum intensity projections of live U2OS cell in **Supplementary Video 10**. See also **Fig. 5k**.

**Supplementary Video 13**: Time-lapse imaging of live Jurkat T cell expressing EMTB 3x GFP after two-step denoising and isotropization prediction, perspective view. Two microtubule filaments are segmented (red, green), emphasizing that they are initially separated (yellow circles), merge together over the nucleus (yellow arrow), and separate again (yellow circles). See also **Fig. 6e**.

**Supplementary Video 14**: Time-lapse imaging of live Jurkat T cell expressing EMTB 3x GFP after two-step denoising and isotropization prediction. Centrosome is identified (cyan sphere, trajectory temporally coded as shown in color bar) and one microtubule filament is segmented (red), emphasizing their correlated, inward movement. Top: lateral perspective view. Bottom: Axial yz view. See also **Fig. 6f**.

**Supplementary Video 15**: Time-lapse imaging of live Jurkat T cell expressing EMTB 3x GFP after two-step denoising and isotropization prediction, emphasizing buckling of two microtubule filaments (red and yellow spheres). Top: lateral perspective view. Bottom: Axial yz view. See also **Fig. 6g**.

**Supplementary Video 16**: Time-lapse imaging of another live Jurkat T cell expressing EMTB 3x GFP after two-step denoising and isotropization prediction. One microtubule filament is segmented (red) and yellow arrows indicate buckling sites after sliding and contacting another filament. Left: lateral view. Right: zoomed-in perspective view. See also **Supplementary Fig. 23**.
